# Functional and phylogenetic analysis of placozoan GPCRs reveal the prebilaterian origin of monoaminergic signalling

**DOI:** 10.1101/2025.04.18.649542

**Authors:** Luis Alfonso Yañez-Guerra, Tatiana D. Mayorova, Li Guan, Gáspár Jékely, Herman Wijnen, Adriano Senatore

## Abstract

Monoamines are biologically active compounds crucial for neurotransmission and various physiological processes. They include neurotransmitters like serotonin, dopamine, and melatonin, which regulate mood, movement, and sleep in humans. In ecdysozoans, monoamines such as tyramine are important for modulating locomotion, learning, and feeding. The monoaminergic signalling system has been considered a bilaterian innovation, with conflicting evidence supporting its existence in earlier branching, non-bilaterian animals. Here, we challenge the bilaterian origin hypothesis by combining large-scale receptor deorphanisation with phylogenetic analyses to identify monoamine receptors from the placozoan *Trichoplax adhaerens*. We demonstrate that these receptors are homologous to known bilaterian GPCRs, and behavioural assays demonstrate that monoamines like tyramine and tryptamine affect the speed of locomotion and body shape of this animal, respectively. These responses, together with the presence of biosynthetic enzymes for these molecules, reveal that monoaminergic signalling is both active and endogenous in placozoans. Our findings provide compelling evidence for a prebilaterian origin of monoaminergic systems, reshaping our understanding of early nervous system evolution.

## Introduction

Monoamines are a class of biologically active compounds that play crucial roles in neurotransmission and a wide range of physiological processes across the animal kingdom. These molecules, which include well-known neurotransmitters such as serotonin (5-HT), histamine, melatonin, and lesser-known “trace-amines” like tyramine (TA), tryptamine (TPA), and octopamine (p-OCT), are derived from aromatic amino acids (Daubner, Le, and Wang 2011; Horie et al. 2009; M. Zhang 2016; Yamamoto and Vernier 2011). In bilaterian animals, monoamines regulate mood, movement, and other key physiological functions. For instance, serotonin modulates mood, appetite, and sleep in humans. Dopamine is critical for reward-motivated behaviour, learning, and motor control, and accordingly, dysregulated dopamine signalling is a primary cause of Parkinson’s disease (Naoi, Maruyama, and Shamoto-Nagai 2018; Meng et al. 2018; Bacqué-Cazenave et al. 2020; Eiden and Weihe 2011). Another monoamine, melatonin, is widely recognized for its pivotal role in regulating the sleep-wake cycle by synchronizing circadian rhythms and facilitating sleep initiation and maintenance (Claustrat and Leston 2015; Zisapel 2001). In insects, monoamines also play essential roles in physiology and behaviour. Octopamine enhances locomotor activity and is crucial for the “fight or flight” response, dependent on both octopamine and its precursor, tyramine (Pauls et al. 2018; Ma et al. 2015; Sinakevitch et al. 2018; Fussnecker, Smith, and Mustard 2006). Tyramine can regulate chloride permeability in the Malpighian tubules of *Drosophila*, indicating its involvement in osmoregulation and ion transport (Blumenthal 2003).

While monoamines are clearly important in regulating behaviour in bilaterian animals, the roles of these molecules in non-bilaterians remain unclear. Non-bilaterians, lack the enzymes tyrosine hydroxylase (TH) and tryptophan hydroxylase (TPH), which are rate-limiting enzymes for the synthesis of serotonin, dopamine, and other monoamines (Francis et al. 2017; Jin et al. 2024; Goulty et al. 2023). However, non-bilaterians do possess aromatic amino acid decarboxylases (AADC) (Francis et al. 2017; Moroz, Romanova, and Kohn 2021; Jin et al. 2024), which, as the name indicates, can decarboxylase amino acids to generate monoamines like tyramine (from tyrosine), phenethylamine (from phenylalanine), and tryptamine (from tryptophan) (Zhu et al. 2020; Speight et al. 2000; H. Zhang et al. 2019; M. D. Berry et al. 1996). Furthermore, a recent study demonstrated that larvae of the sponge species *Amphimedon queenslandica* show behavioural responses to exogenously applied monoamines like tyramine and phenethylamine (Xiang et al. 2023). As such, *A. queenslandica* and perhaps other species in the phylum Porifera may possess receptors for detecting trace monoamines. More recently, epinephrine-like molecules were found to influence locomotion and feeding behaviour of the non-bilaterian *Trichoplax adhaerens* (phylum Placozoa), a small marine invertebrate that is motile but lacks synapses and a nervous system. This study identified a putative receptor via RNA knockdown, where its downregulation diminished behavioural responses to applied epinephrine and related molecules (Jin et al., 2024).

Despite this growing evidence of monoamine signalling in non-bilaterians, previous studies, have argued that monoamine signalling is a bilaterian innovation (Goulty et al. 2023). This claim was based on the apparent absence of certain biosynthetic enzymes, vesicular transporters and receptors in non-bilaterians. Moreover, it was suggested that non-bilaterians lack a complete monoamine biosynthetic pathway, or that they may be using alternative non-canonical pathways for the synthesis of monoamines (Goulty et al. 2023). However, as noted above, AADC alone is sufficient to produce simple monoamines like tyramine (TA), tryptamine (TPA), and phenethylamine (PEA) (Zhu et al. 2020; Speight et al. 2000; H. Zhang et al. 2019; M. D. Berry et al. 1996). This contradiction raises an important question: if non-bilaterians can synthesize monoamines and responding to them behaviourally—as observed in sponges and placozoans—then monoaminergic receptors may indeed be present, suggesting that monoamine signalling has deeper evolutionary roots than previously assumed.

To address this, we investigated the presence of monoamine receptors in *Trichoplax adhaerens*, the most intensively studied species in the phylum Placozoa. By performing comprehensive phylogenies combined with large-scale combinatorial deorphanisation assays, we identified five *T. adhaerens* G-protein coupled receptors (GPCRs) that are highly sensitive to monoamines. Phylogenetic analyses revealed two lineages of placozoan receptors that are sensitive to monoamines, forming distinct clades with bilaterian melatonergic GPCRs and a set of orphan receptors. For comparison, we assayed ligand activation of the two human melatonin receptors, MelA and MelB, finding that in addition to their namesake ligand, these receptors elicited strong responses to N-acetyltryptamine, N-acetyltyramine, and N-acetylserotonin, while tryptamine caused weak activation of MelB specifically. This contrasted the placozoan receptors which were activated by tryptamine, tyramine, serotonin, and phenethylamine. Structurally, the placozoan receptors appear to utilize fundamentally different molecular determinants for binding monoamines, lacking core residues that mediate ligand binding in the solved structures of the two human melatonin receptors (Stauch et al. 2019; Wang et al. 2022; Patel et al. 2020). Finally, *in vivo* testing of these compounds in *T. adhaerens* revealed that these molecules affect behaviour and cellular responses at concentrations similar to those that activate the receptors *in vitro*. Altogether, our work supports a pre-bilaterian origin for monoamine signalling systems.

## Results

### Large-scale combinatorial deorphanisation analyses reveal the existence of functional monoamine receptors in *Trichoplax adhaerens*

We conducted a comprehensive search for monoamine GPCRs and large-scale combinatorial deorphanisations in the species *Trichoplax adhaerens*. To identify monoamine receptors, we generated a profile hidden Markov model (HMM) trained on sequences of functionally characterised monoamine receptors from different species, including, humans, *Drosophila melanogaster*, *Caenorhabditis elegans*, and *Platynereis dumerilii* (Yoshida et al. 2014; Nakagawa et al. 2022; Bayliss, Roselli, and Evans 2013; Branicky and Schafer 2009; Bauknecht and Jékely 2017; Hauser et al. 2006; Cazzamali, Klaerke, and Grimmelikhuijzen 2005). This model was used to search for candidate homologues in genomes of seven cnidarian, seven bilaterian, four placozoan, four ctenophores, five sponge, and four single-celled eukaryote species. Due to the similarities among Family A (Rhodopsin-like) receptors, the model also detected opsins and other neurotransmitter receptors, such as purinergic and neuropeptide receptors retrieving a total of 3,166 hits. To selectively identify candidate monoaminergic receptors, these sequences were clustered and analysed with the all-vs-all clustering algorithm CLANS (P-value of 1e−28), revealing two primary clusters with varying conservation across major metazoan lineages: one dominated by monoaminergic-related receptors, including opsins, adenosine, and cholinergic receptors; and another dominated by neuropeptide and related receptors (*e.g.*, chemokine, P2Y purinergic) (Fig. 1). We identified receptor sequences from cnidarians and placozoans linking to both major receptor clusters. Ctenophores connected only to the monoamine receptor cluster, while poriferans showed no detectable connections at this threshold (Fig. 1).

**Figure 1.**
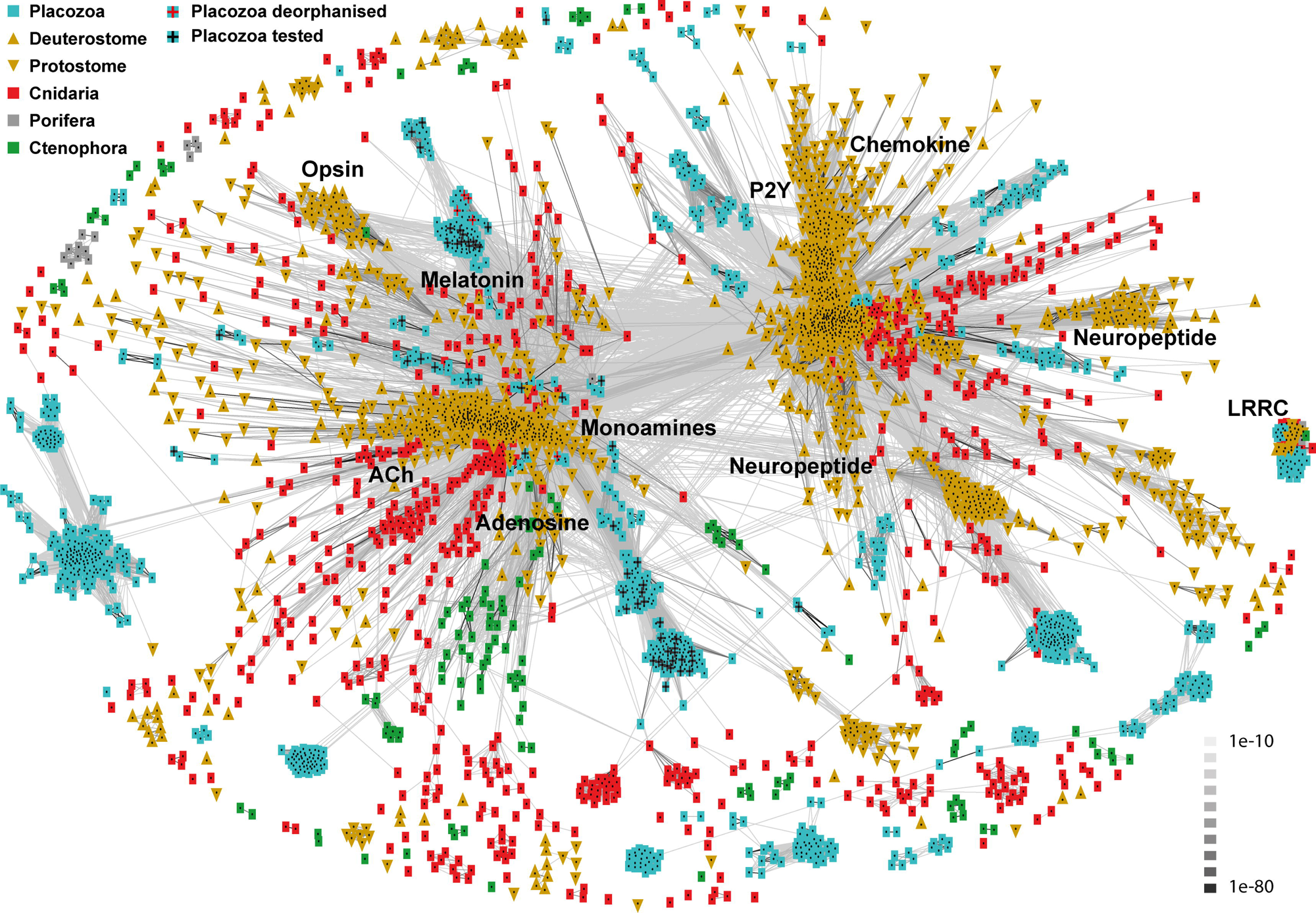
CLANS analysis of monoamine metazoan receptors. Cluster analysis of major class A GPCR groups from metazoans was obtained using an *ad hoc* monoamine hidden Markov model based on deorphanized receptors from bilaterians, with a minimum connection cutoff P-value of 1e-28. Each dot represents a GPCR sequence, colour-coded and symbolised according to the key in the top left. Connecting lines between sequences indicate similarity, with P-values noted in the bottom right. Cluster annotations are based on known deorphanised bilaterian class-A GPCRs. The teal squares with black crosses indicate the receptors tested and red crosses indicate receptors deorphanised in *Trichoplax adhaerens*.

Next, we sought to functionally characterise candidate monoamine receptors from placozoans. We synthesized and cloned the complimentary DNAs (cDNAs) of 56 receptors from *T. adhaerens* based on their association with the monoamine cluster and the presence of 7 predicted transmembrane (TM) helices (*i.e.*, partial receptor sequences were omitted). The list of receptors chosen is available in Supplementary File 1 and the nucleotide sequences of cloned/synthesised receptors are provided in Supplementary File 2. Using a large-scale combinatorial deorphanisation approach, as described previously (Thiel et al. 2024), we expressed these receptors in HEK-293 cells and tested them with five mixtures of monoamines, related compounds and polyamines. Mixture 1 contained; Phenethylamine, melatonin, N-acetylserotonin (N-acetyl-5HT), N-acetyltyramine (N-acetyl-TA) and kynurenine; Mixture 2 included tyramine, p-octopamine (p-OCT), normetanephrine, synephrine, methyltyramine and m-octopamine (also known as Norfenefrine) ; Mixture 3 had L-dopa, dopamine, 3,4-dihydroxy phenylacetic acid and Nw-acetylhistamine; Mixture 4 included tryptamine, 5-hydroxytryptophan (5-HTP), serotonin, and N-acetyltryptamine (N-acetyl-TPA); Mixture 5 contained spermine, spermidine, agmatine and putrescine. This screening identified five receptors responsive to one or more mixtures: Tadh153, Tadh165, Tadh168, Tadh170, and Tadh173 (Fig. 2). These receptors were subsequently tested with individual compounds from the mixtures they responded to at concentration 100 µM, to identify active compounds for further dose-response analysis (Supplementary Fig. 1). Monoamine receptors are known for their promiscuity, often being activated by multiple monoamines due to structural similarities (Xie, Westmoreland, and Miller 2008; Bauknecht and Jékely 2017). Thus, to identify the physiologically relevant ligands, we tested the individual molecules that produced signals at concentrations ranging from 1e-12 M to 1e-3 M and calculated EC50 values for each molecule’s dose response curve (Fig. 3A).

**Figure 2.**
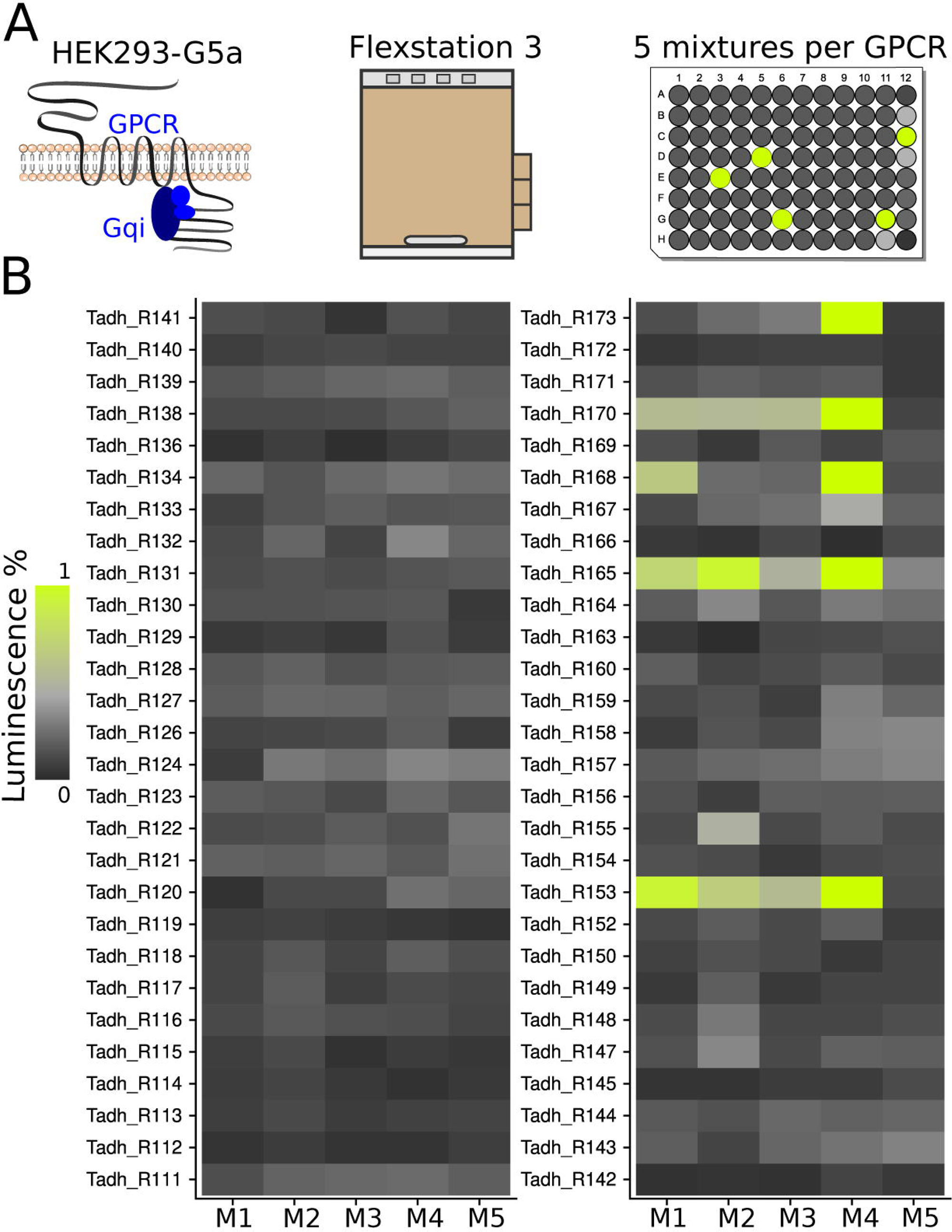
The primary screen of 56 Trichoplax GPCRs against five monoamine mixtures. (A) Pharmacological assay and pipeline for identifying monoamine-GPCR pairs. (B) Heatmap showing relative luminescence values from two replicates. Colour coding represents luminescence levels as shown in the key. Receptor responses were normalized to the negative control (0%) and five times the negative control (100%) to identify potential signals.

**Figure 3.**
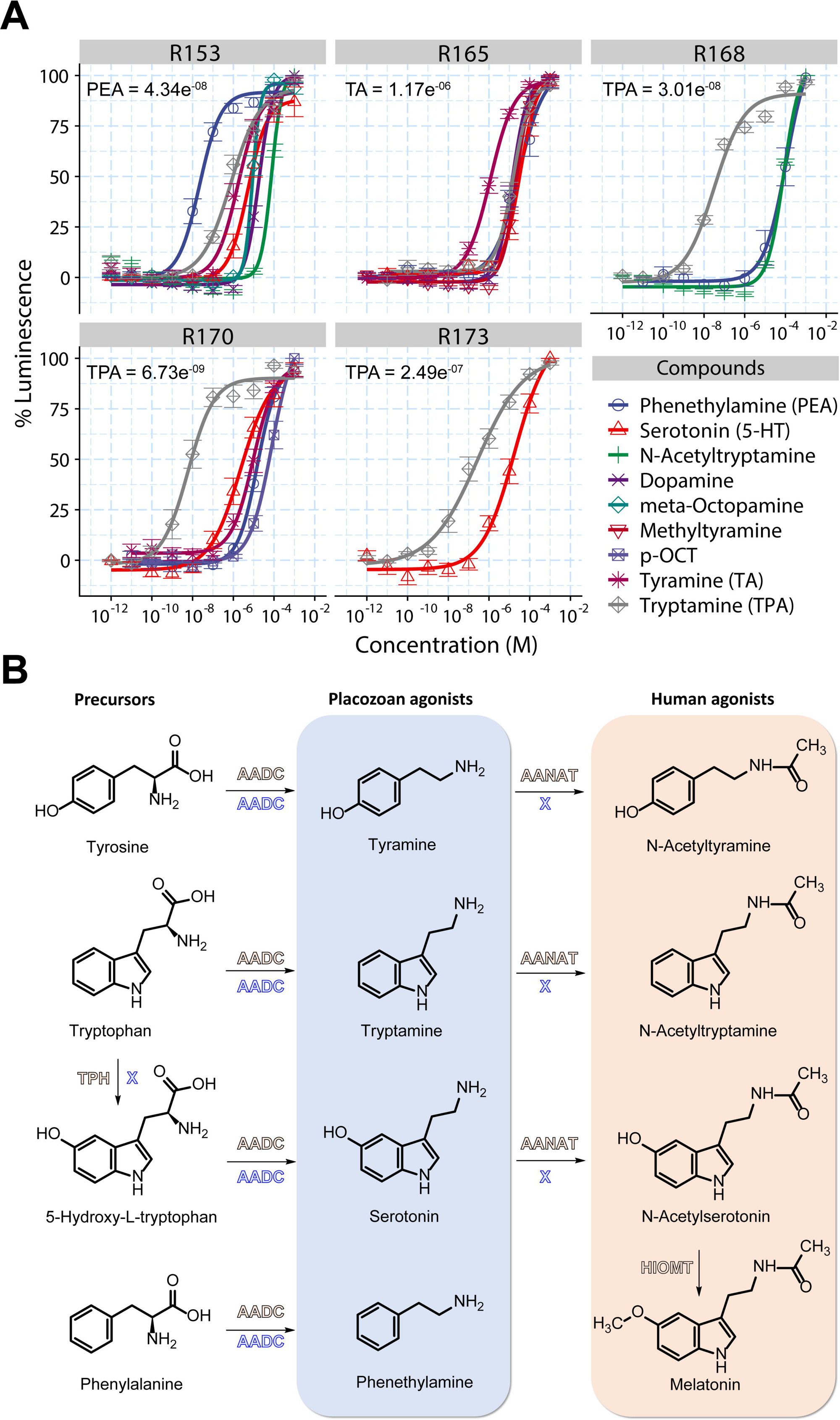
A) Dose-response curves of monoamine-GPCRs from *Trichoplax adhaerens*. Dose-response curves show normalized luminescence plotted against varying ligand concentrations for each monoamine-GPCR pair. The receptor number is indicated above each curve. EC_50_ values for the main ligands for each receptor are shown: phenethylamine for Tadh153, tyramine for Tadh165, and tryptamine for Tadh168, Tadh170, and Tadh173. The complete list of EC50 values for all tested compounds is available in Supplementary file 3. **B) Synthesis pathway of placozoan monoamine receptor agonists compared to the acetylated versions of the compounds that activate human melatonin receptors**. The enzyme AADC that is capable to produce phenethylamine, tyramine and tryptamine is present in placozoans, whereas the enzymes TPH and AANAT are absent in placozoans. Schemes of compounds where drawn using ChemDraw23.1.1

While these receptors displayed varying levels of promiscuity, certain molecules were more potent. Receptors Tadh168, Tadh170, and Tadh173 were most sensitive to tryptamine, with EC50 values of 30.1 nM (3.01e-08), 6.73 nM (6.73e-09), and 249 nM (2.49e-07), respectively. Receptor Tadh165 was preferentially activated by tyramine, with the lowest EC50 value of 1.17 µM (1.17e-06), and receptor Tadh153 responded best to phenethylamine, with an EC50 value of 43.4 nM (4.34e-08) (Fig. 3A and Supplementary File 3). Given the structural similarities of these various ligands, the other monoamines likely elicited responses due to cross-reactivity. For instance, Tadh170 and Tadh173 responded to both tryptamine and serotonin, which differ by a single hydroxyl group (Fig. 3B). However, the potency of serotonin on these receptors was significantly lower with EC50 values: 2.35 µM (2.35e-06) for Tadh170 and 17.7 µM (1.77e-05) for Tadh173 (Fig. 3A, Supplementary File 3). This corresponds to approximately 2–3 orders of magnitude difference in potency. Thus, we classified the receptors based on the molecule that produced the best response: Tadh168, Tadh170, and Tadh173 as tryptamine receptors, Tadh153 as a phenethylamine receptor, and Tadh165 as a tyramine receptor. These results demonstrate that placozoans possess receptors responsive to monoamines at physiologically relevant concentrations, with the primary activators being compounds that are likely found in placozoans, derived via a single decarboxylation step of amino acids by the aromatic amino acid decarboxylase (AADC) enzyme. Indeed, phylogenetic analysis with sequences extracted from available genomic data confirmed that placozoans possess a single AADC homologue, in the species *Trichoplax adhaerens*, *Trichoplax* H2, *Hoilungia hongkongensis*, and *Cladtertia collaboinventa* (Fig. 3B and Supplementary Fig. 2).

### Phylogenetic and receptor pharmacology analyses reveal homology between placozoan and bilaterian monoamine receptors

In response to previous research suggesting that monoamine receptors are absent in non-bilaterians, including placozoans, we sought to better define the phylogenetic relationships between our experimentally characterised placozoan monoamine receptors and bilaterian receptors. Thus, we conducted a large-scale phylogenetic analysis, including all receptors associated with the monoamine cluster, which according to the cluster-based analysis included receptors such as opsins and biogenic amine receptors including acetylcholine, adenosine, and various monoamine receptors (Fig. 1). We selected a set of neuropeptide receptors to root the tree, which according to our CLANS analysis formed a separate cluster from the monoamine receptors.

Among all deorphanised receptors, Tadh173 stood out as the only one not belonging to the melatonin receptor clade. Instead, it clustered within an expanded group of placozoan receptors related to several bilaterian orphan receptors, including GPR21, GPR52, GPR61, GPR101, and GPR161 (Fig. 4A, black dotted-line box). Homologous sequences within this receptor clade were also identified in ctenophores and cnidarians, suggesting the group predates the emergence of bilaterians. Notably, Tadh173 was the only receptor in this clade for which a ligand was identified.

**Figure 4.**
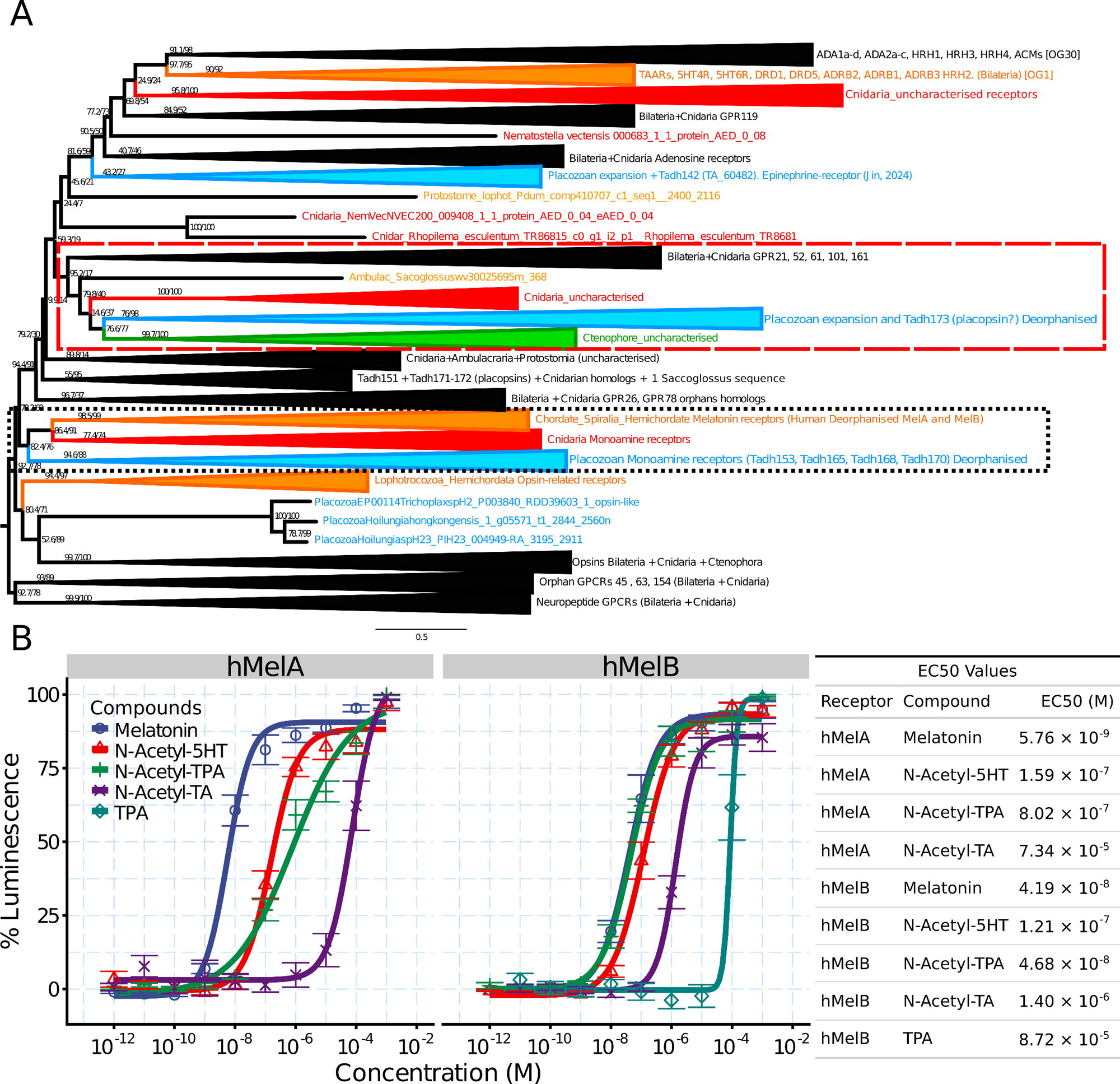
A) Maximum likelihood phylogeny of monoamine GPCRs. The tree was generated using IQ-TREE2 with the best-fit model LG+G4 and rooted with neuropeptide GPCRs. The tree is partially collapsed for clarity. Node support values (shown as numbers) are based on 1000 replicates. Clades are colour-coded as follows: red for cnidarians, blue for placozoans, orange for bilaterian-specific clades, and green for ctenophores. Black clades represent groups found in both bilaterians and non-bilaterians. Dotted or dashed boxes indicate clades containing receptors that were deorphanized in placozoans. **B) Experimental characterisation of human melatonin receptors (hMelA and hMelB) supports the homology relationship of monoamine GPCR between placozoans and humans.** Dose-response curves show normalized luminescence plotted against varying ligand concentrations for each monoamine-GPCR pair. The receptor name is indicated above each curve. The table adjacent to the curves lists EC50 values for each compound tested on human receptors. hMelB is activated by tryptamine only at high concentrations.

In contrast, our phylogenetic analysis revealed that four *T. adhaerens* monoamine receptors - Tadh153, Tadh165, Tadh168, and Tadh170 - are orthologous to bilaterian melatoninergic receptors, with this receptor type also being present in cnidarians and other placozoan species (Fig. 4A, red dotted-line box). In humans, three melatonin-related receptors were identified (MTNR1A, MTNR1B, and the orphan MTNR1L), while species such as *Saccoglossus kowalevskii* (a hemichordate) and *Platynereis dumerilii* (an annelid) possess a greater diversity, with 14 and 7 melatoninergic receptors, respectively. Interestingly, ecdysozoans like *Drosophila melanogaster* and *Caenorhabditis elegans* appear to have lost this receptor family, suggesting a secondary loss (Supplementary Fig. 3). In placozoans, we observed a significant expansion, with *T. adhaerens* harboring 18 melatoninergic receptors in total - four deorphanised (Tadh153, Tadh165, Tadh168, Tadh170), and 14 remaining orphan (Supplementary Fig. 3).

Given our observation that placozoan monoamine receptors are phylogenetically related to melatonin receptors, we decided to test, whether the human MTNR1A or hMelA and MTNR1B or hMelB get activated by similar molecules as the placozoan receptors using the same deorphanisation strategy. We tested the human receptors individually with all the compounds used on the placozoan receptors at 100 µM (Supplementary Figure 4), and compounds that gave detectable signals were used to generate dose-response curves. We found that both hMelA and hMelB showed no response to tyramine or phenethylamine, but tryptamine activated hMelB at high, non-physiological concentrations (Fig. 4). Instead, we observed that acetylated versions of the placozoan receptor agonists tryptamine, tyramine, and serotonin caused activation in human receptors. Specifically, for hMelB, N-acetyltryptamine showed comparable activation as the canonical ligand melatonin, with respective EC_50_ values of 46.8 nM and 41.9 nM. N-acetylserotonin (EC_50_ = 121 nM) and N-acetyltyramine (EC_50_ = 1.4 µM) were less potent as agonists, and TPA produced the weakest response requiring much higher, non physiological concentrations (EC_50_ = 87.2 µM). For hMelA, melatonin (EC_50_ = 5.56 nM) was the most potent activator, followed by N-acetylserotonin (EC_50_ = 159 nM), N-acetyltryptamine (EC_50_ = 802 nM), and N-acetyltyramine (EC_50_ = 73.4 µM). We could not obtain acetylated phenethylamine from a vendor, which potently activates the placozoan receptor Tadh153, to test its effect on the human melatonin receptors.

Recently, high resolution structures of the human MelA and MelB revealed shared mechanisms for ligand binding, requiring three conserved amino acids in the ligand binding pocket that form critical contacts with bound ligands (Stauch et al. 2019; Wang et al. 2022; Johansson et al. 2019). Specifically, the receptor residues asparagine 162 and glutamine 181 in MelA (*i.e.*, N162 and Q181), and N175/Q194 in MelB, form anchoring electrostatic interactions with oxygen atoms located on either end of melatonin and structurally-related analogues like 2-iodomelatonin (Fig. 5A) and ramelteon, while additional anchoring is provided by aromatic stacking between the hydrophobic core of the ligands and a conserved phenylalanine side chain in the receptors (F179 in MelA and F192 in MelB; Fig. 5A). Notably, our observation that MelA and MelB prefer acetylated monoamines is consistent with this structural data, wherein the oxygen atom in the melatonin alkylamide group, also present in the acetylated monoamines N-acetylserotonin, N-acetyltryptamine, and N-acetyltyramine, forms a key hydrogen bond with the noted Q181/Q194 residue in MelA and MelB receptors (Fig. 5A).

**Figure 5.**
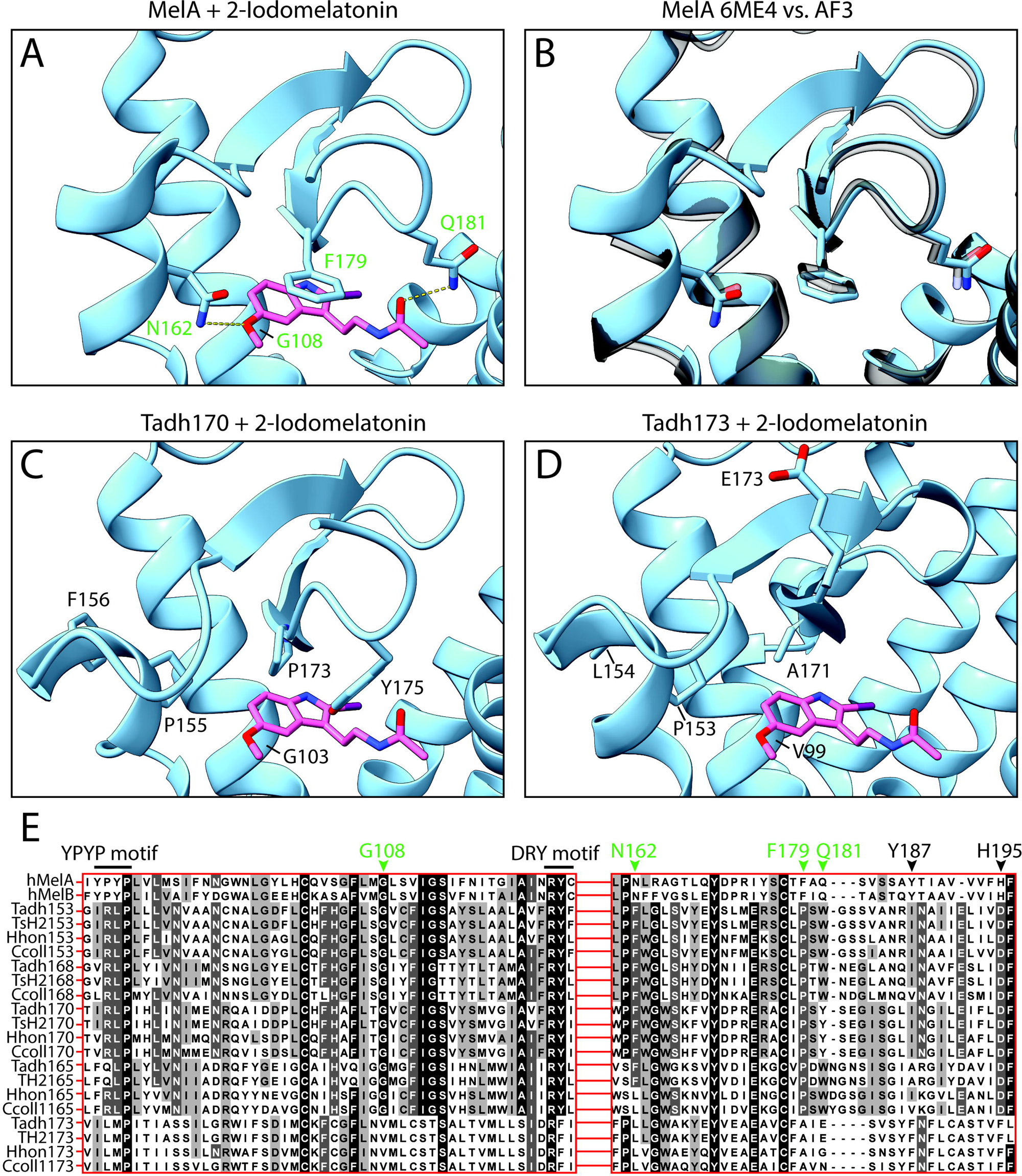
Structural and sequence comparisons of ligand binding between the human MelA receptor and placozoan monoaminergic receptors. **A)** Resolved structure of the human MelA receptor bound by 2-iodomelatonin (PDB number 6ME4). Hydrogen bonds between the N162 and Q108 side chains of the receptor and the methoxy and alkylamide groups of the ligand are shown with dashed yellow lines. **B)** Structural alignment of the resolved and predicted structures of the human MelA receptor. **C)** Superimposition of 2-iodomelatonin in the predicted structure of the *T. ahaerens* melatonin-related receptor Tadh170. **D)** Superimposition of 2-iodomelatonin in the predicted structure of Tadh173. For panels C and D, the position of 2-iodomelatonin is based on structural alignment between the solved MelA receptor structure (*i.e.*, PDB 6ME4) and the predicted structures of the placozoan receptors. **E)** Protein sequence alignments of regions of the human melatonin receptors and identified placozoan orthologues of the *T. adhaerens* monoaminergic receptors that were deorphanized in this study. Chevrons denote amino acids described in the text, numbered according to the human MelA receptor protein sequence. Key residues for binding melatonin analogues in MelA are colored in green. Also shown is the residue Y187 and H195, the former contributing to receptor activation through interactions with the N162 residue, and the latter through interactions with the bound ligand. Also annotated are the YPYP and DRY motifs flanking the ligand accommodating residue G108. All included placozoan receptors lack the first proline in the YPYP motif, a unique feature that distinguishes the human melatonin receptors and their closely related orphan receptor GPR50 from all other human GPCRs. Also notable, all melatonin clade receptors lack a canonical aspartate in the DRY motif, notable because alterations to this residue can increase the baseline activity of G protein signalling (Rovati, Capra, and Neubig 2007).

Leveraging these available structures, we sought to explore whether the binding of monoamines in the placozoan receptors might resemble that of human melatonin receptors through structural prediction and sequence alignments. First, we used AlphaFold3 to predict the structure of wildtype human MelA, since its experimental determination required several modifications to the amino acid coding sequence and chimeric fusion with a *Pyrococcus abyssi* glycogen synthase protein (Stauch et al. 2019). This AlphaFold structure was predicted with high confidence, producing a predicted template modeling (pTM) score of 0.8. Structural alignment with the solved MelA structure revealed low deviation in spatial positioning of aligned backbone alpha carbon atoms, producing a pruned root mean square deviation (rmsd) score of 0.568 angstroms. Accordingly, the predicted and solved structures exhibit similar positioning of the N162, Q181, and F179 side chains that interact with melatonin-related ligands (Fig. 5B). These observations encouraged us to similarly predict analogous wildtype structures of the placozoan receptors Tadh170 and Tadh173, the former being phylogenetically related to the human melatonin receptors, and the latter to the noted orphan receptors (Fig.4A). Both placozoan structures were predicted with high confidence, producing pTM scores of 0.8 (Tadh170) and 0.82 (Tadh173). To infer whether these receptors bear equivalent ligand-binding residues found in human melatonin receptors, we structurally aligned these with the solved structure of MelA bound to 2-iodomelatonin. This produced rmsd scores of 1.068 angstroms (Tadh170) and 1.198 angstroms (Tad173), which was not unexpected given the considerable sequence divergence between the human and placozoan proteins. Superimposing the 2-iodomelatonin ligand from the resolved MelA structure with the predicted placozoan structures revealed an absence of core determinants in the receptors for ligand binding. That is, structurally, both Tadh170 and Tadh173 were predicted to lack N162 equivalent residues with polar side chains positioned to hydrogen bonds with an oxygen atom within the 2-iodomelatonin methoxy group, and both lacked predicted Q181 equivalents side chains for hydrogen bonding with a flanking carbonyl oxygen within the ligand alkylamide group (Fig. 5C and D). Instead, both Tadh170 and Tadh173 were predicted to possess proline residues in the N162 equivalent structural position, and either tyrosine or glutamate in the Q181 position. In a protein alignment, the placozoan proline residues are one amino acid upstream of N162 of MelA and N175 of MelB. Instead, the conserved MelA/MelB asparagine residues align with a phenylalanine in Tadh170 (F156) and a leucine in Tadh173 (L154; Fig. 5E), which in the predicted structures, had side chains projecting away from the ligand binding pocket (Fig. 5C and D). Furthermore, both Tadh170 and Tadh173 lack the critical F179 equivalent required for strong hydrophobic contacts with the aromatic center of monoamine ligands, with Tadh170 bearing a proline in this structural position (which could possibly mediate mild aromatic interactions), and Tadh173 having a neutral alanine. In our protein alignment, we included sequences of all deorphanised monoaminergic GPCRs from *T. adhaerens*, as well as their orthologues from the placozoan species *Trichoplax* H2, *Hoilungia hongkongensis*, and *Cladtertia collaboinventa*. This alignment revealed a similar absence of N162/N175, F179/F192, and Q181/Q194 equivalents in any of these receptors. There is, however, conservation of a glycine below the plane of the bound ligand in all placozoan receptors except 173 which bears a valine in this position (V99). The minimal side chain of this residue in the human MelA receptor (*i.e.*, G108) permits occupancy of ligands in the binding pocket, since mutation to alanine, bearing the next-smallest side chain, completely disrupts ligand activation.

Taken together, these comparisons suggest the mechanisms for monoamine ligand binding are fundamentally different between human and placozoan receptors, despite the phylogenetic proximity of most *T. adhaerens* monoaminergic receptors to MelA and MelB. Of course, all this needs to be confirmed via structural determination.

### Monoamine Receptor Localisation and behavioural Effects in Placozoans

While our receptor activation assays demonstrate functional responses to monoamines, identifying the cells in which these receptors are expressed provides crucial insight into their physiological roles. To address this, we analysed expression datasets from recently published single-cell transcriptomes of four placozoan species (Najle et al. 2023), focusing on the distribution of monoaminergic receptors across different cell types. Additionally, we tested whether the identified monoamines elicit physiological effects in placozoans.

Expression analysis revealed that monoamine receptors are expressed in specific cell types across placozoan species. Some cell types show conserved expression patterns, while others exhibit species-specific differences. Gland cells and peptidergic cells are the primary cell types expressing these receptors, although their distribution varies among species. Receptor 153 is expressed in gland cells in both *Hoilungia hongkongensis* and *Trichoplax adhaerens*, with additional expression in peptidergic cells of *H. hongkongensis*. Similarly, *Trichoplax* species H2 also show expression in peptidergic cells. Receptor 165 is found in peptidergic cells of *H. hongkongensis* and *Cladtertia collaboinventa*, with additional expression in epithelial and gland cells of *H. hongkongensis*. Expression data of receptor 165 is absent in *T. adhaerens* and *Trichoplax* sp. H2. Receptor 168 is exclusively expressed in peptidergic cells in both *Trichoplax* species, with no expression detected in *C. collaboinventa* and no available data for *H. hongkongensis*. Receptor 170 is expressed in gland, peptidergic, and both upper and lower epithelial cells in *C. collaboinventa* and *H. hongkongensis*, but its expression is low in *T. sp. H2* and *T. adhaerens*. Finally, receptor 173 is consistently expressed in peptidergic cells across all species, with additional expression in the gland and epithelial cells of *T. adhaerens.* (Supplementary Fig. 5)

Following the identification of highly sensitive receptors for tryptamine, tyramine, and phenethylamine, combined with confirmation of their expression in placozoans, we hypothesized that exogenous application of these molecules would elicit physiological effects on *T. adhaerens* specimens *in vivo*. To test this, placozoans were exposed to different concentrations of these monoamines, and behavioural responses were recorded. Tyramine application at 50 µM induced significant increase of animal velocity, evident from an elongation of their average post-test path and substantial increase of average distance travelled between time points (Fig.6 A1). We further tested 10 µM tyramine, but it did observe any obvious change of *T. adhaerens* speed as compared with the pre-test measurements. The effect of 50 µM tyramine was observed for more than an hour after the application (Fig.6 A2), so the path travelled 60 minutes after tyramine application was further compared by an analysis of covariance (ANCOVA) between the control and two experimental groups (10 and 50 µM), using respective path travelled 60 minutes before the application as a covariate. The test showed significant difference between control animals and animals treated with 50 µM tyramine (p=3.7E-05); on average, *T. adhaerens* in this experimental group covered 11.6 mm (mean value corrected for covariance; n= 28) distance after adding tyramine, while control animals covered ∼8.6 mm (n=21) distance after adding vehicle solution (Fig. 6 A3). No drastic changes in body area and shape were detected in the experiments with tyramine.

**Figure 6.**
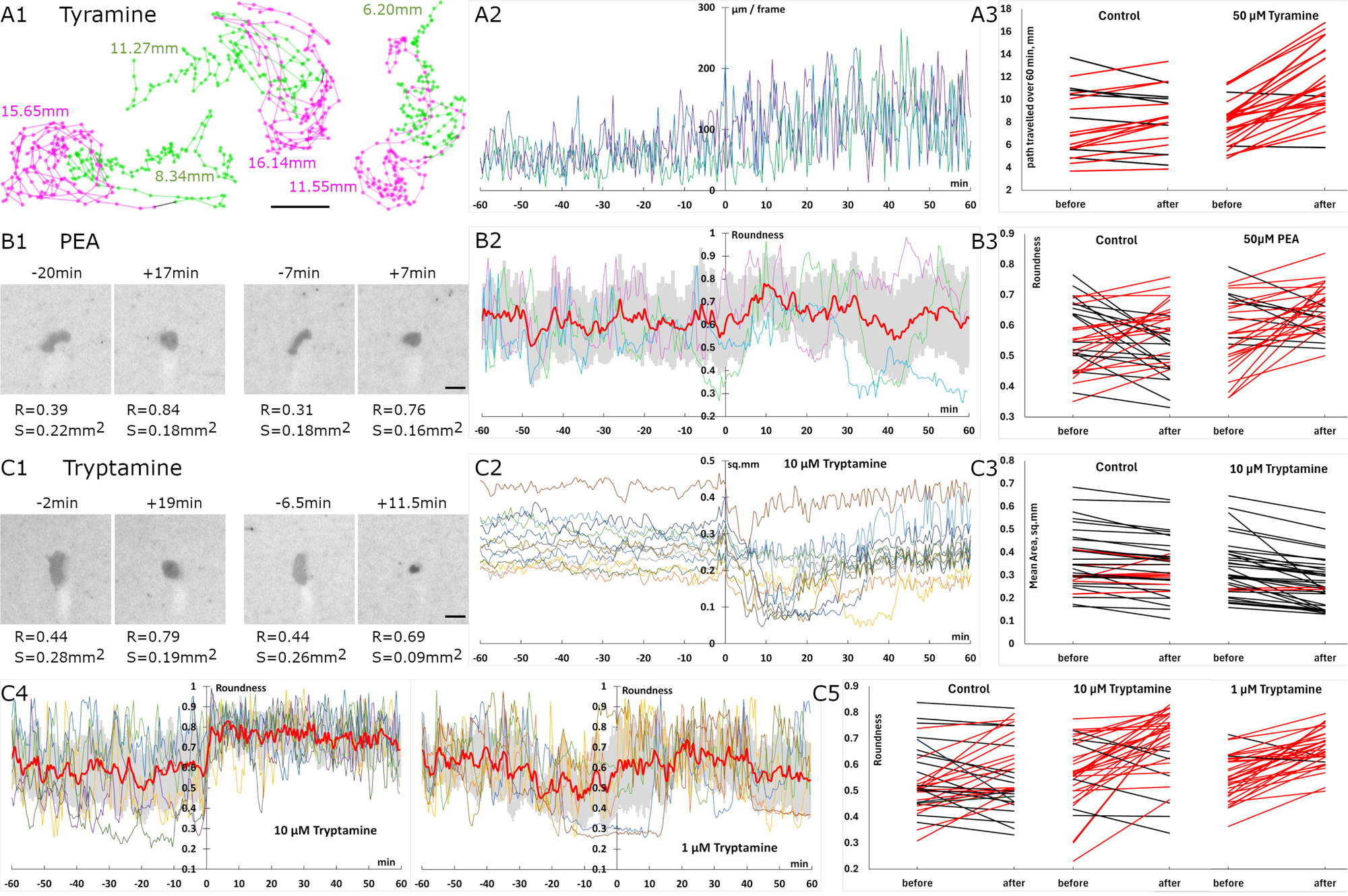
Effect of exogenous application of Tyramine (A), PEA (B), and Tryptamine (C) on Trichoplax behaviour. B1 and C1 show representative images of the animals: the time before (-) or after (+) application is indicated above the image; below the image are roundness value (R) and surface value (S). Charts on A-C2, and C4 show parameters of a few individual Trichoplax plotted over time; a monoamine is added at t=0; thick red line and gray shading indicate mean value and standard deviation, respectively, on B2 and C4. Charts on A-C3 and C5 show how the mean value of the parameter changes upon the application in each individual Trichoplax counted: The increase and decrease are color coded red and black, respectively. A1 – trajectories of three animals recorded for one hour before (green) and after (magenta) 50 µM tyramine application (gray). The values of respective color indicate the distance travelled by the animals during one hour. A2 – dynamic of the distance travelled in three individuals between each frame (30 seconds apart): Note the increase after the application. A3 –the path traveled during one hour in the control and experimental groups before and after the application. B1 – After 50 µM PEA application, Trichoplax round up and contract slightly. B2 – increase in roundness upon PEA application (three individuals plus mean). B3 – changes in roundness in the control vs experimental group. C1 – 10 µM tryptamine elicits crinkling. C2 – area of 12 animals drops significantly after tryptamine application. C3 – decrease of surface area in control vs tryptamine treated group. C4 – roundness variations (6 individuals plus mean) upon 10 µM (left) or 1 µM (right) of tryptamine. C5 – increase of mean roundness in control and in animals treated with different concentrations of tryptamine. Scale bar on A-C1 represents 0.5 mm.

Phenethylamine at 50 µM triggered moderate, yet significant, rounding of *T. adhaerens* body outline, which was apparent during first 30 minutes after application (Fig. 6 B1). During a pre-test time, animals constantly changed their shapes from curved to elongated and to rounded and back, as they would normally do in a culture. However, upon phenethylamine application, the animals started rounding up in about 5 minutes and stayed rounded most of the time for the next 15-25 minutes (Fig.6 B2); after this period animals recovered and resumed their normal stereotypical behaviours. Hence, a 30 min interval was chosen for further statistical analysis. The mean corrected roundness of the control animals was 0.56 (N=34) during the post-test period, while phenethylamine treated animals had mean corrected roundness of 0.65 (N=33); this change was significant at p value of 7.7E-05 (Fig. 6 B3). Although phenethylamine caused animal reshaping, it did not affect the area; likewise, no significant changes were revealed in the velocity.

Tryptamine elicited a very strong contraction of *T. adhaerens* body, reflected by a sharp decrease of the area and increase of roundness, at 50 µM concentration. The effect was disproportionally extreme on larger animals, making them shrink more than twice of a pre-test value. We considered this concentration as too high and did not include the results in further analysis since the potential interaction between the covariate and the independent variable violates ANCOVA assumptions. 10 µM of tryptamine induced consistent and significant contraction of the body during first 30-40 minutes after application regardless of the animal initial size (Fig. C1 and C2). The observed contraction was associated with crinkling and folding of *T. adhaerens*; some animals completely folded up. The ANCOVA test showed significant difference between averaged area scores obtained 30 min before and after application (p=7.0E-05). In the experimental group the post-test area was equal to 0.27 mm2 (N=35) vs 0.31 mm2 in the control (N=35) (Fig. C3). Along with the area drop, 10 µM tryptamine also induced rounding of the body outline, which was observed even when the body area was back to normal, *i.e.*, beyond 30 min after application (Fig. C1 and C4). The changes were significant (p=5.3E-05): The roundness increased to 0.68, while the control group had this value at 0.57. Since 10 µM of tryptamine demonstrated a strong effect, we decreased concentrations and tested the effect of 1 µM of tryptamine. 1 µM concentration had a more moderate effect on *T. adhaerens* reshaping: while we still observed significant (p=4.4E-04) increase in roundness (mean corrected roundness after application was equal to 0.67, N=34; Fig. C4 and C5), changes in the area were subtle. No change in the traveling paths was noted at any tested tryptamine concentration.

## Discussion

The origin of monoaminergic signalling systems remains a topic of debate. Over the years, immunohistological, behavioural, and physiological evidence has demonstrated monoamine functions in various non-bilaterian animals. For instance, dopamine and epinephrine have been implicated in coral (cnidarian) larval settlement (Moeller, Nietzer, and Schupp 2019), serotonin in hydrozoan larvae metamorphosis (Zega et al. 2007; McCauley 1997), while epinephrine influences locomotion and feeding in the placozoan *T. adhaerens* (Jin et al. 2024). A recent study also showed that sponges respond to trace amines such as tyramine and phenethylamine, where application of these monoamines induced significant changes in larval phototaxis in the species *Amphimedon queenslandica* (Xiang et al. 2023), suggesting the existence of functional receptors. A central question that emerged from such previous studies is what receptors could be mediating these responses. As early as 1997, serotonin-sensitive receptors were proposed to exist in the cnidarian sea pansy *Renilla koellikeri* (Hajj-Ali and Anctil 1997). By employing radioligand binding techniques with [³H]5-HT, the authors identified high-affinity serotonin binding sites (Kₐ 23–34 nM), with a receptor density of 8.5–20.6 pmol/mg protein in cell membrane preparations from this species (Hajj-Ali and Anctil 1997). Although it was not possible to ascertain the sequence identity of these receptors, these findings suggested, for the first time, the occurrence and evolutionary conservation of monoaminergic receptors in non-bilaterians. However, alternative interpretations exist, with some studies, including recent research, suggesting the monoamine system may not be present in non-bilaterian animals (Goulty et al. 2023). The difficulty in identifying and experimentally characterising receptors in non-bilaterians has likely contributed to the view that a functional monoaminergic system is absent in these organisms.

### Discovery and pharmacological characterisation of monoamine receptors in *Trichoplax adhaerens*

In this study, we performed comprehensive bioinformatic analyses and large-scale deorphanisation assays to identify direct receptor-ligand interactions. Through this approach, we identified many candidate monoamine receptors in *T. adhaerens*, 56 of which we functionally tested *in vitro* (Fig. 1). Among these, five receptors exhibited high sensitivity to monoamines (Fig. 2) and dose-response assays allowed us to identify phenethylamine (Tadh153), tyramine (Tadh165), and tryptamine (Tadh168, 170, and 173) as the most active ligands (Fig. 3A). With EC_50_ values well within the expected physiological range, demonstrating the existence of functional monoamine receptors in placozoans. The activation of placozoan receptors by these simple monoamines aligns with the presence of a single enzyme responsible for monoamine synthesis in these organisms - aromatic amino acid decarboxylase (AADC) (Fig. 3B and Supplementary Fig. 2). AADC catalyses the synthesis of tyramine, phenethylamine, and tryptamine through a single decarboxylation step (Zhu et al. 2020; Speight et al. 2000; H. Zhang et al. 2019; M. D. Berry et al. 1996). This is not the first study to report simple monoamines as the most active compounds in non-bilaterians. For instance, tyramine and phenethylamine significantly altered larval phototaxis in *Amphimedon queenslandica*, whereas more complex monoamines, such as dopamine—which requires multiple enzymatic steps for synthesis—had a much weaker effect (Xiang et al. 2023). This pattern may also explain why placozoan tryptaminergic receptors exhibited only partial activation by serotonin, with 2-3 orders of magnitude weaker potency than tryptamine (Fig 3B). As discussed before, the tyrosine hydroxylase that participates in the production of serotonin in bilaterians is absent in placozoans (Francis et al. 2017), and consistent with this, capillary electrophoresis with laser-induced fluorescence detection in *Trichoplax adhaerens* failed to detect serotonin or dopamine as endogenous molecules (Romanova et al. 2020).

### The pre-bilaterian origin of monoaminergic receptors

After experimentally characterising monoamine receptors in placozoans, we examined their evolutionary relationship to bilaterian counterparts. To this end, we conducted a large-scale phylogenetic analysis incorporating all receptors from the monoamine receptor cluster identified via CLANS. As shown in Fig. 1, this group included opsins and various biogenic amine receptors, such as those for acetylcholine, adenosine, and multiple monoamine receptors such as melatonin, adrenergic, serotonergic, etc. In 2023, Goulty et al. attempted a similar approach but used a highly stringent CLANS threshold (P-value < 1e−42), likely excluding bona fide non-bilaterian receptors and explaining their failure to detect certain divergent monoamine receptors. For example, their analysis did not recover the placozoan receptors recently shown to be activated by epinephrine (Jin et al. 2024), nor any of the candidates we identify here. Furthermore, their phylogenies report melatonergic receptors in cnidarians but not in placozoans (Goulty et al. 2023). Here, we used a less stringent P-value (< 1e−28) for our GPCR clustering. As comparison, in previous research, we successfully identified neuropeptide receptors in *Nematostella vectensis* using a similarly relaxed threshold (P-value < 1e−25) (Thiel et al. 2024). Then, our phylogenetic analyses revealed a clear relationship between the placozoan monoamine receptors and bilaterian melatonergic receptors (Fig. 1 and Fig. 4), with Tadh153, Tadh165, Tadh168, and Tadh170 (Fig. 4A, Supplementary Fig. 3) identified as orthologous to human melatonin receptors with a high bootstrap support (>75% for both SH-aLRT and ultrafast bootstrap test).

To further resolve the homology between placozoan and bilaterian receptors we performed comparative pharmacological testing. Specifically, we tested human melatonin receptors against the same compound library used to deorphanise placozoan monoamine receptors. Both placozoan and human receptors responded to structurally similar molecules, albeit with a key difference, a methylation. Placozoan receptors were activated by non-acetylated monoamines like tryptamine, tyramine, serotonin, and phenethylamine (Fig. 3A, B). In contrast, human melatonin receptors responded to their acetylated counterparts, such as N-acetyltryptamine, N-acetyltyramine, and N-acetylserotonin (Fig. 4B). Although tryptamine activated the human MelB receptor (hMelB), it acted only as a partial agonist with a very high EC₅₀. These results are consistent with a previous study (Chen et al. 2023), which identified tryptamine as a MelB agonist but did not report detailed EC₅₀ values. N-acetylserotonin and N-acetyltryptamine have previously been established as ligands for melatonin receptors (Dubocovich et al. 1997). However, to our knowledge, this study represents the first demonstration that N-acetyltyramine acts as an agonist of human melatonin receptors.

The differences in ligand selectivity between placozoan and human receptors likely reflect evolutionary divergence in monoamine biosynthesis. While placozoans rely solely on AADC for monoamine synthesis, bilaterians have evolved additional enzymatic pathways, expanding their repertoire of synthesized monoamines. This expansion may have driven the functional specialization of human receptors, enabling them to recognize distinct ligands. Together, our phylogenies and comparative pharmacology demonstrate that placozoan monoamine receptors are orthologous to bilaterian melatonin receptors and that, like their placozoan counterparts, human melatonin receptors exhibit a broad ligand promiscuity.

### The “placopsin” Tadh173 is a tryptamine receptor

Among the monoamine receptors deorphanised in placozoans, the only one that did not show homology to human melatonin receptors is Tadh173—a receptor with low promiscuity, responding only to tryptamine and serotonin. Interestingly, Tadh173 has been annotated in previous bioinformatic analyses. In 2012, it was classified as a putative opsin, annotated as “placopsin” (XP_002113363) (Feuda et al. 2012). More recently, Jin et al. (2024) referred to this receptor as TA_3759 and proposed it as a potential epinephrine receptor based on their phylogenetic analysis. However, their functional assays only tested epinephrine—not serotonin or tryptamine, the actual ligands of this receptor—and knockdown of the gene did not produce detectable physiological effects following epinephrine treatment (Jin et al. 2024). Our phylogenetic, clustering, and deorphanisation data support its classification as a monoamine receptor, aligning it with the broader monoaminergic system rather than with opsins (Fig. 4A). In our phylogenies, Tadh173 is more closely related to the bilaterian orphan receptors GPR21, GPR52, GPR61, GPR101, and GPR161. Two additional placozoan receptors, Tadh171 (XP_002112438) and Tadh172 (XP_002112437) —also labelled as “placopsins” by Feuda et al. (2012)— were cloned and tested, but they did not respond to any of the compounds added. Although previously classified as placopsins along with Tadh173 (Feuda et al. 2012), in our phylogenies these two genes (Tadh171 and Tadh172) clustered within a distinct clade containing cnidarian homologues. This clade is nested within the broader monoamine receptor group, which also includes Tadh173 and bilaterian serotonin, dopamine, epinephrine, histamine, and trace amine receptors. Together, our phylogenetic and pharmacological evidence supports the reclassification of Tadh173 (XP_002113363, TA_3759) as a tryptamine receptor, rather than an opsin.

The placozoan clade of GPCRs most closely related to the bilaterian monoamine receptors—including those for serotonin, dopamine, epinephrine, histamine, and trace amines—comprises the *T. adhaerens* paralogous receptors Tadh149, Tadh148, Tadh142, Tadh150, Tadh147, Tadh144, and Tadh111 (Fig. 4A). None of these receptors responded to the monoamines in our compound panel when tested using the deorphanisation assay. It is possible that this entire clade is activated by a different class of molecules unrelated to canonical monoamines, or that our assay system does not fully recapitulate the downstream signalling complexity of placozoan GPCRs.

Notably, Tadh142 was identified by Jin et al. (2024) as a putative epinephrine receptor—referred to as TA_60482—based on knockdown experiments. We were unable to test its response to epinephrine directly, as HEK293 cells, used in our deorphanisation assays, endogenously express adrenergic receptors, making them unsuitable for assaying responses to adrenaline or noradrenaline (Cullum, Veprintsev, and Hill 2023; Cheng et al. 2010; Atwood et al. 2011). However, while TA_60482 showed a knockdown phenotype consistent with epinephrine signalling, placozoans lack the full biosynthetic pathway for epinephrine (Jin et al. 2024). Furthermore, our phylogenetic analyses support the classification of these receptors—including Tadh142 and its paralogues—as homologues to the broader bilaterian monoamine receptor clade. Thus, they are not one-to-one orthologues of specific receptor subtypes such as epinephrine receptors. Instead, these placozoan receptors form a many-to-many homology with multiple bilaterian monoamine receptor subtypes, suggesting they may have evolved independently and do not directly correspond to any single bilaterian receptor class.

### Expression and behavioural data

While genomic/transcriptomic analyses revealed the presence of monoamine receptor genes, this information alone does not confirm if these receptors are actively expressed or functionally relevant within the organism. Thus, we explored the expression of monoamine receptors in placozoans and assessed the physiological effects of monoamines on these organisms. These receptors are mostly expressed in gland and peptidergic cells across different placozoan species. Expression of monoamine receptors in peptidergic cells suggests an ancient, conserved mechanism where monoamines modulate neuropeptide release. The interplay between monoamines and neuropeptides for modulation is very well documented in bilaterians. In these organisms, monoamines like octopamine and tyramine differentially regulate neuropeptide release at specific sites, influencing signalling pathways (Clark et al. 2018; Komuniecki et al. 2012). This modulation likely occurs through signalling cascades that amplify or inhibit behavioural responses via neuropeptide release (Mills et al. 2012; Hapiak et al. 2013). However, while the expression of receptors is prominent in peptidergic cells, it is not exclusive to them. In fact, all receptors except Tadh168 are also present in other epithelial cells. This suggests that trace amines can also influence these cells directly.

To investigate the functional role of monoamines in placozoans, we identified molecules with the highest affinity for placozoan receptors and conducted behavioural assays. Our experiments revealed distinct effects of monoamines on placozoan behaviour. Tyramine at 50 µM significantly increased the speed of *T. adhaerens*, enhancing their locomotor activity for over an hour, possibly by increasing ciliary beating frequency. This finding aligns with previously published data by Jin et al. (2024), which also reported increased movement upon tyramine application (Jin et al. 2024). However, unlike that study, which additionally observed body crinkling and folding, we did not detect these morphological changes. We believe this discrepancy is due to the much higher concentration of tyramine used in that study (10 mM), which exceeds physiological levels and could have led to artefactual effects.

Phenethylamine at 50 µM caused moderate rounding of *T. adhaerens* body outline, most apparent during the first 30 minutes post-application. Tryptamine at 10 µM induced significant contraction and rounding of the *T. adhaerens* body, characterised by crinkling and folding, with some individuals completely folding up. Body reshaping in Trichoplax is closely linked to cell reshaping, particularly apical constriction, with epithelial cells being the primary drivers. The downstream signalling initiated by both phenethylamine and Tryptamine can ultimately lead to cytoskeletal remodeling and subsequent cell reshaping. Given that the phenethylamine receptor (Tadh153) and some of the Tryptamine receptors (Tadh173) are expressed in epithelial cells, it is possible that these monoamines directly influence their cytoskeletal organization. Alternatively, this effect might be mediated by neuropeptidergic signalling, as both phenethylamine and all Tryptamine receptors are expressed in the peptidergic cells.

These findings demonstrate that monoamines such as tyramine, phenethylamine, and tryptamine play crucial roles in modulating placozoan behaviour, affecting both locomotion and body shape. The differential responses to these monoamines suggest that placozoan monoamine receptors are involved in diverse physiological processes, similar to their roles in bilaterian species.

### Release mechanisms of tyramine, tryptamine and phenethylamine

Previous bionformatic analyses have shown that placozoans lack the vesicular monoamine transporter (VMAT), suggesting that monoamines may not be packaged into vesicles for exocytotic release (Goulty et al. 2023). This absence implies that monoamines could be retained within the cells where they are synthesised, limiting the potential for synaptic transmission. However, unlike other monoamines, tryptamine and phenethylamine are not tipycally stored in synaptic vesicles. Evidence has shown that they can be released directly from the cytoplasm of presynaptic terminals without the need of vesicles, through passive diffusion ((Mark D. Berry 2004; Kosa et al. 2000; Mark D. Berry et al. 2013). This release mode is partly due to their chemical structure: these trace amines lack ring hydroxyl groups (Fig. 3B), which reduces their polarity and increases their membrane permeability. As a result, they can more readily diffuse across lipid bilayers, including cellular membranes and, as demonstrated in rats, the blood-brain barrier (Oldendorf 1971). Indeed, these molecules are known to cross bilayers much more rapidly than dopamine or serotonin, enabling swift access to targets in the synaptic cleft (Mark D. Berry et al. 2013, 2016). Berry and colleagues (2013) provided direct experimental evidence for the passive release mechanism of tyramine, tryptamine, and phenethylamine by measuring their diffusion across lipid bilayers, reporting permeability coefficients of at least 20 Å/sec. This supported the hypothesis that simple diffusion, rather than vesicular release, serves as the primary mechanism of release for these trace-amines (Mark D. Berry et al. 2013, 2016). We propose that a similar passive mechanism underlies monoamine signalling in placozoans, which lack detectable neuron-like cell types.

Still, the possibility of vesicular packaging for other monoamines in non-bilaterians remains. Although VMAT and VAChT were shown to be absent in placozoans, cnidarians, and ctenophores (Goulty et al. 2023), a closer examination of their data reveals additional insights. In their phylogenetic analysis of vesicular transporters (Figure 4a and Supplementary Fig. 7; Goulty et al. 2023), the authors identified homologues of the vesicular polyamine transporter (VPAT, or SLC18B1) across all major non-bilaterian phyla: sponges, cnidarians, ctenophores, and placozoans. For instance, in *Trichoplax* sp. H2, they identified four VPAT homologues: *TrichoplaxspH2_4510323*, *TrichoplaxspH2_4506092*, *TrichoplaxspH2_4510316*, and *TrichoplaxspH2_4510324*. Since its initial functional characterisation, the mammalian VPAT has been shown to transport serotonin and polyamines (Hiasa et al. 2014). Subsequent studies confirmed its multi-specificity, including the ability to also transport acetylcholine with low affinity, demonstrating the multi-modality of this transporter (Takeuchi et al. 2017; Moriyama et al. 2020). Therefore, VPAT may serve dual roles in non-bilaterians—as both a polyamine and monoamine transporter—particularly in species where this is the only vesicular transporter present.

### Conclusion

Our identification of functionally responsive monoamine receptors from *Trichoplax adhaerens* underscores the importance of experimental validation in evolutionary genomics. With the ever-increasing access to high-quality genomic data across diverse lineages, new opportunities arise for re-evaluating previous phylogenetic assumptions. When combined with functional and structural analysis, this re-examination can illuminate the evolutionary origins and diversity of key cellular processes, such as those involved in cellular signalling. Here, we provide compelling evidence for the existence of monoaminergic signalling in non-bilaterians. Specifically, we demonstrate that non-bilaterian GPCRs can be activated by monoamines at low, physiologically relevant concentrations—highlighting conserved receptor function that would be difficult to infer from sequence data alone. As shown here, large-scale pharmacological assays can uncover conserved signalling mechanisms that may be missed due to sequence divergence, incomplete annotations, or stringent homology criteria. Nonetheless, many open questions remain. How do these receptors signal intracellularly? Are there additional placozoan monoamine receptors that our in vitro assay failed to detect? What are the mechanisms of ligand binding in the receptors we identified, and how do the structural determinants of monoamine binding compare across different receptor subtypes? Finally, how are monoamines stored and released in non-bilaterians, and are the vesicular transport mechanisms conserved? These questions present exciting opportunities for further exploration.

## Methodology

### Cluster-based analysis (CLANS) for receptor identification

To identify potential monoamine receptors in placozoans, we selected a set of monoamine-deorphanised receptors from humans, *Drosophila melanogaster*, and *Platynereis dumerilii*. The *Drosophila* and human receptor annotations were obtained from UniProt, while the *Platynereis* receptors were sourced from those previously characterised in this species (Bauknecht and Jékely 2017). Using this dataset, we generated our own Hidden Markov Model (HMM) using HMMER (v3.1b2) and applied it to search the proteomes of various species, Bilateria (*Drosophila melanogaster, Caenorhabditis elegans, Octopus vulgaris, Homo sapiens, Petromyzon marinus, Platynereis dumerilii, Saccoglossus kowalevskii*), Cnidaria (*Alatina alata, Hydra vulgaris, Kudoa iwatai, Nematostella vectensis, Polypodium hydriforme, Rhopilema esculentum, Tetracapsuloides bryosalmonae*), Placozoa (*Hoilunga hongkongensis, Trichoplax adhaerens, Trichoplax H2, Cladtertia collaboinventa*), Porifera (*Amphimedon queenslandica, Ephydatia muelleri, Oscarella carmella, Sycon ciliatum, Tethya wilhelma*), Ctenophora (*Pleurobrachia bachei, Mnemiopsis leydi, Hormiphora californiensis and Bolinopsis mikado*), and the single-celled eukaryotes; *Monosiga brevicollis, Salpingoeca rosetta, Capsaspora owczarzaki, Tunicaraptor unikontum*, using an e-value threshold of 1e−10. Redundant sequences were removed using CD-Hit with a 95% similarity threshold. This approach identified not only monoamine receptors but also opsins, neuropeptides, olfactory receptors, and others. This is because monoaminergic receptors share sequence similarity and are phylogenetically related to neuropeptide and opsin receptors, a relationship that has been described before (Thiel et al. 2024). Olfactory receptors, which form a well-defined cluster restricted to chordates, were excluded from further analyses. As a result, we obtained 3,166 GPCRs that were used to perform a cluster-based analysis CLANS, that uses similarity to identify related sequence clusters. The sequences used for the CLANS analysis are available in Supplementary File 4.

Relationships between receptor sequences from different species were examined using an all-vs-all BLAST-based clustering approach with CLANS. The initial BLAST file was generated via the CLANS online toolkit with the default BLOSUM62 scoring matrix, extracting BLAST HSPs with e-values up to 1e−10. In previous analyses, Goulty et al. attempted to identify monoaminergic receptors using a similar CLANS analysis with a very stringent threshold (P-value < 1e−42), which limited the identification of distantly related homologues. To avoid this limitation, we relaxed the CLANS threshold to a P-value of <1e−28. Cluster annotation was based on receptors from *Drosophila melanogaster* and *Homo sapiens*. All receptors from *Trichoplax adhaerens* connected to the cluster containing monoamine-related receptors from bilaterians were selected. The number of transmembrane domains was predicted using Phobius (Käll, Krogh, and Sonnhammer 2007), and only receptors with seven transmembrane domains were chosen for receptor deorphanisation. These were codon-optimized and synthesized by Genscript, then cloned into the pcDNA3.4 vector. Fifty-six receptors were selected for further analysis. The protein sequences are provided in Supplementary File 1, and the codon-optimized sequences in Supplementary File 2.

### Large-scale receptor deorphanisation

Based on the CLANS analysis, 56 receptors were selected and analyzed for 7TM. Sequences connected to the monoamine cluster and possessing 7TM domains were sent for synthesis and cloning by Genscript in the vector pcDNA3.4. These receptors were transformed in the laboratory and extracted using the Macherey-Nagel endotoxin-free kit (cat no. 11932492), following the manufacturer’s instructions. The plasmids were then used for large-scale combinatorial deorphanisation. The protocol used for deorphanisation has been detailed before (Thiel et al. 2024) and can be accessed via bio-protocols (Thiel, Yañez-Guerra, and Jekely 2023). Below is a summary of the critical steps. HEK293 cells stably expressing the chimeric GFP-Aequorin protein G5A (Product no. cAP-0200GFP-AEQ-Cyto) were cultured in DMEM supplemented with 4.5 g/l glucose, L-Glutamine, sodium pyruvate (Thermo; Cat. No. 10566016), and 10% heat-inactivated FBS (Thermo; Cat. No. 10082147) under 5% CO₂ conditions at 37°C. When cells in a T75 flask reached 80%-90% confluency, they were distributed into 3 clear-bottom 96-well plates and cultured for two days. Upon reaching 90% confluency, transfection was performed using PEI-STAR (Biotechne, Cat. No. 8174) (1 mg/ml), following Durocher et al. (Durocher, Perret, and Kamen 2002). The medium was replaced with 90 µl of 5% FBS-supplemented OptiMEM, and the transfection mix for each well (10 µl of OptiMEM without FBS, 60 ng each of GPCR-containing plasmid and Gqi9 plasmid, and 0.45 µl of PEI) was incubated at room temperature for 20 minutes before being added to the cells. Two days posttransfection the media of the cells was removed and substituted by HBSS 1x media (Gibco; 14025092) supplemented with coelenterazine-H 4 µM (Promega; Cat. No. S2001) and incubated for a period of 2–3 hr. The readings were performed with a FlexStation 3 Multi-mode Microplate reader (Molecular Devices) for 60 seconds per well, with ligand injection after 15 seconds. The whole plate was read with the Flex option.

Pooled mixtures of compounds were used for the initial receptor characterisation: **Mixture 1:** 2-Phenethylamine (CAS 156-28-5, Sigma), Melatonin (CAS 73-31-4, Sigma), N-Acetylserotonin (CAS 1210-83-9, Cayman Chemical), N-Acetyltyramine (CAS 1202-66-0, MCE), and DL-Kynurenine (CAS 343-65-7, Sigma). **Mixture 2:** Tyramine hydrochloride (CAS 60-19-5, Sigma), (±)-Octopamine hydrochloride (CAS 770-05-8, Sigma), DL-Normetanephrine hydrochloride (CAS 1011-74-1, Sigma), Synephrine Hydrochloride (CAS 94-07-5, BIOSYNTH), N-Methyltyramine HCl (CAS 13062-76-5, BIOSYNTH), and *m*-Octopamine (4779-94-6, Sigma [also known as norfenefrine or norphenephrine]). **Mixture 3:** Levodopa (CAS 59-92-7, Sigma), Dopamine hydrochloride (CAS 62-31-7, Sigma), 3,4-Dihydroxyphenylacetic acid (CAS 102-32-9, Sigma) and Nω-Acetylhistamine (CAS 673-49-4, Sigma). **Mixture 4:** Tryptamine hydrochloride (CAS 343-94-2, Thermo), 5-Hydroxy-L-Tryptophan (CAS 4350-09-8, Sigma), Serotonin hydrochloride 98% (CAS 153-98-0, Thermo), N-Acetyltryptamine (CAS 1016-47-3, Thermo) N-Acetylserotonin (CAS 1210-83-9, Sigma). **Mixture 5:** Spermine (CAS 306-67-2, Sigma), Spermidine (CAS 334-50-9, Sigma), Agmatine (CAS Sigma), and Putrescine (CAS 333-93-7, Sigma).

The compounds were diluted into a 100 mM stock solution according to their solubility in water, DMSO, or ethanol. This stock was then used to prepare the mixtures or dose-response concentrations by diluting the compounds in 1× HBSS containing 100 µM ascorbic acid (CAS 50-81-7, Sigma). Due to observed oxidation, solutions were prepared fresh for each experiment, even with the addition of ascorbic acid as an antioxidant. This stock was then used to prepare the mixtures or dose-response concentrations by diluting the compounds in 1× HBSS containing 200 µM ascorbic acid (CAS 50-81-7). Mixtures and individual compound screening assays were tested at the highest concentration of 100 µM (1×10⁻⁴ M). For the initial analysis, mixtures were tested on transfected receptors in at least two independent transfections. Receptors showing responses to the mixtures were then tested individually with all compounds to identify specific responses. Compounds producing individual responses were retested in at least two independent transfections. Those compounds were subsequently subjected to dose-response curves, conducted in at least three independent transfections. For these assays, the highest concentration was 1 mM (1×10⁻³ M), and the lowest was 1 pM (1×10⁻¹² M). HBSS was used as the negative control and for data normalisation. For dose-response curves, responses were normalised to the maximum response obtained following ligand addition in each experiment (100% activation) and to the response obtained with the vehicle (0% activation). Normalised data from independent transfections were used to plot dose–response curves, which were fitted using a four-parameter logistic model with the drc package in R (Ritz et al. 2015).

The analysis of human melatonin receptors was performed after identifying the orthology between placozoan monoaminergic receptors and human melatonin receptors. The same method described above was used, with the key difference that mixture assays were not necessary, as melatonin receptors are already well-established monoamine receptors. Instead, only individual compounds were tested at high concentrations 100uM, followed by dose-response curve analysis. For this assay, human melatonin A (hMelA; MTNR1A Human) and human melatonin B (hMelB; MTNR1B) receptors were obtained from the Addgene PRESTO-Tango GPCR Kit from the Roth laboratory (Kroeze et al. 2015). The raw data used for these dose-response analyses and the scripts used for the data analyses are available on GitHub (https://github.com/Imnotabioinformatician/MonoamineData (available after publication)

### Phylogenetic analysis of monoamine GPCRs

For the phylogenetic analysis, we extracted the sequences of the bilaterian and non-bilaterian monoaminergic GPCR cluster and those receptors that showed any connection to the “monoamine-cluster” (Fig. 1). As an outgroup, we chose a sub-cluster of neuropeptide receptors that showed the strongest connection to the main monoaminergic GPCR cluster. We aligned the extracted genes with MAFFT v7 using the iterative refinement method E-INS-i (Katoh et al. 2002) Alignments were trimmed with TrimAl in gappy-out mode, and maximum likelihood trees were calculated with IQ-TREE2 using the LG + G4 model (Minh et al. 2020) Branch support was calculated by running 1,000 replicates with the aLRT-SH-like and ultrafast bootstrap methods. Protein sequences and trimmed alignments files are provided in supplementary files 5 and 6. The final tree visualization was carried out using FigTree. The tree was rooted using neuropeptide receptors as root.

### Phylogenetic analysis of AADC enzymes

To identify potential aromatic amino acid decarboxylase (AADC) enzymes in non-bilaterians, we first selected a set of known AADC enzymes to serve as our reference dataset. Using this dataset, we generated a Hidden Markov Model (HMM) using HMMER (v3.1b2) and applied it to search the proteomes of the same range of bilaterian, non-bilaterian, and single-celled eukaryotic species as described previously for the GPCR identification. An e-value threshold of 1e−10 was used for the search. Following the identification of potential homologues, redundant sequences were removed using CD-Hit with a 95% similarity threshold. Due to the relatively small number of AADC homologues identified through this process, a cluster-based analysis using CLANS was deemed unnecessary and was not performed. Instead, we proceeded directly to phylogenetic analysis. For this analysis, we extracted the sequences of all identified bilaterian and non-bilaterian AADC homologues. As specified, glutamate decarboxylase (GAD) sequences were chosen and included as the outgroup. We aligned the extracted sequences with MAFFT v7 using the iterative refinement method E-INS-i (Katoh et al. 2002). The alignments were subsequently trimmed with TrimAl in gappy-out mode. Maximum likelihood trees were calculated using IQ-TREE2 employing the LG + G4 model (Minh et al. 2020). Branch support was calculated by running 1,000 replicates with both the aLRT-SH-like and ultrafast bootstrap methods. The final tree visualization was carried out using FigTree. The protein sequences used for the AADC phylogeny are available in supplementary file 7. Trimmed alignment file is provided in supplementary file 8.

### Structural analysis and protein sequence alignments

The structures of human MelA, Tadh170, and Tadh173 were predicted with Alphafold3. All predicted structures were structurally aligned to the solved structure of MelA bound by 2-iodomelatonin (PDB number 6ME4) using the program ChimeraX version 1.10. All predictions scores for Alphafold, and structural alignment scores from ChimeraX, are described in the text. The protein alignment was generated with the program MUSCLE and annotated with Jalview. Both the structural images exported from ChimeraX and the Jalview protein alignments were further annotated in Adobe Illustrator.

### Single cell sequencing analysis

Single-cell atlas datasets for *Trichoplax adhaerens*, *Trichoplax H2*, *Hoilungia hongkongensis*, and *Cladtertia collaboinventa* were obtained from Najle et al. (2023). Expression data for the deorphanized receptors Tadh153, Tadh165, Tadh168, Tadh170, Tadh173, the AADC enzyme and their homologous sequences from the other species were extracted from these datasets, along with the associated cell clustering information. To identify top cellular expression patterns for the various placozoan receptors, single-cell RNA-Seq UMI-frac values for each receptor transcript were analysed, and the sum value for each metacell type was determined.

### Behavioural analysis of monoamines in *Trichoplax adhaerens in vivo*

*Trichoplax adhaerens* (Schulze, 1883) culture was maintained in artificial seawater (Instant Ocean) at room temperature and 35ppt. The water was partially changed on a weekly basis, and a suspension of green algae (*Dunaliella tertiolecta* Butcher, 1959) was added to feed the animals. The algae were grown in ASW supplemented with 0.8% Micro Algae Grow (Florida Aqua Farms) under constant light and aeration.

24 hours prior each experiment, a group of 6-14 Trichoplax was transferred to ASW with no algae. To conduct the experiment, these animals were further transferred to a new Petri dish with 10 ml of filter-sterilized ASW. Individuals were placed well apart from each other so that they do not come into contact with each other during the recording. Once animals adhered, the dish was placed under the camera (Canon EOS Rebel T5) and the images were taken every 30 seconds for three hours. In 1.5 hours after the beginning of recording, 1 ml of ASW was taken out from the dish with Trichoplax, mixed with an aliquot of stock solution of a monoamine or vehicle (control), and gently returned to the dish in a dropwise manner. In this way we tested three monoamines: Tyramine (ThermoScientific, A12220.06), phenethylamine (PEA; TCI, P0086), and Tryptamine (TPA, TCI, A0300). Stock solutions contained 100 mM of a monoamine, 50% DMSO, and 200 µM ascorbic acid; control was with no monoamine. The final working concentration of monoamines in the dish with animals was 50 µM; the concentration was lowered to 10 or 1 µM if the effect of initial concentration (50 µM) elicited a strong response. Each concentration was tested in at least three independent experiments.

Once the recording was over, the images were opened as a stack in the Fiji software, and each *Trichoplax* individual (except those who left field of view, divided, or got into contact with a teammate during recording) was examined with the Analyze Particles command, as per (Heyland et al. 2014). The scored measurements were area and perimeter of the animal, its X and Y coordinates, and shape description. The dynamic of each parameter was evaluated by plotting it against time; in this way we selected a time frame, when the effect of an applied monoamine was apparent (60 minutes for tyramine and 30 minutes for phenethylamine and tryptamine), and further analysed the values obtained for this period of time before (pre-test) and after (post-test) the application in both experimental and control groups. Since Trichoplax dynamically changes its body shape (which includes both changes in the area and body outline), we averaged area and roundness values of each individual obtained during pre- and post-test measurements at each time point. X and Y coordinates were used to calculate distance travelled by the animal and to recreate its track using MotilityLab online resource (Wortel et al. 2021). To infer about the velocity, we measured total pre- and post-test paths for each animal by summarizing the distances the animal travelled between each time points.

We chose an ANCOVA test in order to compare post-test values between experimental and control groups taking into account pre-test values (as a covariate), which largely correlate with the post-test values. Real Statistics resource package for Excel was used to ensure that ANCOVA assumptions were met, as well as to run an ANCOVA test itself. For multiple comparisons (when more than one concentration of a monoamine was tested), Tukey-Kramer post-hoc test with the Bonferroni correction was applied.

## Supporting information

SupplementaryFile7.txt

SupplementaryFile8.txt

## Funding

L.A.Y.G. was supported by a BBSRC fellowship (BB/W010305/2) and funding from the Royal Society (RG\R1\241397). A.S and T.D.M., were funded by an NSERC Discovery Grant (RGPIN-2021-03557) and an NSERC Discovery Accelerator Supplement (RGPAS-2021-00002) to A.S.

## Contributions

LAYG and GJ conceived the project. AS and LAYG conducted the bionformatic analyses. LAYG, LG, HW, and TM performed the experimental work. LAYG wrote the first draft of the manuscript, and all authors contributed to revisions.

## Acknowledgements

LAYG would like to thank Dr. Bryan Roth for his suggestions regarding human deorphanisation methodologies. Additionally, the open resources he provided at Addgene have enabled my recently established laboratory to conduct experiments that would otherwise not be possible.

## Figures

**Supplementary Figure 1.**
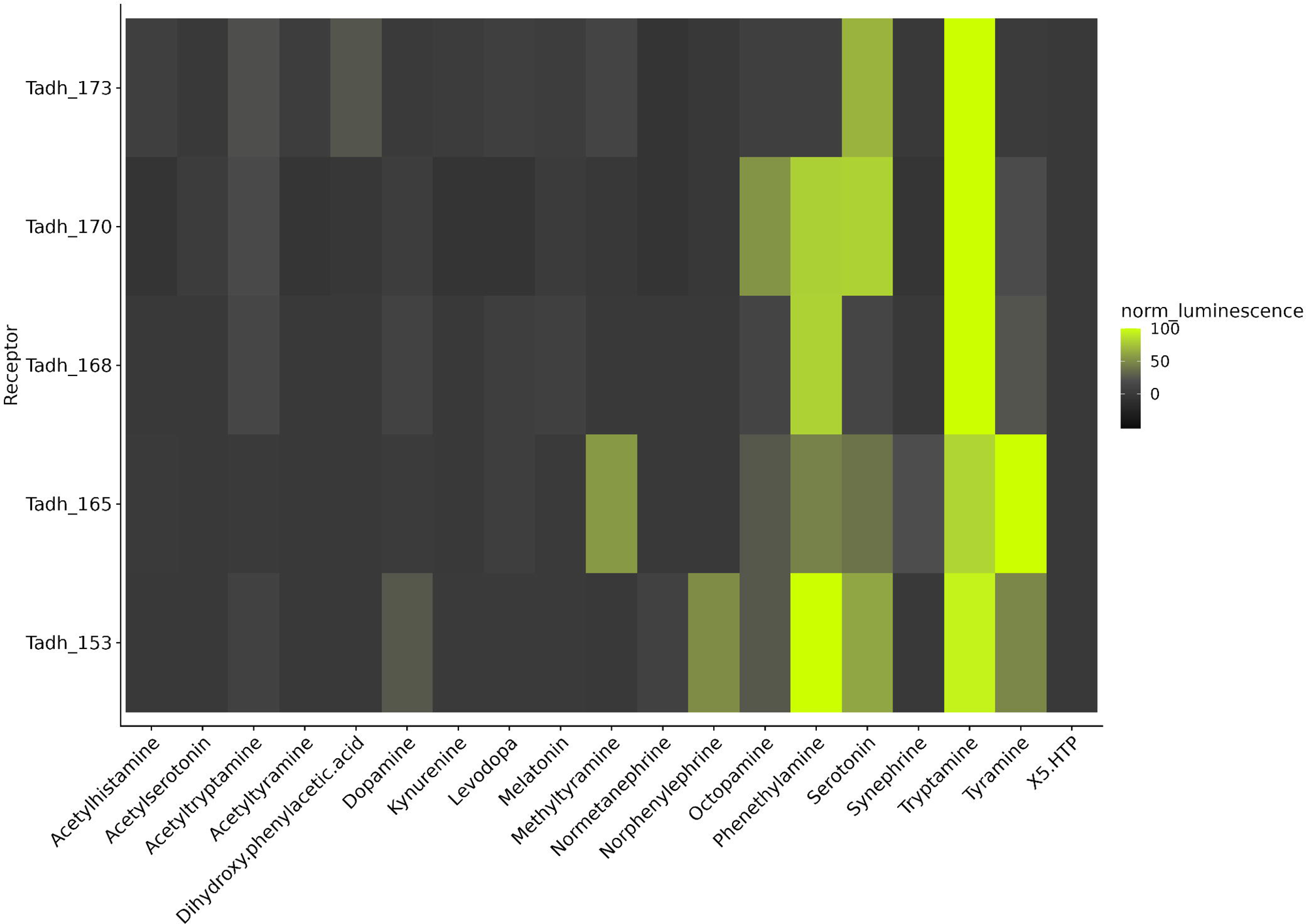
Heatmap of receptor responses to individual compounds identified from active peptide mixes. Relative luminescence values from at least two replicates per receptor in response to individual compounds derived from the active mixes that elicited signals in placozoan receptors is shown. Colour intensity corresponds to luminescence levels, as indicated by the scale bar. Responses were normalised to the negative control (0%) and to five times the negative control (100%) to facilitate the identification of potential signals.

**Supplementary Figure 2.**
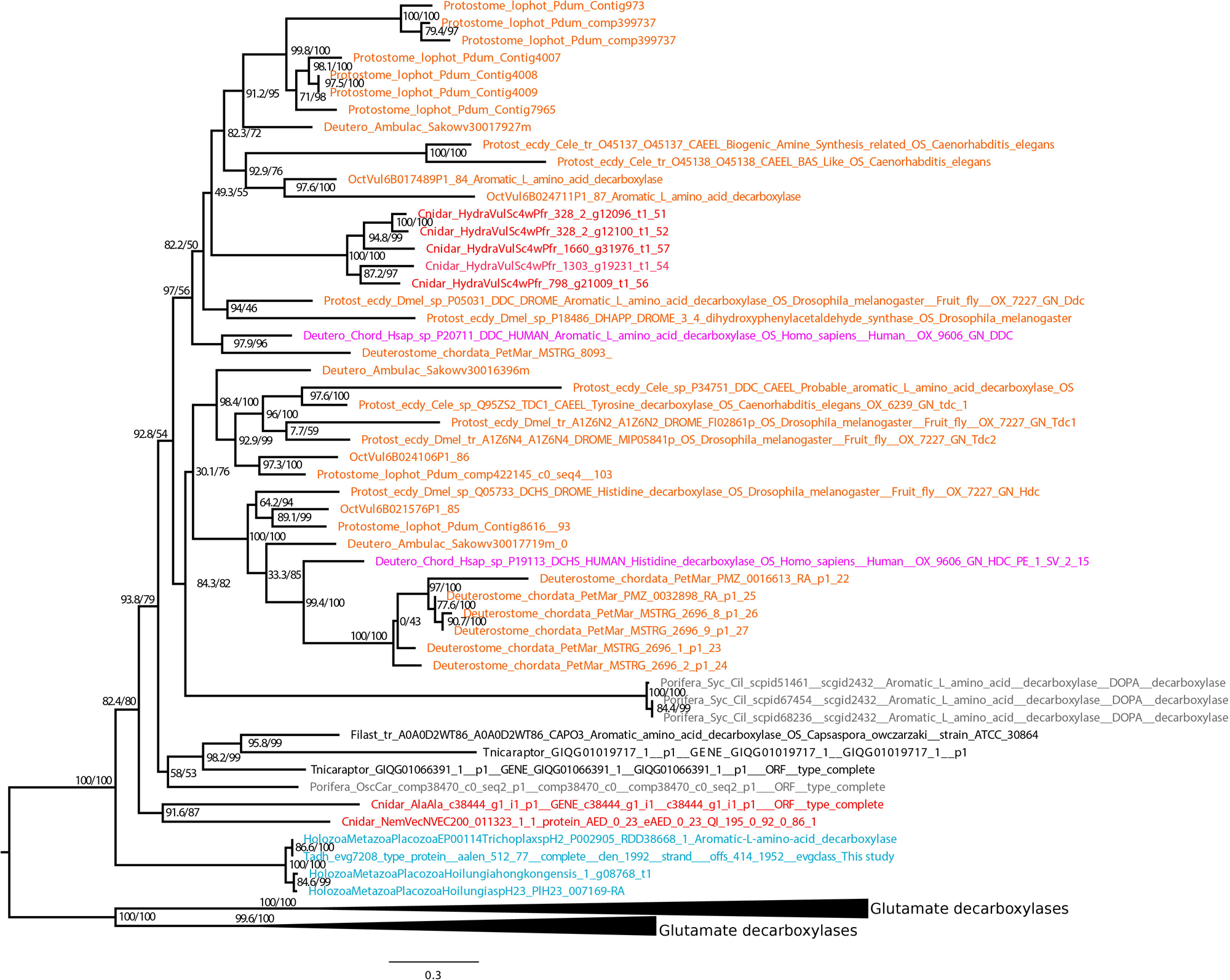
Maximum likelihood phylogeny of aromatic amino acid decarboxylase (AADC) enzymes. Maximum likelihood phylogeny of AADC enzymes generated using IQ-TREE2 with the best-fit model LG+G4. The tree was rooted with glutamate decarboxylases, which are collapsed for clarity. Node support values (shown as numbers) are based on 1000 replicates, representing SH-aLRT and ultrafast bootstrap support. Taxa are color-coded as follows: red for cnidarians, blue for placozoans, grey for poriferans, and black for unicellular eukaryotes. Bilaterians are indicated in orange for bilaterian-specific clades, except for humans that are shown in pink.

**Supplementary Figure 3.**
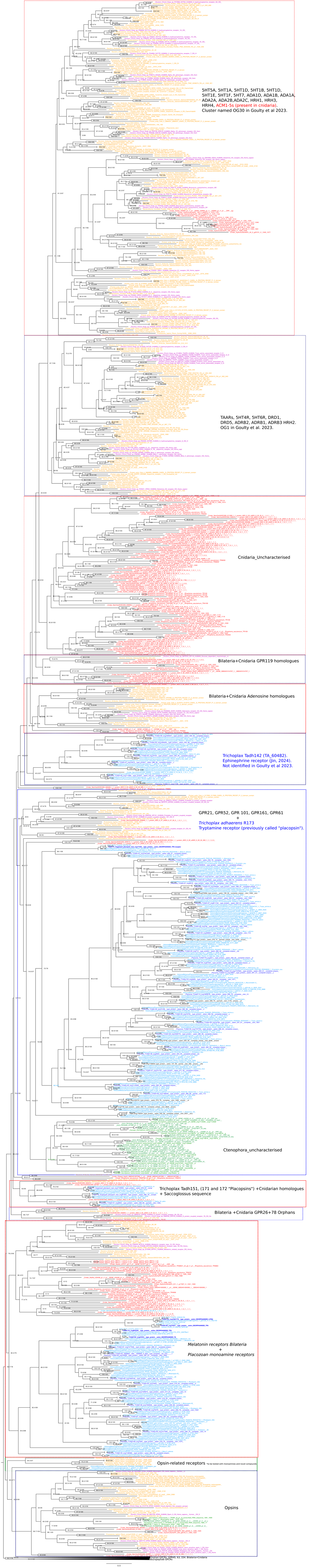
Maximum likelihood phylogeny of monoamine GPCRs. The phylogeny was generated using IQ-TREE2 with the best-fit model LG+G4. The tree is rooted with neuropeptide receptors from both bilaterians and non-bilaterians, which are collapsed for clarity. Node support values (shown as numbers) are based on 1000 replicates and represent SH-aLRT and ultrafast bootstrap support. Taxa are colour-coded as follows: red for cnidarians, blue for placozoans, green for ctenophores, and grey for poriferans. Bilaterians are shown in orange, with humans highlighted in pink. This tree is the non-collapsed version of the tree presented in the figure 4A.

**Supplementary Figure 4.**
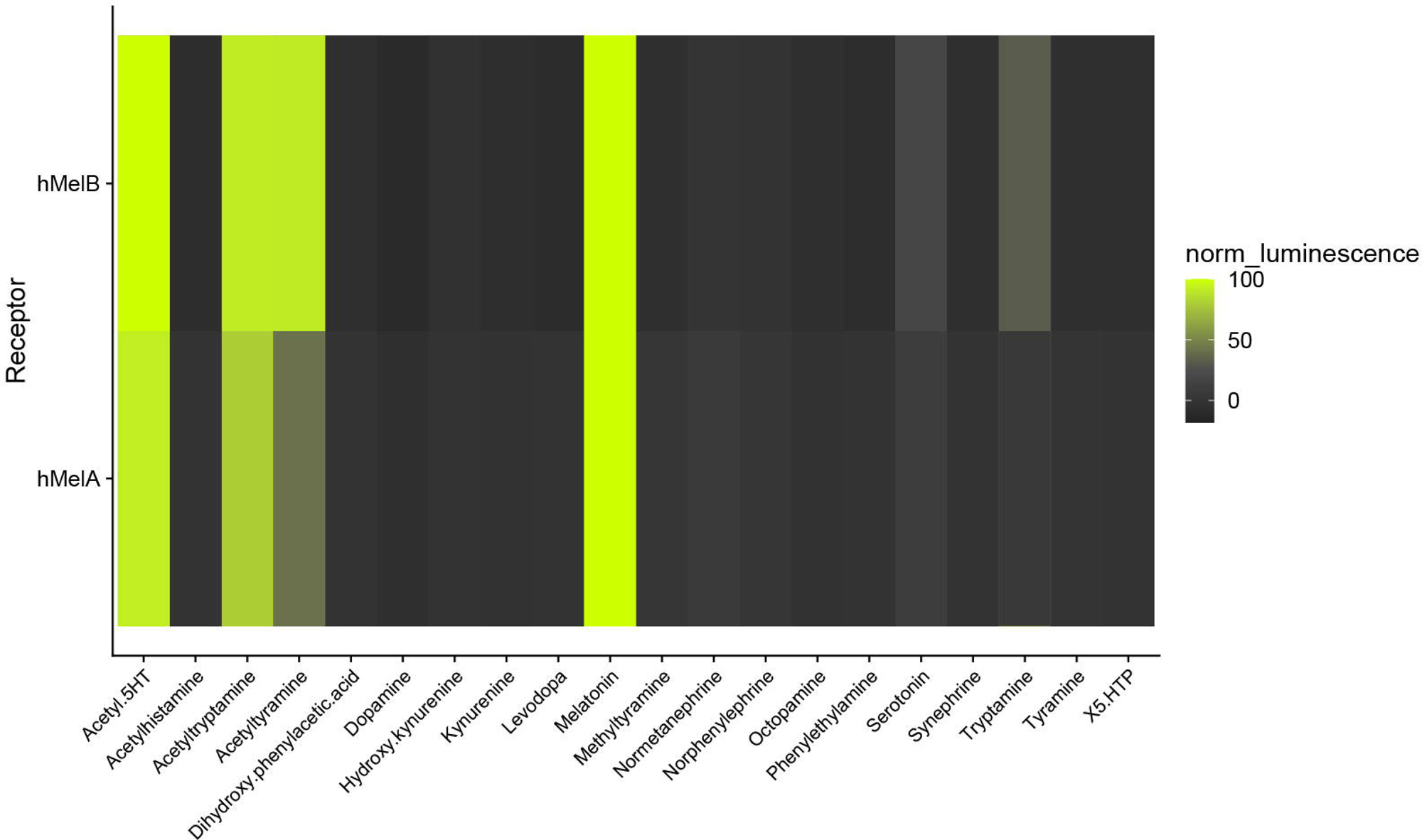
Heatmap of human melatonin receptor responses to individual compounds. Relative luminescence values from at least two replicates per receptor in response to individual compounds are shown. These compounds were initially identified from active peptide mixes that elicited signals in placozoan receptors, but were tested individually on the human melatonin receptors. Colour intensity corresponds to luminescence levels, as indicated by the scale bar. Responses were normalised to the negative control (0%) and to five times the negative control (100%) to facilitate the identification of potential signals.

**Supplementary Figure 5.**
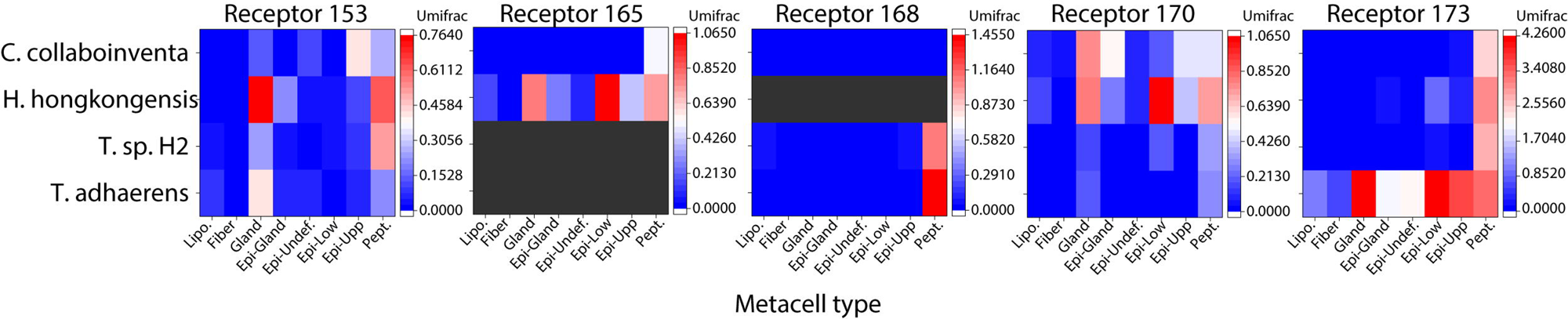
Heat map plots of sum Unifrac expression of the *T. adhaerens* receptors Tadh153, Tadh165, Tadh168, Tadh170, Tadh173 and their homologues from three other placozoan species within different metacell types extracted from the available scRNA-seq data. The key to the right of each graph shows relative expression levels in different clusters, and the boxes in black indicate an absence of data for a given receptor/species.

## Notes

### Competing Interest Statement

The authors have declared no competing interest.

## Bibliography

Atwood, Brady K., Jacqueline Lopez, James Wager-Miller, Ken Mackie, and Alex Straiker. 2011. “Expression of G Protein-Coupled Receptors and Related Proteins in HEK293, AtT20, BV2, and N18 Cell Lines as Revealed by Microarray Analysis.” BMC Genomics 12 (January): 14.

Bacqué-Cazenave, Julien, Rahul Bharatiya, Grégory Barrière, Jean-Paul Delbecque, Nouhaila Bouguiyoud, Giuseppe Di Giovanni, Daniel Cattaert, and Philippe De Deurwaerdère. 2020. “Serotonin in Animal Cognition and Behavior.” International Journal of Molecular Sciences 21 (5). 10.3390/ijms21051649.

Bauknecht, Philipp, and Gáspár Jékely. 2017. “Ancient Coexistence of Norepinephrine, Tyramine, and Octopamine Signaling in Bilaterians.” BMC Biology 15 (1): 6.

Bayliss, Asha, Giuliana Roselli, and Peter D. Evans. 2013. “A Comparison of the Signalling Properties of Two Tyramine Receptors from Drosophila.” Journal of Neurochemistry 125 (1): 37–48.

Berry, M. D., A. V. Juorio, X. M. Li, and A. A. Boulton. 1996. “Aromatic L-Amino Acid Decarboxylase: A Neglected and Misunderstood Enzyme.” Neurochemical Research 21 (9): 1075–87.

Berry, Mark D. 2004. “Mammalian Central Nervous System Trace Amines. Pharmacologic Amphetamines, Physiologic Neuromodulators.” Journal of Neurochemistry 90 (2): 257–71.

Berry, Mark D., Shannon Hart, Anthony R. Pryor, Samantha Hunter, and Danielle Gardiner. 2016. “Pharmacological Characterization of a High-Affinity p-Tyramine Transporter in Rat Brain Synaptosomes.” Scientific Reports 6 (November): 38006.

Berry, Mark D., Mithila R. Shitut, Ahmed Almousa, Jane Alcorn, and Bruno Tomberli. 2013. “Membrane Permeability of Trace Amines: Evidence for a Regulated, Activity-Dependent, Nonexocytotic, Synaptic Release.” Synapse (New York, N.Y.) 67 (10): 656–67.

Blumenthal, Edward M. 2003. “Regulation of Chloride Permeability by Endogenously Produced Tyramine in the Drosophila Malpighian Tubule.” American Journal of Physiology. Cell Physiology 284 (3): C718–28.

Branicky, Robyn, and Wiliam R. Schafer. 2009. “Tyramine: A New Receptor and a New Role at the Synapse.” Neuron.

Cazzamali, Giuseppe, Dan A. Klaerke, and Cornelis J. P. Grimmelikhuijzen. 2005. “A New Family of Insect Tyramine Receptors.” Biochemical and Biophysical Research Communications 338 (2): 1189–96.

Chen, Haiwei, Connor E. Rosen, Jaime A. González-Hernández, Deguang Song, Jan Potempa, Aaron M. Ring, and Noah W. Palm. 2023. “Highly Multiplexed Bioactivity Screening Reveals Human and Microbiota Metabolome-GPCRome Interactions.” Cell 186 (14): 3095–3110.e19.

Cheng, Zhijie, Denise Garvin, Aileen Paguio, Pete Stecha, Keith Wood, and Frank Fan. 2010. “Luciferase Reporter Assay System for Deciphering GPCR Pathways.” Current Chemical Genomics 4 (December): 84–91.

Clark, Tobias, Vera Hapiak, Mitchell Oakes, Holly Mills, and Richard Komuniecki. 2018. “Monoamines Differentially Modulate Neuropeptide Release from Distinct Sites within a Single Neuron Pair.” PloS One 13 (5): e0196954.

Claustrat, B., and J. Leston. 2015. “Melatonin: Physiological Effects in Humans.” Neuro-Chirurgie 61 (2–3): 77–84.

Cullum, Sean A., Dmitry B. Veprintsev, and Stephen J. Hill. 2023. “Kinetic Analysis of Endogenous β -Adrenoceptor-Mediated CAMP GloSensor^TM^ Responses in HEK293 Cells.” British Journal of Pharmacology 180 (10): 1304–15.

Daubner, S. Colette, Tiffany Le, and Shanzhi Wang. 2011. “Tyrosine Hydroxylase and Regulation of Dopamine Synthesis.” Archives of Biochemistry and Biophysics 508 (1): 1–12.

Dubocovich, M. L., M. I. Masana, S. Iacob, and D. M. Sauri. 1997. “Melatonin Receptor Antagonists That Differentiate between the Human Mel1a and Mel1b Recombinant Subtypes Are Used to Assess the Pharmacological Profile of the Rabbit Retina ML1 Presynaptic Heteroreceptor.” Naunyn-Schmiedeberg’s Archives of Pharmacology 355 (3): 365–75.

Durocher, Yves, Sylvie Perret, and Amine Kamen. 2002. “High-Level and High-Throughput Recombinant Protein Production by Transient Transfection of Suspension-Growing Human 293-EBNA1 Cells.” Nucleic Acids Research 30 (2): E9.

Eiden, Lee E., and Eberhard Weihe. 2011. “VMAT2: A Dynamic Regulator of Brain Monoaminergic Neuronal Function Interacting with Drugs of Abuse.” Annals of the New York Academy of Sciences 1216 (January): 86–98.

Feuda, Roberto, Sinead C. Hamilton, James O. McInerney, and Davide Pisani. 2012. “Metazoan Opsin Evolution Reveals a Simple Route to Animal Vision.” Proceedings of the National Academy of Sciences of the United States of America 109 (46): 18868–72.

Francis, Warren R., Michael Eitel, Sergio Vargas, Marcin Adamski, Steven H. D. Haddock, Stefan Krebs, Helmut Blum, Dirk Erpenbeck, and Gert Wörheide. 2017. “The Genome of the Contractile Demosponge Tethya Wilhelma and the Evolution of Metazoan Neural Signalling Pathways.” Cold Spring Harbor Laboratory. 10.1101/120998.

Fussnecker, Brendon L., Brian H. Smith, and Julie A. Mustard. 2006. “Octopamine and Tyramine Influence the Behavioral Profile of Locomotor Activity in the Honey Bee (Apis Mellifera).” Journal of Insect Physiology 52 (10): 1083–92.

Goulty, Matthew, Gaelle Botton-Amiot, Ezio Rosato, Simon G. Sprecher, and Roberto Feuda. 2023. “The Monoaminergic System Is a Bilaterian Innovation.” Nature Communications 14 (1): 3284.

Hajj-Ali, I., and M. Anctil. 1997. “Characterization of a Serotonin Receptor in the Cnidarian Renilla Koellikeri: A Radiobinding Analysis.” Neurochemistry International 31 (1): 83–93.

Hapiak, Vera, Philip Summers, Amanda Ortega, Wen Jing Law, Andrew Stein, and Richard Komuniecki. 2013. “Neuropeptides Amplify and Focus the Monoaminergic Inhibition of Nociception in Caenorhabditis Elegans.” The Journal of Neuroscience : The Official Journal of the Society for Neuroscience 33 (35): 14107–16.

Hauser, Frank, Giuseppe Cazzamali, Michael Williamson, Wolfgang Blenau, and Cornelis J. P. Grimmelikhuijzen. 2006. “A Review of Neurohormone GPCRs Present in the Fruitfly Drosophila Melanogaster and the Honey Bee Apis Mellifera.” Progress in Neurobiology 80 (1): 1–19.

Heyland, Andreas, Roger Croll, Sophie Goodall, Jeff Kranyak, and Russell Wyeth. 2014. “Trichoplax Adhaerens, an Enigmatic Basal Metazoan with Potential.” *Methods in Molecular Biology (Clifton*, N.J.) 1128: 45–61.

Hiasa, Miki, Takaaki Miyaji, Yuka Haruna, Tomoya Takeuchi, Yuika Harada, Sawako Moriyama, Akitsugu Yamamoto, Hiroshi Omote, and Yoshinori Moriyama. 2014. “Identification of a Mammalian Vesicular Polyamine Transporter.” Scientific Reports 4 (October): 6836.

Horie, Takeo, Masashi Nakagawa, Yasunori Sasakura, and Takehiro G. Kusakabe. 2009. “Cell Type and Function of Neurons in the Ascidian Nervous System.” Development, Growth & Differentiation 51 (3): 207–20.

Jin, Minjun, Wanqing Li, Zhongyu Ji, Guotao Di, Meng Yuan, Yifan Zhang, Yunsi Kang, and Chengtian Zhao. 2024. “Coordinated Cellular Behavior Regulated by Epinephrine Neurotransmitters in the Nerveless Placozoa.” Nature Communications 15 (1): 8626.

Johansson, L. C., B. Stauch, J. D. McCorvy, G. W. Han, N. Patel, X. P. Huang, A. Batyuk, et al. 2019. “XFEL Structures of the Human MT2 Melatonin Receptor Reveal the Basis of Subtype Selectivity.” Nature 569 (7755). 10.1038/s41586-019-1144-0.

Käll, Lukas, Anders Krogh, and Erik L. L. Sonnhammer. 2007. “Advantages of Combined Transmembrane Topology and Signal Peptide Prediction--the Phobius Web Server.” Nucleic Acids Research 35 (Web Server issue): W429–32.

Katoh, Kazutaka, Kazuharu Misawa, Kei-Ichi Kuma, and Takashi Miyata. 2002. “MAFFT: A Novel Method for Rapid Multiple Sequence Alignment Based on Fast Fourier Transform.” Nucleic Acids Research 30 (14): 3059–66.

Komuniecki, Rick, Gareth Harris, Vera Hapiak, Rachel Wragg, and Bruce Bamber. 2012. “Monoamines Activate Neuropeptide Signaling Cascades to Modulate Nociception in C. Elegans: A Useful Model for the Modulation of Chronic Pain?” Invertebrate Neuroscience : IN 12 (1): 53–61.

Kosa, E., A. Marcilhac-Flouriot, M. P. Fache, and P. Siaud. 2000. “Effects of Beta-Phenylethylamine on the Hypothalamo-Pituitary-Adrenal Axis in the Male Rat.” Pharmacology, Biochemistry, and Behavior 67 (3): 527–35.

Kroeze, Wesley K., Maria F. Sassano, Xi-Ping Huang, Katherine Lansu, John D. McCorvy, Patrick M. Giguère, Noah Sciaky, and Bryan L. Roth. 2015. “PRESTO-Tango as an Open-Source Resource for Interrogation of the Druggable Human GPCRome.” Nature Structural & Molecular Biology 22 (5): 362–69.

Ma, Zongyuan, Xiaojiao Guo, Hong Lei, Ting Li, Shuguang Hao, and Le Kang. 2015. “Octopamine and Tyramine Respectively Regulate Attractive and Repulsive Behavior in Locust Phase Changes.” Scientific Reports 5 (January): 8036.

McCauley, D. W. 1997. “Serotonin Plays an Early Role in the Metamorphosis of the Hydrozoan Phialidium Gregarium.” Developmental Biology 190 (2): 229–40.

Meng, Da, Hui-Quan Li, Karl Deisseroth, Stefan Leutgeb, and Nicholas C. Spitzer. 2018. “Neuronal Activity Regulates Neurotransmitter Switching in the Adult Brain Following Light-Induced Stress.” Proceedings of the National Academy of Sciences of the United States of America 115 (20): 5064–71.

Mills, Holly, Rachel Wragg, Vera Hapiak, Michelle Castelletto, Jeffrey Zahratka, Gareth Harris, Philip Summers, et al. 2012. “Monoamines and Neuropeptides Interact to Inhibit Aversive Behaviour in Caenorhabditis Elegans.” The EMBO Journal 31 (3): 667–78.

Minh, Bui Quang, Heiko A. Schmidt, Olga Chernomor, Dominik Schrempf, Michael D. Woodhams, Arndt von Haeseler, and Robert Lanfear. 2020. “IQ-TREE 2: New Models and Efficient Methods for Phylogenetic Inference in the Genomic Era.” Molecular Biology and Evolution 37 (5): 1530–34.

Moeller, Mareen, Samuel Nietzer, and Peter J. Schupp. 2019. “Neuroactive Compounds Induce Larval Settlement in the Scleractinian Coral Leptastrea Purpurea.” Scientific Reports 9 (1): 2291.

Moriyama, Yoshinori, Ryo Hatano, Satomi Moriyama, and Shunsuke Uehara. 2020. “Vesicular Polyamine Transporter as a Novel Player in Amine-Mediated Chemical Transmission.” Biochimica et Biophysica Acta. Biomembranes 1862 (12): 183208.

Moroz, Leonid L., Daria Y. Romanova, and Andrea B. Kohn. 2021. “Neural versus Alternative Integrative Systems: Molecular Insights into Origins of Neurotransmitters.” Philosophical Transactions of the Royal Society of London. Series B, Biological Sciences 376 (1821): 20190762.

Najle, Sebastián R., Xavier Grau-Bové, Anamaria Elek, Cristina Navarrete, Damiano Cianferoni, Cristina Chiva, Didac Cañas-Armenteros, et al. 2023. “Stepwise Emergence of the Neuronal Gene Expression Program in Early Animal Evolution.” Cell 186 (21): 4676–4693.e29.

Nakagawa, Hiroyuki, Shiori Maehara, Kazuhiko Kume, Hiroto Ohta, and Jun Tomita. 2022. “Biological Functions of Α2-Adrenergic-like Octopamine Receptor in Drosophila Melanogaster.” Genes, Brain, and Behavior 21 (6): e12807.

Naoi, Makoto, Wakako Maruyama, and Masayo Shamoto-Nagai. 2018. “Type A Monoamine Oxidase and Serotonin Are Coordinately Involved in Depressive Disorders: From Neurotransmitter Imbalance to Impaired Neurogenesis.” Journal of Neural Transmission 125 (1): 53–66.

Oldendorf, W. H. 1971. “Brain Uptake of Radiolabeled Amino Acids, Amines, and Hexoses after Arterial Injection.” The American Journal of Physiology 221 (6): 1629–39.

Patel, Nilkanth, Xi Ping Huang, Jessica M. Grandner, Linda C. Johansson, Benjamin Stauch, John D. McCorvy, Yongfeng Liu, Bryan Roth, and Vsevolod Katritch. 2020. “Structure-Based Discovery of Potent and Selective Melatonin Receptor Agonists.” ELife 9 (March). 10.7554/eLife.53779.

Pauls, Dennis, Christine Blechschmidt, Felix Frantzmann, Basil El Jundi, and Mareike Selcho. 2018. “A Comprehensive Anatomical Map of the Peripheral Octopaminergic/Tyraminergic System of Drosophila Melanogaster.” Scientific Reports 8 (1): 15314.

Rovati, G. Enrico, Valérie Capra, and Richard R. Neubig. 2007. “The Highly Conserved DRY Motif of Class A G Protein-Coupled Receptors: Beyond the Ground State.” Molecular Pharmacology 71 (4): 959–64.

Sinakevitch, Irina T., Gabriella H. Wolff, Hans-Joachim Pflueger, and Brian H. Smith. 2018. Biogenic Amines and Neuromodulation of Animal Behavior, 2nd *Edition*. Frontiers Media SA.

Speight, G., D. Turic, J. Austin, B. Hoogendoorn, A. G. Cardno, L. Jones, K. C. Murphy, et al. 2000. “Comparative Sequencing and Association Studies of Aromatic L-Amino Acid Decarboxylase in Schizophrenia and Bipolar Disorder.” Molecular Psychiatry 5 (3): 327–31.

Stauch, Benjamin, Linda C. Johansson, John D. McCorvy, Nilkanth Patel, Gye Won Han, Xi-Ping Huang, Cornelius Gati, et al. 2019. “Structural Basis of Ligand Recognition at the Human MT Melatonin Receptor.” Nature 569 (7755): 284– 88.

Takeuchi, Tomoya, Yuika Harada, Satomi Moriyama, Kazuyuki Furuta, Satoshi Tanaka, Takaaki Miyaji, Hiroshi Omote, Yoshinori Moriyama, and Miki Hiasa. 2017. “Vesicular Polyamine Transporter Mediates Vesicular Storage and Release of Polyamine from Mast Cells.” The Journal of Biological Chemistry 292 (9): 3909–18.

Thiel, Daniel, Luis Alfonso Yañez Guerra, Amanda Kieswetter, Alison G. Cole, Liesbet Temmerman, Ulrich Technau, and Gáspár Jékely. 2024. “Large-Scale Deorphanization of Nematostella Vectensis Neuropeptide G Protein-Coupled Receptors Supports the Independent Expansion of Bilaterian and Cnidarian Peptidergic Systems.” ELife 12 (May). 10.7554/elife.90674.3.

Thiel, Daniel, Luis Alfonso Yañez-Guerra, and Gaspar Jekely. 2023. “GPCR Deorphanization Assay in HEK-293 Cells.” Bio-Protocol. 10.21769/p2493.

Wang, Qinggong, Qiuyuan Lu, Qiong Guo, Maikun Teng, Qingguo Gong, Xu Li, Yang Du, Zheng Liu, and Yuyong Tao. 2022. “Structural Basis of the Ligand Binding and Signaling Mechanism of Melatonin Receptors.” Nature Communications 13 (1): 454.

Wortel, Inge M. N., Annie Y. Liu, Katharina Dannenberg, Jeffrey C. Berry, Mark J. Miller, and Johannes Textor. 2021. “CelltrackR: An R Package for Fast and Flexible Analysis of Immune Cell Migration Data.” Immunoinformatics (Amsterdam, Netherlands) 1–2 (100003): 100003.

Xiang, Xueyan, Arturo A. Vilar Gomez, Simone P. Blomberg, Huifang Yuan, Bernard M. Degnan, and Sandie M. Degnan. 2023. “Potential for Host-Symbiont Communication via Neurotransmitters and Neuromodulators in an Aneural Animal, the Marine Sponge.” Frontiers in Neural Circuits 17 (September): 1250694.

Xie, Zhihua, Susan V. Westmoreland, and Gregory M. Miller. 2008. “Modulation of Monoamine Transporters by Common Biogenic Amines via Trace Amine-Associated Receptor 1 and Monoamine Autoreceptors in Human Embryonic Kidney 293 Cells and Brain Synaptosomes.” The Journal of Pharmacology and Experimental Therapeutics 325 (2): 629–40.

Yamamoto, Kei, and Philippe Vernier. 2011. “The Evolution of Dopamine Systems in Chordates.” Frontiers in Neuroanatomy 5 (March): 21.

Yoshida, Midori, Eitaro Oami, Min Wang, Shoichi Ishiura, and Satoshi Suo. 2014. “Nonredundant Function of Two Highly Homologous Octopamine Receptors in Food-Deprivation-Mediated Signaling in Caenorhabditis Elegans.” Journal of Neuroscience Research 92 (5): 671–78.

Zega, Giuliana, Roberta Pennati, Arianna Fanzago, and Fiorenza De Bernardi. 2007. “Serotonin Involvement in the Metamorphosis of the Hydroid Eudendrium Racemosum.” The International Journal of Developmental Biology 51 (4): 307–13.

Zhang, Haixia, Lin Wang, Kun Shi, Dongqian Shan, Yunpeng Zhu, Chanyu Wang, Yixue Bai, Tianci Yan, Xiaodong Zheng, and Jin Kong. 2019. “Apple Tree Flowering Is Mediated by Low Level of Melatonin under the Regulation of Seasonal Light Signal.” Journal of Pineal Research 66 (2): e12551.

Zhang, Mengliang. 2016. “Two-Step Production of Monoamines in Monoenzymatic Cells in the Spinal Cord: A Different Control Strategy of Neurotransmitter Supply?” Neural Regeneration Research 11 (12): 1904–9.

Zhu, Yuling, Taiwei Yang, Yueyang Chen, Cong Fan, and Jifeng Yuan. 2020. “One-Pot Synthesis of Aromatic Amines from Renewable Feedstocks via Whole-Cell Biocatalysis.” ChemistrySelect 5 (45): 14292–95.

Zisapel, N. 2001. “Circadian Rhythm Sleep Disorders: Pathophysiology and Potential Approaches to Management.” CNS Drugs 15 (4): 311–28.

