## SupplementaryFile1 for "Functional and phylogenetic analysis of placozoan GPCRs reveal the prebilaterian origin of monoaminergic signalling"

>Placozoa_TriAdh111_evg17738 type=protein_ aalen=334_51%_complete-utrpoor_ clen=1946_ strand=| offs=292-1296_ evgclass=main_okay_match_evg36580_pct_99_100_-sense_ 6697

MNLTVTVQTFSFIVSMTTILLAAVSIVANTCILIIFFNVRRLRSTTNCLVANLAASDLCFSIFAIPCQLFERFLPESEAICLLNSISTIFFTTSSIFSLCFIGIIKYINITKPLKNNLIITRTRAYKAIVFIWLFSSLLTFVPVIFITHRKLLCHGFKYRTGKLLDKFYFLTVFVITYLIPTVILIYVYTVIYKIARRQCSRIRRSIKSIHRNTNEIRSSLPSSKSATFLLALVTCFIVCYTPFFCSELIHRFIAPISNNVSLYITICFITANSAINPFLYGILKPEFKYNIQFYFRLSLCKDDGNITHSSSYSQRLPFQRSPSQVSTTNPRYV

>Placozoa_TriAdh112_evg100509 type=protein_ aalen=368_37%_complete-utrbad_ clen=2958_ strand=-_ offs=2638-1532_ evgclass=maina2_okay_match_evg869784_pct_99_100_._ 6433

MIDLTTTNVSTEATTLDQWQPSSAFIYYFIPGAVIIMIVSGLGNTFCLTIVCRTSLKQDVDTLFIANLAITDLISALLVLPSIIVTNMCYVYNQLSQGLCNFQGFLLIWIMGTSLFTTALISIDRCIAVTNPFFYRSRMTVNKAIFQIVIAWLVPFCYATMPLLGKIVGLNGYRFSVLCNLDVKGVNTTNYVYGYVISSYLIVGVSVTITCYIVIFIVAYSKNVAGYFQDRNLKKSIRTTSLIVGTNMLCWVPLLVLAIAESSAPTQNQFLPPSTQRMVYLLTFLNTAVNPILYTLTNGILRRKLKLFIHRYLSCNRPTKDLPRRRHRVGSTPVQSPRSSLPIKKSASASTCLSAQQVSITKITIPES

>Placozoa_TriAdh113_evg14837 type=protein_ aalen=341_51%_complete-utrpoor_ clen=1989_ strand=| offs=658-1683_ evgclass=main_okay_match_evg111746_pct_99_100_._ 6579

MANGSHFSYSAQPLLHADFKFLIPILTTVCIASMFFNSFNLMVISRKLSLRNIQNSFLTCLNITDLCTGSLAIPCLIVALWYQNGDAICQVQGFIASFCNGMALLLSTTIALDRCYAVVFPYHYLANSSIRRYAIIITSCCISSIIAALLPLLPISLRGFGKYQLKNFCWLSLTASSENYILLVIAAVLTVIIVSTVISCYSIIFYVAYSKSRSRIAHCSSIKKSIKTTCLILGTNILCWLPVTLSGTMSVISFLAHNKAIKINQYVEMIALVLLYCNSVVNPLIYAFTNEILRKEYKKLAAKWSRKIISFCSHQTLVTTFIEKENTESNDCHIKVNSLPS

>Placozoa_TriAdh114_evg1609078 type=protein_ aalen=326_70%_complete_ clen=1397_ strand=-_ offs=1079-99_ evgclass=maina2_okay_match_evg1708920_pct_99_100_._ 6619

MYLNATLSTIKSDQKLLFSPEIISLLFLSTIAIVGNLFNLIVIYKSPLLRSLDNILLIALLITGFSTGLILIPLILLVKLTGNHVLCQIQGFAITLLNCCSLTLTVALSVDRCHAVINPFSYLRQSNKAKYTIIILSIILFSSMISIFPILGLERFGLGKYILGSICWCSLVIDQTNVIILTFLVSAMIIHILIIIFCYIILFAIAYQKGHQDLSGSFGIKTPLRTVILIVGMNMICWIPMIVVLLHGIFQYNHGSTNATLPYWRIAANISFFLLDAYIAVNPIIYLTTNSILRKKWIYLMNLLLCNGRLASSRSLQLPQRRSYKV

>Placozoa_TriAdh115_evg1609534 type=protein_ aalen=340_41%_complete-utrbad_ clen=2485_ strand=-_ offs=2060-1038_ evgclass=maina2_okay_match_evg1751786_pct_99_100_._ 6621

MTIVTEIYSTLHWEMNTTSATLYGYNQSTVPLNNTQLTFVTDLIIRISISLGGFISVVAVFGNLFNIFFIYRYPNLHNMNNLIIANLSFTDFFNGLIVIPMIVVFGLTGRPVYSLCQFQGFILTWLYTASLTTSAAVSADRLYAIMYPFRYYANITVKKLCLTMICIWFVPTIIALIPLLALENYGLGKYRYMAFCAVSFIFYSENYIICWILISFIIITIIIILCSYLAIFTIAYRKTVRDLSHSGNLKKSIRTTSLIVGTNLLCWLPYLFLNFAQYIDRVNNTPDINSSSRSGPTSVVAIFLLFSNAAINPIIYMVTNSYLRKQIWYLFCKTRVHPIP

>Placozoa_TriAdh116_evg1737835 type=protein_ aalen=359_51%_complete-utrpoor_ clen=2092_ strand=| offs=719-1798_ evgclass=main_okay_match_evg14841_pct_99_88_._ 6670

MYIYTRVFSICTDKELLGMDNISDLNNSYRPPLHMNFKFLIPILMIISVVAIFSNFYNLIIIYRKPGLRDFRKSFLTYLNITDLCTGIVTIPCLIIALWYQDGDAVCQVQGFFASFCTGMATFLSCAIALDRCYAVTYPYQYLASVDSRRNNIAIITACIISIIFALLPSLPKPLTGLGKYRIQNLCWISLTVSNENYITITMAAVNTIVIIVVFISSYSITFYTAYIKSRSNFASYGSIKKSIRTTSLIVGTNGICWLPVMISGTISAVHFLLYNKIYKISIYVQVIALTLVYLNAATNPLIYAGTNEILRKEYKKLILKLHSNIMSVCHNQAVISFLPNRETSRVITINIKENSLPM

>Placozoa_TriAdh117_evg1797191 type=protein_ aalen=328_48%_complete-utrpoor_ clen=2052_ strand=-_ offs=1610-624_ evgclass=maina2_okay_match_evg3190_pct_98_100_._ 6717

MATITTQNSTWTSNASKEPGFITWERLSPNDPRFILAVTSCSFICFIAIFGNLRNLIMIYNCRRLHDMNNLCICNLAVVDFLTGLIVIPLGISTLAAETLGYLLCQFQGFFTLLLYLSSLAATSIISVDRCSAVVNPFGYKSHITVSKCAFIAIFTWFIPSIISILPLLGFERIGLGKFYVLLICGISFDHPIENFIVSWILTIFLVVNIVTICMSYLVIFTVAYRKTVQDLTRNSKLSRSIRTTGLIVGTNFLFWLPYIVLNVQDFVYNTSVPPVPTTLNYFTTIACLMALSSPAINPIIYAATSKHLRKRFTRVGIPSQSTYASRE

>Placozoa_TriAdh118_evg3203 type=protein_ aalen=343_81%_complete_ clen=1271_ strand=| offs=59-1090_ evgclass=main_okay_match_evg1730358_pct_99_100_._ 6766

MENTTSFNTTYSAREDGHYLYMQLFTLILITIISVGANFFNLVIIYNSPALWSINNCLLTALTITDFLTGLIVMPCAISALLQGSDAILCKIQGFLVTLFNGASLVMATAVSVDRCSAVYNPYNYLSDLRASRYAISIAAVWLVVLPLAIIPLLPFEKYGLGRYTYQNVCWISLSVDSSNYAAIGIFAASITCAILIIVCCYTIIFYIAYRKSNTPLVSFNGIQKSIRTTFLIVGTSMICWLPIATMTVYGLAYYVKYNRKFIIAHKLQIVILLLSYCNVAINPIIYTMTNGIIRRRILSSVYLFCQLICIFPRFQRGLLNYSFLLNKSQTTSRVTPMEITTS

>Placozoa_TriAdh119_evg3211 type=protein_ aalen=350_43%_complete-utrpoor_ clen=2421_ strand=| offs=414-1466_ evgclass=main_okay_match_evg1609203_pct_99_100_._ 6768

MNLLFLVFMAIITTRVSLFVADTAENQTATATPFIFSDPSFTTNFIIVILIIILSVFGNITILYIFYHSPLLHDVNSLLAANLALINLLISLFILPISLILISQGNINNSLCRVQEFIFILLYAASLAFTASIAVDRCIAIINPIRYVSAVTWRKGFITSLLIWFLPILLATLPELNLIKNGMEILPCRTIRGLYFFNTDKNALIYGLIISFCIVYCLIIVLCYFIVFIIANRKRTALQNFSNGNSLTKSTRAAALLATSTLLCWTPITVALTISYVHTLYRTDTVYKISPYVLRTFTMLAIAVAAVHPIISVCTNHIIRAKFRQIFSRHHCKTCCKRHRTVIPLVLINA

>Placozoa_TriAdh120_evg3218 type=protein_ aalen=333_76%_complete_ clen=1309_ strand=| offs=254-1255_ evgclass=main_okay_match_evg22615_pct_99_100_._ 6769

MAGQPQFLIDNLSVLSKEIVCESTISPTDYDPSQVKITIIATVCIVISLISIPCNLFNIIIIYVSPILHNMNNILLCNLLFIDLMNGLIVLPLITVVSLNRNHDYNLCQLQGIILHLNYSASLTASLAITVDRCFAVVNPYRYKSYINKKRCFLIAICTWTVPTILAFLPVLGLHKYGLGKYFAPIFCGTSFDQFDNNHVLQAILNIFIVVITISIITCYTVIFIIAYEKSAQDILGSSNIKKSIRTTSLIVGTNLLCWLPFMIFNLLEYINRNHEISNKPTPFNITSLLLMLSNAAVNPIIYASTNKFIRKRYHQFFSSKNVFNIHDQNFQS

>Placozoa_TriAdh121_evg3219 type=protein_ aalen=329_79%_complete_ clen=1246_ strand=| offs=226-1215_ evgclass=main_okay_match_evg1776384_pct_99_100_._ 6770

MANSLTTTQISPPLTQFNTSVLNLATNQSISSPNIQFIYNPAKAVIYAVVLSIFSMISIPLNLFNLITLYRNPILHNMNNLLIGNLALVDFLNGLIVLPLIILTSLIGYPGHFICQLQGFFVLMLYLASLSSTSAISVDRFYAVIKPFRYSSIITPKKCIIITLYIWLIPLIFGITPTLGLQKYGLGRYFLLYFCAISFEKFQENYLIHALLMFYIIIIMFVIMLSYGIMFYVAYRKSVQDLTANSTLRRSIRTTVLLVGTNIICWLPWLAYSVDSYQTRNIFNLDKPTYLNIVSLLLVFSNTAMNPVIYSLTNSQLRAKLRKILPSFN

>Placozoa_TriAdh122_evg3453 type=protein_ aalen=326_74%_complete_ clen=1319_ strand=-_ offs=1043-63_ evgclass=main_okay_match_evg1725703_pct_99_100_._ 6776

MAVLTEPNSTLSISNSNGFNFLVWERLSYTDPRYIITVTCLAFISIIAVYGNLRNLTLIHQCRRLHDVNVLCIANLAFIDFLTGIIAIPLAITVLTLNPLEYPICQFQGCVTLLLYMTSLVATSLISVDRCNAVVNPFGYDSQITASKCVFIVILTWLIPFIIAILPLLGLERYGLGKIYLELLCGISFDYLHENFIISWVMTAFVIINILVIFFSYLIIFTVAYRKTVQDLTRNSKLSRSIRTTSLIVGTNFIFWLPYIIFLGQKFSYNISIPPLPKFSDYFTVIAIIMVMSNPAINPIIYATTNAHLRKHFIITIPVPQATDQN

>Placozoa_TriAdh123_evg368847 type=protein_ aalen=336_88%_complete_ clen=1145_ strand=-_ offs=1022-12_ evgclass=main_okay_match_evg1061730_pct_100_41_._ 6783

MDWKRQFESSHQFNSTYSVNASSSTHILSNLRIAIICVCSIISVFSIMGNLLSLMAISRSKYWHTPNGLLLTNLAIVDFLLGLICLPLVIAVAYAGQNVPTVLCQIQAFSLGVLNCTSINTAVIISIDRYVAVAQPYYHAASVGRSKVIKAIVTAWSLSAIMSSIPIIAQGKYGLGIYDNRNQICWFFMRDYAKNHIINSIFITAMLLAIAIVLFCYAIMFGIAYQKSTATPLYHFGSSYRNIKKTIFTTTLIVGTKIICWLPLSVVSVLDACTTVPLTSLVGIMVYFFAFANSALNPMIYIATSSTLRLKVLQLIHRNRRVNNLVTVSSTFTRNT

>Placozoa_TriAdh124_evg712 type=protein_ aalen=369_42%_complete-utrpoor_ clen=2594_ strand=| offs=390-1499_ evgclass=main_okay_match_evg1722407_pct_99_100_._ 6855

MSKFEGYNLTNHFTTTSSSLAANSSILPFTWSLHYNIPILTLFFSIVMIFAVTGNLFSLITIYRSVSLHTINNMFIFNLAVVDLLSSLLVIPLGLAVVSSQSHFSNTLCQIQGFLLMFLNTTSLMTTTLISIDRVIAVTKPYYYAAIAKFHAMIALIAGIWILSILFATLPLLHLQHYGFGYYTLVYICWINIHDYDANFVIAIAAICLILFSMMAIIISYTIIFCIACNKVANQLRPNGYGDVMKSVRTTAIIVGTNLLFWLPFVTACIIEIKDGHYHRRDPDFDETLFKVAYLAPFGSAASNPIIYITTNGVLRLKFQKLFQFKTSLQPLRKRTAISPSRRNNSTAACIVEQINSLNDSYRPDDENQ

>Placozoa_TriAdh126_evg728 type=protein_ aalen=337_70%_complete_ clen=1442_ strand=| offs=350-1363_ evgclass=main_okay_match_evg1420869_pct_99_100_._ 6858

MINLTNSSFPYMPILSYKDIFVVTILGIIASVASFGNLFNLYLIYHSRSSQLVGTLLLANLGLADLFSGLVVMPLAIYSFIVQDTISPFLCQLWAVLASLSIGVSFYTTAAISIDRCIAVASPFYYSQSVNLRVLKVQIVILWLITSVIAIMPMLGLKAYGIGHYSYITGSQQCWVDFTDRDHNYYIGLVLLSILSCTIVIIVLCYSIIFIIAWRKGLQNLSGHGTIKKSIRTTGLIVGTSLMCWIPLITISYVEFIDFMLLISDDIVLAAYLLTFCSSAINPVVYSLTNSELKRRVRLTLRRTNVIAFSQSKRKGKVNPIGIDSVIGNLHSSVIVG

>Placozoa_TriAdh127_evg811142 type=protein_ aalen=352_51%_complete-utrpoor_ clen=2076_ strand=-_ offs=1669-611_ evgclass=noclass_okay_ 6869

MVTINQTNSTIANPGINQILSISFISIVILIGSIGNSINIYFIYKIPSLHSYNNMLIAHLAVADFLQATLTLPMAIANSLLRHAIPPAMCQAFAFILNFLLAIALAATAAISIDRCLAVIYPYQYQAKMKTNYIIMIVIFIWILAAVLAGTPLLGLQRYGLGEYTFIADGLQCWFDFQYRQRSSIIFIILYIYLLLTILTTLVSYLIIFCIAYNKGIADISAVGYASLRRSIRTTALIVGSNFACFIPTLISVSISYFSQRELNPGLTVAAYLLSFCNSAINPLIYAFTNGILRQKIKQHCCCSKKRYCLTKFGLYRTSKITPSNTAAGQPAAKSDSEPSYWQTVINDYESS

>Placozoa_TriAdh128_evg87029 type=protein_ aalen=353_81%_complete_ clen=1305_ strand=| offs=220-1281_ evgclass=main_okay_match_evg892328_pct_99_100_._ 6879

MSRMDPINVLVNVSNHSQNLASNISQNQSYANAPVNFMFMIPALSIISVIAIIANIFCLTTIYHTPSLHDISNVMIANLAIIDLLTGMTVIPTIITVLLLGNETPSWLCKLQGFLTVAMYGASLFAIVGISIDRFRAVTRPYHYSSIITKTIIVKLLMTIPIPIIVASLPLFNLVNHGFGSYIKEFVCWSPACRHNQPAYISSAATAVFIVICIGIVLFCYITIFIIAYHKNKKNIHLKRNINIKSIRTTALIVGTNLICFLPQLSAITSALINCNNNSPEFFQAVFYTITLSNSALNPIVYISTNSILRRKLVQLMLVKALSKKDPYSQRDHLPQSRSLELSNNTSILTTRF

>Placozoa_TriAdh129_evg93544 type=protein_ aalen=344_83%_complete_ clen=1244_ strand=-_ offs=1212-178_ evgclass=main_okay_match_evg595_pct_100_90_._ 6893

MMESILSVDSSSILSVLGSNATKGTYGMQMGLSNSIYLNIVYAAIVVLSTLGNVTYLSVIYRWPELRSTNNIFLANLASANLLMALVVIPFHIIVNVMQYSMTTLVLYQILNSFLLFIFGISLIMMMAVNIDRCFAIASPYNYTGMVTRKKLVNLLILTWCSALIIAVVPLIFFLIMLDRIPQQRSHSTEPIESETEKIELFHTTYRLYLMILIALLFLTICTVLICYSIIFIIALIKGKYNTKVKAFPKHAINIRKSIAITLFITIAHLVCLIPLAMVLLVHCEYKSNVLFRVSINQVNLIHLICLITTALNPLAYIITNRSLRKKLKYLHQYYCNASDAILA

>Placozoa_TriAdh130_evg97324 type=protein_ aalen=328_50%_complete-utrpoor_ clen=1972_ strand=| offs=622-1608_ evgclass=main_okay_match_evg1733964_pct_98_100_._ 6901

MDLTRPPKIYNSSNHLDDYNHKITIQISYLFLITIMAVLGNLFNLFHIFRSSTLRDMKNTMLTGLAFADLCTGSIVPPCMILTLIYQLEDTTCQVQGFVVTYFHGVSLVMSAAISIDRCSAISDPYTYLAHLQVWRYSIIVTSIWIIPLPFAITPLLPFQSLGFGSYELTSLCWLRLSTNKYNYVALAALAAGICSVVTIIICCYVIIFCIAYRKKNSHIDSYGSIRKSIRTAFLIVGSNVFCWLPLGVTWSVSVIKYFVFQGEIRINSQLESAVLLLPYSNVAINPLIYAGTNEILKDRYAKFGVRIYNYVSDRLPLPASNAINDAA

>Placozoa_TriAdh131_evg101767 type=protein_ aalen=411_55%_complete-utrpoor_ clen=2245_ strand=-_ offs=1464-229_ evgclass=main_okay_match_evg1365777_pct_99_100_._ 6443

MEDNFSLVYNYGNLTDDPLSNTQIIICSIILTLATLLSLIGNSLVITVIIRNRPLHTNAGMLMANLAVADFLVGLVIMFPTLIAFICQKDIYPRIACVAQGSAITVLVGASTVTILAITADRFISIFLSLRYNSIVTTAVIIGTIAFSWLFATVASTIPLLGLQKYGLGRYEFLHGMSYCWLDPLQVQQNNVIISIIGFYVVLEIFIITTCYLVIFIYARQSVRKVIHLSVGDLPKGRPKITKTTKTVMIVVGVFMFCWMPSLVNKFLIIFKVKTLSLSVRLIFTWLTFFNSAANPIIYSIFNRNFITGVRNLFKKSMSERSIIVSMSERRSRSFRRRSLSITKFRNSLLPMHRSNTVQDVRFPSCNQSLQVPKIVIAKSKSTEALFKRRRDKIVIHATLNEVPIEIEDVA

>Placozoa_TriAdh132_evg15166 type=protein_ aalen=365_48%_complete-utrpoor_ clen=2262_ strand=-_ offs=2001-904_ evgclass=maina2_okay_match_evg15165_pct_100_99_._ 6599

MNNLSRPTMDINSSYDKISHDYSFNNSDDTDLAILLPILVTFIAIIMTTAMAGNLYIFYAIYRCRSLHKTSSFLIANLAVADILTAIIVLPPMAYSAIQGKMLFNDTFCQISAYINYIIFGASLNTSITITMDRFVAVLWPFRYERITTPSRTAFMISTSWLFPSIVAMLPLLSLQRYGLGRYRYTNSTYYCGIDLHDYQDNYLFNVISAASILNSVAVVAIAYLFIFRVAYKKWVKSQQKSIDIKGNKEASLLRTVRTTALIVGTNILCWLPAVIQVLIYTINPPNQLANKVDVLGIIFVWLPYTNCAVNPIIYISSNSRLRNLVIKQWEYYRRNIMCFKFCHPRNKNVIGPWLTSEPSNSFHL

>Placozoa_TriAdh133_evg1659776 type=protein_ aalen=380_58%_complete-utrpoor_ clen=1962_ strand=| offs=567-1709_ evgclass=maina2_okay_match_evg19761_pct_99_100_._ 6643

MDSINSTQSSLPYYSSTNYLAIVILILLAIACILGNLLSIIVIYRNKNMHDPSGLLMANLATADLCIGLIFMTTTVVYTWKQSYMFNELSCAMRAYLITVIVTVSQFTAAYIAIDRCIYVCKPLHYPMWITKWKMGSLIIFTWILASLFGSIPLMQLERYGLGRYTYVSYMYSCWLDIKKFQQNGSILLLLFNTFIGLLLIVSICYIIILYKAKKVHTTIQPFRRQFNIRQQLPTRMVKLSWINKPTKTSVILVGVYLICAAPTAIFGITMTLNFEYYNVSDAYYRVLLYWCMYANALANPFIFGFSHRDFRQTYRSLCCKNPMRYIMQFLPVKKQPPSGSLSFTIVRSSDSNHHSISHYPASRSTRVNAAGGKNLFLDF

>Placozoa_TriAdh134_evg1747753 type=protein_ aalen=358_40%_complete-utrpoor_ clen=2656_ strand=-_ offs=1869-793_ evgclass=maina2_okay_match_evg19748_pct_99_100_._ 6682

MNELKVKYTCENYNSTLLINITYEYIPTEEALTLTVVVLIFLLTSLVGNIAAVVVVYRNRYLHDPSGILIGALAVSDIGVGFSSVISTIIYDNPCSRLNHAVCVFLAFMSSSFIGSTYSAIAGITVDRCIAICKPLHYRQYVTNTRIIQILLFNWLCILALAALPVMGLQQYGIGKYEYVPYLTSCWIDITAYSKNYLAIIVFYLPVFSILILVMICYVIILLKANSTFKRMAISGWIYNENKQLLKKSTRTTIIVVGVFFICTTPGLALTFATLVNQGSVVSGKAYKYLLNWGIYVNITVNPLLFGFSHKDFRHGYHEICSHFYSKKSKISDISLPIRPSVNPSVLSGYNLVQNSTF

>Placozoa_TriAdh136_evg19758 type=protein_ aalen=337_61%_complete_ clen=1651_ strand=| offs=408-1421_ evgclass=main_okay_match_evg1758788_pct_99_100_._ 6729

MQSECNEWNQSHCFCNQSETSFHITKIILRCITPPVFILALGGNFLSVLVVYRKRSLHNPSGFLIATLAATEMATGLIVTCYFFAFSALHCGMLENFCIVLAYLSTTFVCFSHITVGLISIDRYIAITMPLHYPMYITNKRLSRILILSITHSCFCASLPLMNLNQIQLGRYQYVEYLSLCWIDITDLSENFIFLVIMYSALLMVICIVVSCYVLIFVKANQIHRKRRNLELTKSSVRHLSKKLTRTILIVITVYLIFIIPSTATILITLSHRRLFLSPTMSKIMTFWGLHIKIIINPLIFGFSHRDFRQSYNQLFYYLVKRRRTRIFDQVLRPSIQ

>Placozoa_TriAdh138_evg2834 type=protein_ aalen=323_40%_complete-utrbad_ clen=2394_ strand=| offs=1362-2333_ evgclass=maina2_okay_match_evg101819_pct_99_100_._ 6758

MDRLAESKGLFLSNNITAVHSNYATTTLFTVMTAIIFGSTVFGNIMSLFVIYKTSSLWDNSGILLGNMAVADLIVGLICMPTAFISMCIQRSYLAPILCVITGYINTVVIGTSLQTLAFICVDRCISVLKPLHYHMYVTNNVLRLMLVISWVFPSVVSIIPLLGLQQYGLGKYCFVNYLSTCWLDVIQVQQNRVFLLMIYSLVVIILTCIVICCVLILLVIHKLRNTVADICTSSIHARAVSKSTVTLMLIVGVFILCLMPAVVVTFITCFTNVQFHVTIYQMAICLTYLNSAINPILYGFSSQDFRNSYYTLWKRHFRLYRF

>Placozoa_TriAdh139_evg374227 type=protein_ aalen=362_55%_complete-utrpoor_ clen=1971_ strand=-_ offs=1570-482_ evgclass=main_okay_match_evg1123651_pct_100_98_._ 6790

MSIDDTYLREATWSNIVITVIFSFLFLLIVIGNVLALFVIYRRKSLQDPAGVALANLAVGDLLTSIPLLGSTIIFWIDDAHINPLACKILAFMSFYPFTIILWTTAWITFDRCFAVYHPFKYHRIITVKTVVSLLAIAWILLLVWCMIPLLSLGHGLGNYNYVNYTHSCWVDVGNYRQNGVFLLITYSAVLVLVIIVVLCYSILLIKARLVLRRITTIPTFHYPASSSLPAPSHKSVKTTLIVVGASLMCILPAVIIIYITIMSHRLISEPLIYNLFCFWTFYVDAALNPFIFGFSHKTFRLGYRKLYRHFMSKFITPHHRIYPAIPSAPASVYNSIQSKMNDIASNGYDCSTIFNSNLNNE

>Placozoa_TriAdh140_evg370902 type=protein_ aalen=374_83%_complete_ clen=1341_ strand=-_ offs=1304-180_ evgclass=main_okay_match_evg1315926_pct_99_100_._ 6784

MKGWNWQNTPNGSVDNSTTQHNYSPIYTFQAIALSVAVIAGFPANIITLIACCYRNRLNVVSNIFIANLCIYELCMILLDFPLSLVNAVNGRQIVTSSLCKFQCIIGTMCSAGIILSLVCVTIDRYFTIVKPMKYGMIMTIRKAVTMIICVWVYVFLLIPAYYIPEVQPTCLYYYEMSSCFTNWAQSPIVGYFILATCYGTALIAMIYTYWRIFVIARHHNKVGVEQLKELTKNGYTNRPSITQMRRKGYKTIKVVIVILITYICCWTPITILAATNIQKFSDMPPWGLAIANCLVVINPTINPIVYGIMSQEYRHSYQRIYRIIRNSQTIRRSLIIPSQASFATNQKDGNGNLSLSHDVITTNHESREQETVT

>Placozoa_TriAdh141_evg1041245 type=protein_ aalen=449_37%_complete-utrpoor_ clen=3638_ strand=-_ offs=2666-1317_ evgclass=maina2_okay_match_evg374057_pct_99_100_._ 6459

MQQQIYLRSPNINISDPIPTGGEYGINQSSPYWILQTSSYIGIKLTIHVLLLMIILIGNITAIAVVKSTQLLHRPTGWFILNLAITNFLSGLFLSPIIIITTATASRWSLGDGMCQWVGFVNSTLFTESIITIAMMSVDSYLAIVKPFRYHKWITHNRTLAMLIYSWLQSIVVSILPLIGWSRYTYHPSEAMCFVDIFVGTTFTFFLFGISFICPVIIIVILYILVLRVAVKKVGDMPLMTINQPPPEEKGRRFSLMFRQRQQIRRSKSKLKAVANIFIILGPFLACWLPYIIINIYRLVSNDKKFSPIALSIISTMTILNSAINPILYPRYSKNYRIGLGRLISLYCHPKCLPCYVREYALERSLSFVSEYHHRRLSMRQFDEQNLLKNVDNVRAAAIRHIAVNPLLGRSGSIHPVADDDPTLSTTCSSNTVNGMSKSPTTASVKSQS

>Placozoa_TriAdh142_evg1094259 type=protein_ aalen=343_48%_complete-utrpoor_ clen=2124_ strand=-_ offs=1936-905_ evgclass=maina2_okay_match_evg58816_pct_99_100_._ 6481

MKIESIGHNPNITIYNGSNHIPDENYEEITPLGITSLAIQISIVVLSVICNSLVIATIVGYRRLHNNTNYLVVNQCVVDFLFATLIIAVYPIQALLGYWPFIKEWCAIHYCLSIGLVMVSILNLCAISFDRYIAICHPYHHPRLMTKKSLIIILLFVWIQPLLISLLPIMLWRTDHDLPFGICSYDINRSLIQERVYLASFLMINYVIPAFIMLLLYSLIFRTACRQVHQINKQNMISSSSLSDSIRTHKTSDKDIKLVLLFLAIVGTLYICWTPLYTLMMISYFDVFYQPHIGTFVAIMLAFSNCAINPIIYSCSNAPIRKGLFKLVGIKVNIRSERYVTTV

>Placozoa_TriAdh143_evg1115093 type=protein_ aalen=362_59%_complete-utrpoor_ clen=1836_ strand=-_ offs=1142-54_ evgclass=maina2_okay_match_evg766275_pct_99_100_._ 6489

MDVNSTTDYNISTPVAVIFIIYMSIATVIGVIGNIMALMVIYHQSSLHNTAGILMANLAVADLLITIFNMPATMVSLISSRWPLGSINCQFSGFMMTFATGVSLFTLAVISIDRCKSIIKPLRYQQHYPFRQYIALISLLWLFPCLMATMPLLGLQSISLGKYQYLDSKAYCWIAVDQHRPNLGYVSLVMLSMTAALAIFTTCYFMIYRSAKKSYALIASLTQLCEINRQKFKSNLKTASTLSVIVGVFMFCWSPLVLIDIVSIFAQEPFYNVVSIAALRLALSNSAYNPIIYGIFNQSFRKGFKNIMIHCCLLMKRKHSNEVKQFYAEARGADIKSEIRRKTTIQYSIHIDNNICNNLSVK

>Placozoa_TriAdh144_evg111895 type=protein_ aalen=372_21%_complete-utrbad_ clen=5257_ strand=-_ offs=4525-3407_ evgclass=maina2_okay_match_evg590195_pct_99_100_._ 6491

MDNLSLVVSNDSVNSTSILPTTPGNVKIYTIYEQLILLLYFFTVMVVSVFGNLFVVIVIIKHHQLRTITNFFIINVAAADILVGALVMPFTIYAVITSVHRLTDSRWCIVEWFLSAFLPSASSLSLCCVSLDRFFAITNPYFYRQIVTKQRAVIGEILSWTIAAICASTQFTWTTNPILFCDKLAYGTGLLHDKIYMTVALCIVCFIPILLFIYIYGTIFSIAAAHAKKIRRDSRASSISSRKSMKKSRCNSEQRESKAAKATAIIVGAFVLCWLPFTIVMLVNRWLGRTPLVINAFTAMVVMGNSAINPILYVVFKRDIKCVVKKICCCKRVQQYTRQNNSFSTTHIKLGPNNSSLVETNSNAAETESYRF

>Placozoa_TriAdh145_evg1182385 type=protein_ aalen=348_50%_complete-utrpoor_ clen=2085_ strand=| offs=410-1456_ evgclass=maina2_okay_match_evg485056_pct_99_100_._ 6501

MDNTTSPLSSNLTVTSDVYILRYVTLPIIMFFSIFGNLLVCMVTYSKLQNVTNIMIFNLALADLIVGIVIIPAVLSSTIIGEWIGDATSCKVIGILNASILVVSVWMLVVISIIRYVLVAFPHRSHTLLTKRKAYAVVVLLWIWSFVSAATAELWSPYYYHPQIFDCWFDYSIPFKHQFLMILINILLPLSIMIFAYIKIYLILRKQYRSFQNHIPRTRSMVIRVAMKVGSSVVVPMLNKKSQRSPSQDALGNRLPLARLKDDAKVTRMMFIVLLAFLLAWSPGAFILQFFFFIQPVSPFILNLVFIVGYSNSAINCFIYGFLNKTFRHSFERILTCHKCRKSTNSSF

>Placozoa_TriAdh147_evg12615 type=protein_ aalen=389_46%_complete-utrpoor_ clen=2496_ strand=-_ offs=2149-980_ evgclass=maina2_okay_match_evg370408_pct_99_100_._ 6527

MSWNYRSSNSTIVKEGPQSRSITPQALTFVCITLSIFVVSFAANLIVAGTILCRKHLRNITHLFVFNQCIADLIRVSLQIPPLVVNMLLGYWPFGPVYCNLSYPITIFLMSVSILNVCSISIDRYFSITSPLLYQARQIMSKKIVLVLLLFVWLEPLVIALLPMFVWRDIGYDPRPGSCFVQWTAAPSFERPYLVCLMVVNFFVPCLAMSMIYARIFVIASKHAKRVKYESSAYSIEASTRARIDAKDIRIIAVLATIVGFFLICWLPFFINGLVLLFSPRLMSTVLYTASVYLSFANAAINPLIYAIFTRDIRHQICFLFCRPLASRFHLIYDNYGRNPTRSGLTSQTYSSTPHCRSMVTSDFLIQGSSSLNNARVIMAKGMQEDSTI

>Placozoa_TriAdh148_evg1429139 type=protein_ aalen=426_56%_complete-utrpoor_ clen=2271_ strand=-_ offs=1355-75_ evgclass=maina2_okay_match_evg1738611_pct_99_100_._ 6564

MRAKESSRFLLIIYIHILFFSIKSMAIEFGDIRVEAKGFPPLSGNHVAGNQVSNINSRYGQEIDFKTVYIFTVFITGIICNAIVIATIAYNRQLHSVTNYFIVSQCCADFLFAALVLPAYCLQGLLGYWPFMPEWCAIHACLSISLVMTSLLNLFAISLDRYLAICRPYQYHTWMTTRFVIGMLLYIWFQPLLLALIPVFGWRGVQSTIGSHCVVDLFLYIKQERIYLLIITLINFYIPATIMFTLYILIFKVAIRHIHNIEQQRNVQRKQHYFGSIQDQLNHRYLRLAKNDFRLILVFVVVNGAFYLCWTPFMTFGLISLFHLGPVTDNLGIACSLMAFTNCTLNPIIHNFFNREIRSGFFRMVCFHTRRKPINRSNTHMSGTQCSSTHGSTFVIRHDLISAKDKAIIEQLVNVSRMKAKKVTYI

>Placozoa_TriAdh149_evg1609341 type=protein_ aalen=429_53%_complete-utrpoor_ clen=2394_ strand=-_ offs=1729-440_ evgclass=maina2_okay_match_evg914988_pct_99_100_._ 6620

MIKMDLWNLTYQIKNVTSSGHDGYTAKPNDTDEALPLNPSEWIVIGFSLAVVIISVVCNVIVIATILCNRFLRTTTNYFMISQSIADLLFALIVILGMVIQALLGYWPFAPFWCAIYNACGIGLVMVSILNFCAISIDRYIAICYPFLQHRLNFTLLFILASAYIWMQPIILSLLPMVIWHDYTQVDINVCGHVITHTREKIYFLVLVIVNIFLPALLLSVLYSIVFKTACGHIQKIKQQNSLHRQLSRAADIKRPITSKDFKLLLIFVFIIGTFYLTWTPFFVVLIINYFDDDPYHPSREGFRFFSSIAILISFTHCAINPIAYNFMNSEMRRGLWNLLGISRRRRRASLRLSLQATYTKTSRASRYSKRSNGGPRTSSNCNGQQLVAKNQDNKLTPKAGLEDRLYPLISPLQSYPPNKEPAMALTTV

>Placozoa_TriAdh150_evg19734 type=protein_ aalen=355_45%_complete-utrpoor_ clen=2323_ strand=| offs=484-1551_ evgclass=main_okay_match_evg900155_pct_99_100_._ 6726

MACTNITAILNRTLTPTGINKLAENANIPQEQDSSLNLGEKFKLVTIILIMAVAVIANGIVITAILLSKKLRSKHKFLLSECCANFLFALLIMPLSPIQGLLGWWPFGHIYCNINFSVSIALVFTSILNILALTIERYISICHPFKRHFILKSSHVVVMLLYVWIQPLVLSLIIFIWRDISYQYRPKQCMPESPQKYQTYEKPYFTFLITTNFFIPVLLIAVMYVCIIRIAWRQNQHIRSQHKQLNHSLNKKMNITIKTTDLRATFVSVIILGIFCVCWMPFFIITLTVAYDIRYNPGQFLFISIILAYVNFAINPIIYTMLTSDIKSAVKSLFPCQESYAFRNRSRSLSKTTNV

>Placozoa_TriAdh152_1606805 854 1606805_m.854 type_complete len_372 | 1606805_598-1713| 6418

MSNHSLSSNQTSGFTPLANAEAQVHFTAYVIVMVIAIPANIVTIFSFIIDRTLCTNFNLFLLNMAFIDLLTSLFRIGFNAFCIATIGNSTWPYGQPLCDFDGFIQCVVYAANIYTLMVIAVCRYIAIVHSKQHLITKKVVCIIIAIIWAFSSLCSVFPIMGWNRFRYQPYEHACVPDWSYEKSFPFYVMATEFVLPMIILLYCYTGIFINLRRNSFRMRTVSELNSGNNEAMKKIAQREGKVSRGMFIIFLAFVICFAPYSICMFLLKPVFNIELNRGLVFFCGFMANVNSLLNPIIYAILYKRFRQAYYQVLCYCYSKLTFSSLVDHAGCNRNHGLRSSEMGVIENNNANHSTSPSMKASNDDGKKYRVF

>Placozoa_TriAdh153_evg1004178 type=protein_ aalen=344_36%_complete-utrbad_ clen=2838_ strand=-_ offs=2259-1225_ evgclass=maina2_okay_match_evg1220421_pct_99_100_._ 6432

MTELLNNGTCPSNFTTMNMDTVSIRLVCLSIIGFLGITGNIMVILVTYTKIGRKSPTDYLIINNAVIDAIQSGIRLPLLLVNVAANCNALGDFLCHFHGFLSGVCFIGSAYSLAALAVFRYFSVCKSPAIVLHTRHAVYAIGIVWMITLVICLLPFLGLSVYEYSLMERSCLPSWGSSVANRINAIIELIVDFIVPLVITLTSYLQVFLTLRRIDHNLVRHVSNGGLQRRNTTSSRKPIHLTIALWLLVLNFLLCYITWSILTFIIIPFSNYQISISDDLYFFSVTMLYVNNAINPAIYLVMLKSFRRRYIILLKKIFCWWNKRSSVVRVKPLKSDNTREENSK

>Placozoa_TriAdh154_evg1096055 type=protein_ aalen=416_53%_complete-utrpoor_ clen=2359_ strand=-_ offs=2266-1016_ evgclass=maina2_okay_match_evg768579_pct_99_100_._ 6482

MSVLSCGFLLNSFKLIFERMSVLVSLRGSSHDHHGNATILNFSPAILLARGLAYIILAIATVISNCIVLYVFHKHRSLLRIPSNLFILNLATIDLLNGLFRDTFNGVSWLNNGWTFGEPLCRFNGFMQTICYVVTIFTLLATGICRYLVIVHSYGKVITRKVVLIIIGCIWIYSILDSLMPIFGWNRYIFQPLEFSCLPDWLSVSGASYVMFALIADTIIPSILIGLCYIVIYIVIARNAKRMANRFHIEHSNNVIPSIGNSVDNSVANTPENDRNRQARARRSNGNTLGRKEVKITRAMFSVYVIFLACYLPYALTIFIIAPAGGNVSHNIIFMVGYLVNVNSAINPFFYGVIYRQFRKAYIELFCCCGLSAVSVKFPTSRSRHRHLLNSTPIIDRKNRVDVLRDGMTDASTSNL

>Placozoa_TriAdh155_evg1224494 type=protein_ aalen=346_66%_complete_ clen=1572_ strand=-_ offs=1458-418_ evgclass=maina2_okay_match_evg369914_pct_99_100_._ 6513

MSAGAIHANHCFNNFTIPLSPSYRIWPSIAIGIIFTIALTGNLAIIFFTISIKKLHTSTNILIANVALNDLFITIIVMPICFTISLTLVWPFSCTFCIIGGFLDMILYNNALCSLAILSINRYYAIVCAMDPNFVAKMSIKRAKIMAVGVWMYSAIWAAPPLFGWNRYVFYFSKLMCNMQQRDPSLYVTVVTLFCTVLPTFTILYCSGRVVITLWKNAKNSSIATSNRQRIERRVTSMIGLILLAYVLCIAPNFVAGSLYAFDSSNSEDLSKYNDFQTIATIVYFMNSAVNPIINGAMNPYLRRAVKQQYHWPRCLAAWRHHSVNPITATKKVRKSFSQYVFNTSE

>Placozoa_TriAdh156_evg12947 type=protein_ aalen=465_65%_complete_ clen=2140_ strand=| offs=321-1718_ evgclass=main_okay_match_evg1688447_pct_99_100_._ 6536

MRNLHGHDDDDSYAEGDYFNWTTNHTLPETYGLQEYLQTISIGILFFMSLIGNSLVLRYIVKKRKLHTATFILIANNCAVDLAITIFIMPMCFMVSILRDWPFNYAACQFFGFIDMTLYTGSLMALSAISVIRYFAIVCIRDKNYHLKTSKKTAITMIFICWFYSLAWGSFPLMGWNHYIFFIEKLMCNMNHRDLNHYTTASTICTTLVPCLVVIYCSFRIILAIRQHSHHNHFHQTQQPLQRSRVDRKVTTTLIIVFIAYCSCTIPQVVVASIYKWDEDDTMNQRISYRNIKIMATILYLSNCTVNPIIYGVCNKNLRSAFHFRGYFDRFFDVDGVSQGPTAMGPTSIQVNVNESPTTRSRKSPISLSKVKCFETNSSKNWQAKTLQNQSGLRPMSAKLVNKTSPMVARARLSDVSLVSIESTQTAIEPISPLKSNPNVRNWKNNTGIAVTHPRINHLKNHDIH

>Placozoa_TriAdh157_evg12949 type=protein_ aalen=340_71%_complete_ clen=1428_ strand=-_ offs=1167-145_ evgclass=main_okay_match_evg99149_pct_99_100_._ 6537

MDLNASTNSTSNDKWPQGTEAIIISVILIVIFTIALIGNFPVVFMTYWEKKLHTSTNILIANLAFCDLLITIFIIPICFIVVQVGMWPFSSITCQILGFLEMVSYTASLSSLAMISINRYYAIVRALDQDFAKKMSVKRAKYMIAIVWIYSLVWASPPLYGWNRYSFVLGKLMCNMNHREGQYPLLATIFCTLLPSTILFYCTIRIIVYMWQNFTNRSINLTRSNRLERRVTIMMGIIAIVYFLCIGPNLLMATVVRFNEYSGSLHLYYRIQAIATILYFSNSAWNSIIYGAMNSYLRRIVQQKLLTNICCRCRSDRHSSTVIPLSVRFRSKRANSHLML

>Placozoa_TriAdh158_evg1356946 type=protein_ aalen=359_58%_complete-utrpoor_ clen=1855_ strand=-_ offs=1741-662_ evgclass=maina2_okay_match_evg1265808_pct_99_100_._ 6549

MVSNNSSLYPEYGPTSLAIIRVAAYSVIGVISILGNLLVLIAFYREKSLRIPTNYLIINLAITDILNGIFKDSFIILGTLTNLHIRYKAFCDFDGFMQILLYLLTIYSLLATAIFRYLVIIHSYGKKITVRVVKITVAIIWTYSLAISLSPVLGWNRYVYQRMESTCLPDWLYPAGRSYVMYAFITDSIIPLSLLGICYLCIYIFIRRNAKRFMVNIKKIQPDSLNRSLLRKSIKKEIEITKKMFFVYLVFVICYFPYTIVVFILAPSGVAVPPVAFFIVGFLVNANSAVNPFVYAVIYKSIRKAYYEIIFRYIPSCCSLPASTLDFNQTKLSTAIDKCSTDIIYDTGDQRENLVLVDI

>Placozoa_TriAdh159_evg1417095 type=protein_ aalen=346_70%_complete_ clen=1474_ strand=| offs=368-1408_ evgclass=maina2_okay_match_evg1417096_pct_99_100_._ 6562

MWQNLSTNLIADYHPLQNVWIANVSLTARDRYIRIGLYSVLSLITILGNMLVFLTFYKHASLRTTSNLFIINLAITDLLTGGIKDTLFIYGLTSYNWPKSAILCSFVGFINCVCYVATVYTLTVIAVFRYLAIVCNLGRKIKRKHSILTIACVWIYSSACCLMPIIGWSRYIYEPTECTCVRSLDKKYYSYTIYILVADFLLPLSVVSFCYGNIFAKMKRNHHHIKRDLSDGQKLDVAEKLSRREEIVARRMFIILAEHVVCFLPYTIIVTMLAANGVDIEPVWYFIVGYLLNLNSALNPVTYIVVNPRLKKGYISVICCKDNFTRVKPTKAGGNQGFLELRNVAK

>Placozoa_TriAdh160_evg1429675 type=protein_ aalen=317_61%_complete_ clen=1562_ strand=-_ offs=1377-424_ evgclass=maina2_okay_match_evg1001127_pct_99_100_._ 6565

MYHGQSNYSDYINGNYSGVPISEVEQTLRAAAYTFTALTGLIGNTIVLIIFYQSPSLRVPVNYHLINIAIVDIITCLSRQIFNIIGVVTNGWPYGKDWCDINGFMQTLCYLLTVMSLLAIAVSRYMVIVHRRSVSHKQVCITIIIIWIYSLVHSLLPIIKWNRYFYHKKEFACVPDYKYELTYPLFCVIADFLIPQIIINYCYLKIYLIARNNSLKVFNKSARLDKIRYNVKLTRNMFYVFLAFLICYIPYSISMLFISPFGIEFPVWYIWFCGWLVNLNSSINPILYYFIFRRFRQLYIAMLCKCNCKIAPEVSLR

>Placozoa_TriAdh163_evg19497 type=protein_ aalen=319_24%_complete-utrbad_ clen=3980_ strand=-_ offs=2811-1852_ evgclass=main_okay_match_evg70947_pct_99_100_._ 6723

MADDHAIVNNSNLPLGNAIIRSSVYCLLFITGFFGNLIFLVIFAKCPYMRIPINYFLVNLAIVDITATIFRELALIIGYFNCQWLYNYHTCNLFGFMRNICCAVTIFTLLAISVCRYLVICQHKGRRMTRHAVLCIIAFVWFYTILVCLCPIFGWNRYIYHQNQYVCSIDWSYQPSYIRYIIIVDVFLPTIITIICYTFIYIVVHRHSRRIQDILRMSGSLSSAQYNRIAREKKVTKIMFAVYINFLICYLPFSIITAILIPSGIKPPAELFFVGTFMIDLNSSINPMLYPFIYRRCKEAYCEVLCCHRSQTSPTSNCY

>Placozoa_TriAdh164_evg19528 type=protein_ aalen=387_61%_complete_ clen=1902_ strand=-_ offs=1450-287_ evgclass=main_okay_match_evg1358279_pct_99_97_._ 6725

MSNHSLSLNQTRGFTPYTLAEAQVHFTAYVIVMVIAIPANIVTIFSFIIDRTLCTNFNLFLLNMAFIDLLTSLFRIGFNAFCIATIRNSTWPYGQPLCDFDGFIQGVVFAANVYTLMVIAVCRYIAIVHSKQHLITKKIVYIIIVIIWVFSSLCSVFPIIGWNRYRYQPYEHACLPDWSYEKSYPIYIMATEFVLPMIILLYCYTGIFINLRRNSFRMRTVSELNSGNNEAMKKIAQREGKVTRGMFIIFLAFVICFAPYSVCMFLLASAFNIPIDKGLMFFCGFMANVNSLLNPIIYAILYKRFRQAYYRVLCCCCSKLTFSSSSDHAGCSSNHGLRSLAMGAIENKNANHSTSPSMKTSSILPFRISPFLVLSSIFLVIFFGKLI

>Placozoa_TriAdh165_evg43354 type=protein_ aalen=309_34%_complete-utrbad_ clen=2698_ strand=-_ offs=969-40_ evgclass=main_okay_match_evg936935_pct_99_98_._ 6795

MDNRTNETHHTPLEVIVIKSTIFIIIAIAAVIGNSLVIIVISQYGRNATQTVTNYLIMNLAILDIINGLFQLPLYLVNIIADRQFYGEIGCAIHVQIGGMGFIGSIHNLMWIAIIRYLVVVKSKKLQSRHAYYCIGIIWATCILVPAVSFLGWGKSVYVDIEKGCVPDWNGNSISGIARGIYDAVIDFAIPLIVITYCYLFIFLFLSRKRRDMICDGFQSPTAMDKANVTSTMFIIFIAFLICIAPWSLLAFIIIPISKVIVPDGVYFTSSALVFCNSAINPVLYAYRSKDFRIGYKQAIHKICRCRYN

>Placozoa_TriAdh166_evg70376 type=protein_ aalen=356_79%_complete_ clen=1343_ strand=-_ offs=1186-116_ evgclass=maina2_okay_match_evg19501_pct_99_100_._ 6852

MIGNHTSHETSNMDSSYISSGQISLSVSIYLILTASTVVGNIVVLYTFYLYPYLRVPSNLFIINLAITDILNAMGRNNFVLIGLIMGPEPYTHLFCNISGFINSLGHVVMTATLAATAICRYLVIVRSYARKITTTVVRRIILTTWCYAILNCSMPWLGWSRYTYHPFEFTCLPDWHDENNFSYIIFVFIIDSVIPCILLIACYWRIYAFISANANRMLIHIRQSSLKARRHLPLRIMIKNEIKITKVLFIICVLFMICILPYSIVIFILIPCNVDVSPYVVFYCGVLINLNGSLNPVLYAAIYPKLRKIYRRLGCYLCHYATKIHQESSRRRIVAYPLNDSNRIKIEETKVTTIE

>Placozoa_TriAdh167_evg70886 type=protein_ aalen=391_33%_complete-utrbad_ clen=3482_ strand=| offs=308-1483_ evgclass=maina2_okay_match_evg19493_pct_99_100_._ 6854

MDNKAKYTYNHNNASFYGSSNQNDHDDINREAFASGIIFWRVFIYCVLIAATVIGNGIVLFIFYKRPSLRIPVNFFIINLAITDILTGLTRQPIIVIDLLSRGESFIHTICGLSGVTLCVCYVVTIFTLLATALCRYLVIVRSYGRKLSTKVVVIIMSFIWVYGTIDCLLPIFGWNRYVYQPEEYTCLPDWQDSKGASYIIYILIIDSLIPWTVTAVCYTGIYLFIKKHVHHMLGHLQANTLTVHQQLHCQALQREVKITKVMFLYFLTFILCYMPYCLTMFILVPNNFHISHHLVFFVGCLINLNSAINPVLYALSSKKFRKAYNEIFGRFSLRLYVTHAMSRRTAIIQPFPNPSGLDKINELQNIEQNIFQAISATRIFPQQTISGTEQ

>Placozoa_TriAdh168_evg74047 type=protein_ aalen=341_48%_complete-utrpoor_ clen=2097_ strand=| offs=236-1261_ evgclass=maina2_okay_match_evg910466_pct_99_100_._ 6860

MTWPIMSNSCYENISDLSPTKIGLRAGYLAIIGFFGIVGNIFLLISLFLDKRIKSAINYLVINHTCLDLLQSGVRLPLYIVNIIMNSNGLGYELCTFHGFISGIYFIGTTYTLTAMAIFRYLVVCQSPAVNLAPRHAIYAIVLVWIIAIIMCLLPFWGLSHYDYNIIERSCLPTWNEGLANQINAVFESLIDFGLPLLTTFICYLKIYTVLRKIKMDLMRYRCESITANTCQYSSDRAKKRFTLTLSILFLNFLLCFTVWSIISFIVYPVTAGTACVPDDLHYFAISLLYLNSALNPLIFFARMRSLRMKFYFILETMTGRKQPETTLATSGTKRNNTIER

>Placozoa_TriAdh169_evg7934 type=protein_ aalen=318_75%_complete_ clen=1274_ strand=| offs=254-1210_ evgclass=main_okay_match_evg49754_pct_99_100_._ 6866

MVNDTTSNITSQSYSTVTVISVLSYVILMIVTVSGNALALRVFHRYVELRTPVNYLLINLAIADILKGGIQDGLVLCGLLTPDWGYNRVICNVSGFMLSAFHVGTIFTLTLTGIFRYLIVVNSMKHWITRKVVVAAILVIWLYACLIAFLPILGWNHYTYDKMQLLCLPNITMEISYPIFAITVDIMVPIFLLGYCYLRVYMVMRQNMRRTTVTLHGKTRSRSYSRLHKREVQMTRMMFSVYLAFVICYIPYVIQAYVLIPLQIDVPNVIVFLTGYILNTNSAINPFLYGFTSEKFRQVYSAYLCCLFSCRKRVMPSR

>Placozoa_TriAdh170_evg803040 type=protein_ aalen=353_77%_complete_ clen=1364_ strand=| offs=284-1345_ evgclass=main_okay_match_evg1753631_pct_100_90_._ 6868

MENEIYDGSNTSNILSESIEFTIVKGAILTTLGVVALLGNLMVVFSILLPKKMRSPTNYLLFNLAVLDIITVTIRLPIHLINIMENRQAIDDPLCHFHAFLTGVCFIGSVYSMVGIAVFRYIVVCRSLSVKLSNRHALYAIGLIWLITVFIALWPFWGWSKFVYDPRERACIPSYSEGISGLINGILEIFLDFGIPLTTIFVCYWKIYRYIRATHQRLASFRESANVKHEYRKRRVTLTMWIVFVAFFILYAPWSIMAFVVFPVTNGQANVPDGVYFMAQALLYSNSAANPIIYGVMMGQFKRQYKKVITCGCLTSNEQMIQSDTDAERVVPYASRAITIDWLKAMYINCPIY

>Tadh171_Supposed_placopsin_test_evg1145940 type=protein; aalen=370,57%,complete-utrpoor; clen=1943; strand=+; offs=654-1766; test_evgclass=maina2,okay,match:test_evg370964,pct:99/100/.; 6413

MRSLITPQNETYNKTVPSIVAEASTMTFFAIAAIAGNILVLLSIKRKKSLQTIRNVFVANLAVADLLFAIIGMSLAAVSSVTVRWIFGYEMCQIQGFANSFFCIVSLLTLSAVSIDRYYVILRPLDYHRKMSPRTVTCMVVYIWLHAFVFAALPFTGWSTYRYYYEESLCTADWGHSLSYTLTTIGAAFFLPLAIMAYCYYHILRVARAQSKIIAAEMASLNLPGTPSEDNNNNSNQEHNQSNGSNSLPVDIRDKSRITNAFRKYRREEKATITLMTVMGTFMICWSPHFFGIICLTFPYCPFPKLFFTATTWLALMNSACNPVIYGFLNRKFRHSFYELLGVNKCCKKQKMIINDDVVMGVTYRQSNGE

>Tadh172_Supposed_placopsin_test_evg81053 type=protein; aalen=384,18%,complete-utrbad; clen=6213; strand=-; offs=1326-172; test_evgclass=main,okay,match:test_evg364583,pct:99/100/.; 6414

MADTYINNFTNKSLELCNGSLVVSDCLWRIVLSCFMTLFMLMSCIGNGAVLLVLRYHHDDIKSASNYFITNLALTDFLLGVLCMPCILISCLNGQWVFGQNLCSLTGFANSFFCINSMITLAAVSVEKYCAIASPLTYHHYMSKSKVTCVISIIWIHSAINASLPFLGWGEYVYLPFETICTVAWWSFPNYVGFIVGINFGLPTVIMSCTYFLILKIARKHSRRIGVSTSTVAISTYLSPTGTYNNLSPVNVTTSDHQETLPLPCDTDLGILTSNSISQRIRLQYVREMLTSRRIHYKTHIKATLMLLIVIGSFIVCWLPHLISMVYLTIYEISPLPCSFHQITTWLAMANSAFNPIIYGAMDTSIRKGLKTLLGSWVKYCKLY

>Tadh173_Supposed_placopsin_test_evg17446 type=protein; aalen=340,22%,complete-utrbad; clen=4518; strand=+; offs=1385-2407; test_evgclass=maina2,okay,match:test_evg372020,pct:99/100/.; 6415

MDNNNSNTTVPKMDLVYLILQATILSIITLFMIVGNFIVIVVVNRSEQLQNATGIFMANLAVTDLTLGVILMPITIASSILGRWIFSDIMCKFCGFLNVMLCSTSALTVMLLSIDRFIAISRPMQYIKIMSKKRALVLSTYMWIHSAIVSTFPLLGWAKYEYVEAEAVCFAIESVSYFNFLCASTVFLAIFLIIITHIYILKVAIKQSQQIVTLTPGFEDVTRETMRQMNRKTAKTVLIIVGVYLACWIPYISLTYVQIYANYVPPPIAITITSGLIFVNSASNPIIYGTFHRRFRNAFIRSFMPILTKCSIVDEYAHNREQASYLHTSNTVRKLNNSNH
