## SupplementaryFile2 for "Functional and phylogenetic analysis of placozoan GPCRs reveal the prebilaterian origin of monoaminergic signalling"

>Tadh111_evg17738 type=CDS; aalen=334,51%,complete-utrpoor; clen=1946; strand=+; offs=292-1296; evgclass=main,okay,match:evg36580,pct:99/100/-sense

ccaccATGAATTTAACAGTCACTGTGCAAACCTTTTCCTTTATTGTATCAATGACTACCATACTGTTGGCTGCAGTGAGTATTGTGGCTAACACCTGCATCTTGATTATTTTTTTTAACGTAAGAAGATTAAGATCAACGACTAATTGTTTAGTCGCCAATCTTGCTGCCTCTGATCTTTGCTTCAGTATTTTTGCAATACCCTGCCAGTTATTCGAAAGATTTCTACCTGAATCTGAAGCAATCTGCTTACTAAATTCGATATCAACTATATTTTTCACTACTTCGTCTATCTTCAGCCTTTGTTTCATTGGTATTATAAAATATATTAATATTACTAAACCTTTGAAAAATAACCTTATCATCACTCGAACTCGGGCATATAAAGCTATCGTATTTATTTGGCTGTTTTCTTCTCTATTAACGTTCGTACCCGTTATTTTTATAACGCATAGAAAATTACTATGTCATGGATTTAAATACCGAACCGGAAAGCTTCTTGATAAATTTTATTTTTTAACCGTGTTTGTTATAACTTATCTTATACCAACTGTAATATTAATTTATGTATACACAGTTATATATAAAATAGCTCGACGCCAATGCTCACGTATACGGCGATCAATTAAATCCATTCATCGTAATACAAATGAAATTAGGTCCTCTTTGCCTTCATCGAAATCTGCAACATTTTTGTTGGCTTTAGTTACCTGTTTTATCGTATGTTATACGCCATTCTTTTGTTCTGAATTAATTCATAGATTTATTGCACCGATATCGAATAATGTATCGCTATATATTACAATTTGTTTCATTACTGCTAATTCAGCGATTAATCCCTTCCTGTACGGAATACTAAAACCCGAATTCAAATATAATATTCAATTTTATTTTCGGTTATCATTATGCAAAGACGACGGCAATATTACTCATTCATCAAGCTATAGCCAAAGATTACCGTTTCAGCGATCACCGAGTCAAGTATCTACAACAAATCCACGATATGTATAA

>Tadh112_evg100509 type=CDS; aalen=368,37%,complete-utrbad; clen=2958; strand=-; offs=2638-1532; evgclass=maina2,okay,match:evg869784,pct:99/100/.

ccaccATGATTGATCTCACTACTACCAATGTCAGTACGGAAGCTACTACATTGGATCAATGGCAACCATCTAGTGCTTTTATATATTACTTTATTCCAGGAGCTGTGATCATTATGATCGTGTCTGGTTTAGGTAACACTTTCTGCTTAACCATCGTCTGTCGGACATCATTAAAACAAGATGTAGATACACTATTTATTGCCAATTTAGCCATAACGGATTTAATTAGTGCGCTGTTAGTCTTGCCAAGCATTATTGTAACCAATATGTGCTACGTATATAATCAATTATCACAAGGACTATGTAATTTTCAAGGCTTTCTCCTGATTTGGATTATGGGTACTTCACTTTTTACCACTGCTCTTATTAGCATTGATCGCTGTATCGCAGTCACTAATCCGTTCTTTTATCGCTCGAGAATGACCGTCAACAAAGCCATCTTTCAAATTGTTATAGCATGGCTTGTTCCTTTTTGCTATGCTACTATGCCATTACTAGGCAAAATTGTGGGTTTAAATGGGTATCGATTTAGTGTCTTGTGTAATTTAGATGTCAAAGGAGTCAACACAACCAATTATGTCTATGGTTATGTTATTAGTAGTTATTTAATCGTTGGTGTTAGCGTAACTATCACTTGTTACATTGTTATTTTTATTGTTGCTTACAGTAAAAATGTAGCTGGTTACTTTCAAGATCGAAATCTGAAAAAATCTATTAGAACAACATCTTTAATTGTTGGCACTAATATGTTGTGTTGGGTACCATTATTGGTACTCGCCATCGCGGAATCGTCTGCTCCAACACAAAACCAATTCTTACCACCTTCTACTCAGAGAATGGTTTATTTACTTACGTTCTTAAATACGGCAGTTAATCCAATATTATATACACTTACTAATGGTATCTTACGACGGAAACTAAAGTTATTTATCCATCGATACTTGTCTTGTAATCGACCTACAAAAGATTTACCACGTCGTCGACATCGTGTGGGTTCTACCCCTGTGCAATCGCCAAGATCAAGTCTTCCTATAAAAAAGAGCGCCAGTGCAAGTACATGTTTATCAGCACAGCAAGTATCCATTACCAAAATAACCATACCGGAATCATGA

>Tadh113_evg14837 type=CDS; aalen=341,51%,complete-utrpoor; clen=1989; strand=+; offs=658-1683; evgclass=main,okay,match:evg111746,pct:99/100/.

ccaccATGGCCAACGGTTCGCATTTTAGTTATAGTGCTCAGCCTTTACTTCATGCGGACTTTAAGTTTCTTATTCCAATTCTAACGACAGTCTGTATTGCGTCAATGTTTTTTAATTCATTTAATTTGATGGTTATTTCTCGCAAGTTAAGCTTACGCAATATCCAAAATAGTTTTCTAACATGCTTAAATATTACCGATTTATGCACTGGTAGTCTAGCTATACCATGTTTAATAGTAGCGTTATGGTATCAGAACGGCGATGCTATATGCCAAGTTCAAGGTTTCATTGCCTCATTTTGCAATGGTATGGCATTACTTTTATCCACGACTATCGCCTTAGATCGATGCTATGCTGTCGTATTTCCTTACCACTACTTAGCAAATTCAAGTATTCGACGATATGCTATCATTATAACTTCTTGCTGTATTTCATCTATCATTGCCGCTTTACTACCTTTATTGCCAATATCTTTGCGCGGCTTCGGTAAATATCAATTAAAAAATTTTTGTTGGTTGTCATTAACGGCATCCAGTGAAAACTATATCCTTCTTGTAATTGCAGCGGTGCTTACAGTTATTATTGTCAGTACAGTCATTAGCTGTTATTCTATCATTTTCTATGTAGCTTATAGTAAGTCAAGATCTCGCATTGCTCATTGTAGTAGCATCAAAAAATCTATTAAAACAACATGTTTAATTTTGGGTACAAATATTCTGTGTTGGCTACCAGTAACTTTGAGTGGAACTATGAGTGTGATATCATTTCTTGCTCATAATAAGGCCATAAAAATTAACCAATACGTAGAAATGATTGCGTTAGTATTACTTTATTGCAATTCCGTTGTAAATCCGCTAATATATGCCTTTACTAATGAAATATTACGAAAAGAATATAAGAAGTTGGCAGCAAAATGGTCTAGAAAAATAATTTCCTTTTGCAGTCATCAAACGCTTGTAACCACATTTATCGAAAAAGAAAACACCGAAAGCAACGACTGTCATATTAAAGTGAACTCTTTACCTTCCTAA

>Tadh114_evg1609078 type=CDS; aalen=326,70%,complete; clen=1397; strand=-; offs=1079-99; evgclass=maina2,okay,match:evg1708920,pct:99/100/.

ccaccATGTATCTTAATGCCACACTTTCTACAATTAAAAGCGATCAGAAATTACTGTTCTCACCTGAAATAATCTCATTATTGTTTCTATCAACGATAGCTATTGTTGGTAACTTATTTAACTTAATAGTAATTTATAAATCGCCATTACTGCGTTCTCTTGATAATATTTTACTTATCGCGTTATTGATTACTGGCTTTAGTACTGGGTTAATTCTTATTCCTTTAATATTATTGGTAAAGTTAACGGGTAATCATGTATTATGCCAAATACAAGGATTTGCAATTACGCTTTTAAATTGTTGCTCGTTAACTTTAACAGTTGCTCTCAGTGTCGATCGTTGTCACGCTGTTATAAACCCTTTTAGCTATTTGCGTCAATCTAATAAGGCCAAATATACAATAATCATTCTTTCTATAATACTGTTTAGCAGTATGATCTCAATATTTCCAATATTAGGCTTAGAAAGATTCGGTTTAGGTAAATATATTCTGGGTTCGATCTGTTGGTGTTCACTTGTTATCGATCAAACTAATGTCATAATTCTAACATTTTTAGTCTCGGCTATGATTATCCATATCCTAATAATTATCTTCTGCTATATTATACTCTTCGCTATTGCATATCAGAAAGGACATCAAGATCTATCAGGGTCTTTTGGTATTAAGACGCCTTTACGAACTGTTATTTTAATTGTAGGCATGAATATGATTTGTTGGATACCCATGATAGTAGTCCTATTACATGGAATATTCCAATATAATCATGGAAGTACGAACGCTACATTACCCTACTGGCGAATTGCAGCTAATATTTCGTTCTTTTTATTGGACGCCTATATTGCAGTTAATCCGATTATTTATCTAACTACTAATTCGATTTTACGTAAAAAATGGATCTATTTAATGAATCTATTATTATGCAATGGCAGGCTGGCTTCCTCTAGAAGTTTGCAATTGCCACAACGTCGAAGCTATAAAGTATAA

>Tadh115_evg1609534 type=CDS; aalen=340,41%,complete-utrbad; clen=2485; strand=-; offs=2060-1038; evgclass=maina2,okay,match:evg1751786,pct:99/100/.

ccaccATGACAATTGTGACAGAAATTTATTCAACCTTACACTGGGAAATGAATACTACATCAGCAACTTTGTATGGCTATAATCAATCGACAGTACCGTTAAATAATACGCAACTAACATTTGTTACTGATCTAATTATAAGAATATCCATTAGTCTAGGCGGATTTATTTCCGTGGTTGCAGTTTTTGGTAATTTGTTTAACATTTTCTTCATTTACCGATATCCAAATTTGCATAATATGAACAATTTAATCATCGCCAACTTATCCTTTACAGATTTTTTTAATGGTTTAATTGTTATACCTATGATTGTTGTGTTTGGATTAACTGGTAGACCGGTATATTCCTTATGTCAATTCCAAGGATTCATTCTGACTTGGCTATATACAGCTTCTCTAACAACATCAGCGGCTGTCAGTGCTGATCGCCTTTACGCTATAATGTATCCTTTCCGATATTATGCCAACATAACAGTGAAGAAACTCTGCCTTACTATGATATGCATTTGGTTCGTACCGACTATAATAGCTCTGATACCTTTGCTTGCCTTAGAAAATTATGGCTTAGGCAAATATAGATATATGGCATTTTGCGCCGTTAGTTTTATCTTCTACAGCGAAAATTACATAATTTGTTGGATATTAATTTCCTTCATTATTATAACTATTATAATAATTCTATGTAGCTACTTAGCCATATTCACTATCGCATATCGAAAAACAGTTCGAGATCTATCCCACAGTGGAAATTTGAAAAAATCTATTCGAACAACTTCGTTAATAGTAGGTACAAATCTACTATGCTGGCTTCCATACTTGTTTCTAAATTTTGCTCAGTACATCGATCGTGTCAATAATACCCCCGATATTAATTCCAGCAGTAGATCAGGTCCCACGTCGGTAGTTGCTATCTTTTTATTATTTAGCAACGCTGCTATTAATCCTATTATTTATATGGTAACGAATAGCTATTTAAGAAAACAAATATGGTATCTGTTTTGTAAAACTCGCGTACATCCTATACCGTAA

>Tadh116_evg1737835 type=CDS; aalen=359,51%,complete-utrpoor; clen=2092; strand=+; offs=719-1798; evgclass=main,okay,match:evg14841,pct:99/88/.

ccaccATGTATATATATACTCGTGTATTTTCAATTTGCACAGACAAAGAATTATTAGGCATGGATAATATATCGGACTTGAATAATAGCTATCGACCTCCACTCCATATGAACTTTAAGTTCCTTATTCCAATTTTAATGATAATCAGTGTCGTGGCGATATTTAGTAATTTCTACAATTTGATTATTATTTATCGAAAGCCAGGATTACGCGATTTCCGAAAAAGCTTTTTAACGTATTTAAATATAACTGATTTATGTACTGGTATTGTTACTATACCATGTTTGATCATAGCATTGTGGTATCAAGATGGCGACGCAGTATGTCAAGTCCAAGGTTTCTTTGCTTCATTCTGCACTGGTATGGCAACATTTTTATCCTGTGCGATTGCTTTGGATCGGTGCTACGCTGTAACTTATCCTTACCAGTATTTAGCGAGTGTAGATAGTCGACGTAATAACATCGCCATCATTACTGCTTGCATTATTAGCATAATTTTTGCTCTACTTCCTTCGTTACCAAAACCTTTAACCGGTTTGGGTAAATATCGAATACAGAATTTATGCTGGATTTCGTTAACGGTATCTAATGAAAATTATATCACTATTACAATGGCAGCTGTTAATACAATTGTTATTATTGTCGTATTTATAAGTAGTTATTCTATCACTTTCTATACAGCTTACATTAAATCAAGATCTAATTTTGCCAGCTACGGTAGTATTAAAAAGTCCATTAGGACTACTAGTCTAATTGTGGGTACAAATGGTATTTGTTGGCTACCTGTAATGATATCTGGAACCATTAGTGCTGTACATTTTCTACTTTATAATAAGATATATAAAATAAGTATTTATGTCCAAGTTATCGCGCTGACATTAGTCTACTTAAATGCAGCTACAAATCCACTTATATACGCTGGCACGAATGAAATATTAAGAAAAGAATATAAGAAATTAATACTAAAGTTACATTCAAATATCATGTCTGTATGCCACAATCAAGCAGTTATTTCCTTTTTACCGAACAGGGAAACATCGAGAGTAATAACAATTAATATTAAAGAGAACTCATTGCCTATGTAA

>Tadh117_evg1797191 type=CDS; aalen=328,48%,complete-utrpoor; clen=2052; strand=-; offs=1610-624; evgclass=maina2,okay,match:evg3190,pct:98/100/.

ccaccATGGCTACTATAACAACACAAAATTCAACATGGACAAGCAACGCTAGCAAAGAACCCGGCTTCATAACTTGGGAAAGGCTAAGTCCAAACGATCCGAGATTTATTTTGGCTGTAACTTCTTGTTCATTTATCTGCTTTATAGCAATATTTGGCAACTTGCGTAACTTAATAATGATATATAACTGCCGAAGATTACACGATATGAACAATTTATGTATCTGTAATTTAGCCGTAGTTGATTTTCTAACTGGCCTTATTGTTATACCTTTGGGAATATCTACGTTAGCAGCAGAGACATTAGGATACTTACTTTGCCAGTTTCAAGGATTTTTTACGTTACTCTTGTACTTATCTTCGTTAGCAGCAACATCAATAATAAGTGTAGATCGATGTAGTGCAGTTGTTAATCCTTTTGGCTACAAAAGTCATATAACTGTATCTAAATGTGCCTTTATTGCTATTTTTACTTGGTTCATACCGAGTATTATTTCTATATTACCACTACTTGGCTTTGAGAGAATTGGCCTTGGTAAATTCTATGTCTTACTCATTTGCGGTATCTCCTTCGATCACCCAATAGAAAATTTTATAGTTTCATGGATACTGACGATATTTTTGGTAGTCAACATAGTAACAATCTGTATGAGCTACTTAGTTATTTTTACTGTAGCATATCGGAAAACTGTGCAAGATTTAACTCGAAACAGCAAATTATCTCGATCGATTCGAACCACTGGTTTAATTGTTGGAACTAACTTCCTTTTTTGGCTGCCATACATTGTTCTTAATGTACAGGATTTCGTCTATAATACCTCGGTCCCTCCAGTACCAACGACATTGAACTATTTTACGACCATTGCATGTCTCATGGCACTGAGTAGTCCTGCTATTAATCCAATAATTTATGCAGCAACGAGTAAACACTTGCGAAAGCGATTTACTAGAGTCGGAATACCCTCTCAATCTACGTATGCAAGTAGAGAATAA

>Tadh118_evg3203 type=CDS; aalen=343,81%,complete; clen=1271; strand=+; offs=59-1090; evgclass=main,okay,match:evg1730358,pct:99/100/.

ccaccATGGAGAATACCACATCATTTAATACTACCTATTCCGCGAGAGAGGATGGACATTACCTTTACATGCAGCTTTTTACTTTAATTTTAATTACTATAATATCTGTTGGAGCTAATTTTTTCAACTTAGTCATTATTTACAACTCTCCAGCCTTATGGTCGATCAACAACTGTTTATTAACAGCACTTACTATCACAGATTTCCTGACGGGTTTAATTGTAATGCCTTGTGCTATTTCGGCATTACTTCAAGGCAGCGATGCTATCCTGTGTAAGATACAAGGTTTTTTAGTAACATTATTCAATGGGGCGTCACTGGTAATGGCAACAGCAGTAAGTGTTGATCGATGTTCTGCTGTTTATAATCCATATAATTATTTATCAGATTTACGAGCATCGCGTTATGCAATATCAATTGCAGCAGTTTGGCTCGTAGTTCTTCCATTAGCCATTATTCCACTACTGCCTTTTGAAAAGTATGGCTTGGGTAGATATACCTACCAGAATGTTTGCTGGATTTCGTTATCTGTTGATTCCTCAAATTACGCTGCTATTGGAATATTTGCAGCTTCCATTACTTGCGCAATTCTTATTATTGTGTGCTGTTACACAATTATATTTTATATTGCATATCGTAAAAGTAATACTCCTTTAGTGAGCTTCAACGGAATTCAAAAATCAATTCGTACTACGTTTTTAATTGTCGGCACTAGTATGATATGTTGGTTGCCAATTGCAACAATGACTGTCTATGGATTGGCATATTACGTTAAGTATAATAGAAAATTTATTATTGCACACAAACTTCAAATTGTTATCCTGTTATTATCTTACTGCAATGTAGCCATAAATCCAATAATTTATACCATGACAAACGGCATCATAAGGAGGCGAATATTATCATCGGTATACTTATTCTGCCAATTAATCTGTATTTTTCCAAGATTTCAACGCGGTCTGTTGAATTATTCATTCCTCTTAAATAAGAGCCAAACAACTTCTCGCGTAACACCGATGGAAATAACTACGTCCTAA

>TriAdh119_evg3211 type=CDS; aalen=350,43%,complete-utrpoor; clen=2421; strand=+; offs=414-1466; evgclass=main,okay,match:evg1609203,pct:99/100/.

ccaccATGAATCTTTTATTTTTAGTATTTATGGCGATCATTACAACCCGAGTATCTCTCTTCGTTGCTGACACGGCGGAGAACCAAACAGCAACAGCTACACCATTTATCTTTAGCGACCCTAGCTTTACTACCAATTTCATCATTGTGATTCTAATTATTATTTTATCAGTTTTCGGTAATATAACTATTTTATATATTTTCTATCATTCGCCTTTACTGCACGATGTTAATAGCTTATTAGCTGCTAATTTAGCCTTAATCAATTTATTGATTAGCTTATTTATTCTACCTATCAGTTTAATTCTGATAAGCCAAGGTAATATTAACAATTCTTTATGTCGAGTCCAAGAATTTATCTTTATTTTATTGTACGCTGCTTCACTGGCTTTTACAGCTAGTATTGCTGTTGATCGTTGTATAGCGATTATTAATCCAATTCGCTATGTATCAGCCGTAACCTGGAGAAAAGGTTTTATTACATCATTATTGATATGGTTCCTTCCTATCCTATTAGCGACGCTGCCTGAATTAAATTTAATAAAAAACGGAATGGAAATCCTCCCATGTAGAACTATACGCGGGCTATATTTTTTTAATACTGATAAGAATGCCCTTATATATGGCTTAATAATTTCATTTTGCATCGTATATTGCCTCATTATCGTTCTATGTTATTTTATTGTTTTTATTATCGCTAACAGGAAGAGAACGGCCTTGCAGAACTTTTCTAATGGTAATAGTCTAACAAAATCAACGCGTGCAGCTGCTTTACTAGCTACTTCAACGCTACTGTGCTGGACACCAATAACTGTAGCCCTTACTATCAGTTACGTTCATACGTTATATCGAACAGATACGGTGTATAAAATTTCACCATATGTACTTCGAACTTTTACGATGTTGGCTATTGCTGTCGCAGCTGTTCATCCTATCATTTCCGTTTGTACAAATCACATAATTCGAGCTAAATTTCGGCAAATATTCAGTAGGCATCACTGTAAAACCTGCTGTAAACGTCATCGAACAGTTATACCACTGGTCCTAATCAATGCTTAA

>Tadh120_evg3218 type=CDS; aalen=333,76%,complete; clen=1309; strand=+; offs=254-1255; evgclass=main,okay,match:evg22615,pct:99/100/.

ccaccATGGCTGGTCAGCCACAATTTTTAATCGATAATTTATCAGTTCTATCGAAAGAAATTGTATGCGAATCCACAATATCTCCAACAGATTATGATCCAAGTCAAGTAAAGATTACCATTATTGCCACCGTGTGTATTGTAATAAGCTTGATTTCTATCCCATGCAATCTATTTAATATCATTATTATATATGTTTCACCGATTCTTCACAACATGAACAACATATTGTTGTGTAACTTATTGTTTATAGATCTTATGAACGGCTTAATCGTTCTTCCGCTGATAACTGTAGTATCGCTGAATAGAAATCATGACTACAACTTATGCCAATTGCAAGGAATCATTCTACATTTAAATTATTCGGCATCTTTAACGGCTTCATTGGCTATAACCGTAGATCGCTGTTTTGCAGTGGTCAATCCATACCGCTATAAATCCTACATTAATAAGAAGCGATGTTTTTTAATAGCCATATGCACCTGGACTGTCCCTACCATTCTTGCATTTCTACCGGTATTGGGTTTACATAAGTACGGTTTGGGAAAATATTTCGCGCCGATTTTCTGTGGAACTTCATTTGACCAATTTGATAATAACCACGTCTTGCAAGCCATATTAAACATCTTTATTGTTGTAATCACAATTAGTATTATAACTTGTTACACCGTTATATTTATTATTGCGTATGAAAAATCGGCACAAGATATTTTGGGAAGCAGCAACATAAAAAAGTCGATTCGAACTACATCTTTAATTGTTGGCACTAACCTTTTATGTTGGTTACCTTTCATGATTTTTAATTTACTTGAATATATCAACCGTAATCATGAAATATCCAACAAGCCAACTCCTTTCAACATTACTTCCTTGCTACTAATGCTTAGCAATGCCGCTGTCAATCCTATCATATATGCTAGTACAAACAAATTCATACGTAAGCGTTATCACCAATTTTTTTCTTCCAAAAATGTGTTTAATATTCATGATCAGAACTTTCAATCTTAA

>Tadh121_evg3219 type=CDS; aalen=329,79%,complete; clen=1246; strand=+; offs=226-1215; evgclass=main,okay,match:evg1776384,pct:99/100/.

ccaccATGGCCAACAGCTTGACCACTACCCAGATAAGCCCACCACTTACACAGTTTAATACAAGTGTCTTAAATTTGGCTACCAACCAATCAATATCTAGCCCAAACATACAATTCATATACAACCCTGCGAAAGCCGTCATATACGCTGTTGTTCTATCTATTTTCAGCATGATATCCATTCCACTTAATCTATTTAATTTAATCACCTTGTATCGTAATCCTATCTTGCACAACATGAACAATTTGCTCATTGGAAACCTAGCTTTAGTCGATTTTCTAAATGGACTGATTGTTTTACCACTTATTATTCTTACATCGCTAATAGGCTATCCAGGCCATTTCATATGCCAATTACAAGGATTTTTCGTACTTATGCTGTATTTGGCTTCGTTATCGTCTACATCAGCAATTAGTGTTGACCGTTTTTATGCAGTTATCAAGCCCTTTCGTTATAGTTCCATCATTACTCCTAAAAAATGTATCATTATAACATTATATATTTGGCTTATACCGTTAATATTTGGGATTACACCTACGCTTGGATTGCAAAAATATGGTTTGGGACGTTATTTTCTGCTATATTTTTGCGCGATCTCTTTTGAAAAATTTCAAGAAAACTACTTGATACATGCCTTACTAATGTTTTATATTATTATTATTATGTTTGTGATCATGCTTAGCTATGGAATTATGTTTTATGTGGCTTACCGAAAATCTGTACAAGATTTAACTGCGAACAGTACGTTAAGACGATCAATTAGAACTACGGTCTTACTGGTAGGAACAAATATTATTTGTTGGCTTCCCTGGCTGGCATATAGTGTTGACAGTTACCAAACACGTAATATATTCAATCTTGATAAACCTACCTATTTAAATATTGTGTCCTTATTACTTGTATTCAGCAATACAGCTATGAATCCTGTTATATATTCCTTAACAAATAGTCAATTACGTGCAAAATTAAGAAAAATCTTACCGAGTTTCAATTGA

>Tadh122_evg3453 type=CDS; aalen=326,74%,complete; clen=1319; strand=-; offs=1043-63; evgclass=main,okay,match:evg1725703,pct:99/100/.

ccaccATGGCTGTGTTAACTGAACCAAATTCTACTTTGTCAATTAGCAATAGCAATGGATTCAACTTTTTAGTTTGGGAGAGATTAAGCTATACCGATCCAAGATATATTATTACGGTAACTTGTTTGGCATTCATCAGTATTATAGCAGTTTATGGTAACTTACGCAATCTAACATTGATCCATCAGTGTCGAAGACTACATGATGTAAACGTTTTATGTATCGCTAATTTAGCCTTTATCGATTTTCTTACTGGGATCATAGCCATACCTCTTGCAATCACTGTATTAACCCTAAATCCATTAGAATATCCCATTTGCCAATTTCAAGGATGCGTTACATTACTCTTATACATGACTTCGTTAGTAGCTACATCCCTAATAAGTGTCGATCGATGTAATGCCGTTGTGAATCCTTTTGGTTACGATAGTCAGATCACAGCAAGTAAATGTGTCTTTATCGTAATTTTGACTTGGCTTATACCCTTCATTATCGCTATATTACCTCTACTAGGTCTCGAGCGATATGGTCTTGGTAAGATTTATCTCGAGCTGCTTTGTGGTATCTCATTTGATTATCTACATGAAAATTTCATAATTTCATGGGTAATGACGGCATTTGTGATTATCAACATATTAGTAATCTTTTTTAGCTACCTAATTATTTTTACTGTGGCATATCGAAAAACAGTACAAGATTTAACCAGAAACAGCAAACTATCAAGATCAATTCGAACCACCAGTTTAATCGTCGGAACAAATTTTATTTTCTGGCTTCCGTACATTATCTTTCTGGGACAAAAATTTAGCTATAATATTTCTATCCCTCCATTGCCCAAATTTTCAGACTATTTTACAGTCATTGCAATCATTATGGTAATGAGCAATCCTGCCATCAATCCAATTATATATGCAACAACCAATGCTCATCTACGAAAGCACTTTATCATAACCATTCCGGTACCACAAGCTACTGATCAAAACTAA

>Tadh123_evg368847 type=CDS; aalen=336,88%,complete; clen=1145; strand=-; offs=1022-12; evgclass=main,okay,match:evg1061730,pct:100/41/.

ccaccATGGATTGGAAAAGGCAATTCGAATCGAGCCACCAATTTAATTCCACCTATTCTGTCAATGCTTCATCTTCCACACATATATTAAGTAATTTACGAATTGCAATTATATGTGTTTGCTCGATCATCTCAGTATTTAGTATAATGGGTAATTTGTTGTCGCTGATGGCTATCTCTAGATCTAAATACTGGCATACACCTAATGGTTTACTCTTAACTAATCTTGCTATAGTTGACTTTTTACTTGGGTTGATATGCTTACCGCTAGTAATTGCAGTAGCTTATGCAGGTCAAAATGTTCCAACAGTTTTATGTCAAATTCAAGCATTTTCATTAGGAGTTCTCAATTGTACTTCTATCAATACTGCTGTAATTATCAGTATCGATCGCTATGTAGCAGTAGCTCAACCTTATTATCATGCTGCATCTGTAGGTAGAAGTAAAGTTATTAAAGCAATCGTCACCGCTTGGTCACTATCGGCAATAATGTCCTCGATACCAATTATCGCTCAAGGAAAGTATGGACTAGGAATCTATGATAATCGAAATCAAATATGTTGGTTTTTTATGAGAGATTATGCTAAAAATCATATCATTAATAGCATATTTATTACAGCAATGTTGTTAGCTATTGCTATCGTTTTATTTTGCTATGCAATTATGTTTGGAATTGCTTATCAGAAAAGTACTGCTACGCCTCTGTATCATTTTGGAAGCAGCTACCGTAATATTAAGAAAACCATTTTTACTACCACCTTAATAGTGGGAACTAAAATAATTTGTTGGCTGCCATTATCAGTTGTCAGTGTTCTTGATGCTTGCACTACAGTACCATTAACTTCTTTAGTAGGAATTATGGTTTATTTCTTTGCTTTTGCCAATTCGGCTTTAAATCCGATGATTTATATAGCTACTAGTAGTACTTTGCGCTTAAAAGTACTGCAACTTATCCATCGAAATCGTCGAGTTAATAATTTAGTCACTGTATCAAGCACTTTCACCAGAAATACATAA

>Tadh124_evg712 type=CDS; aalen=369,42%,complete-utrpoor; clen=2594; strand=+; offs=390-1499; evgclass=main,okay,match:evg1722407,pct:99/100/.

ccaccATGTCGAAGTTCGAAGGTTATAATTTAACTAACCATTTTACGACGACTTCATCATCACTTGCTGCTAATTCTTCAATACTTCCATTTACTTGGTCGCTACATTATAATATTCCGATCTTAACCCTATTCTTTAGCATCGTCATGATTTTTGCAGTGACTGGTAATCTATTCTCTCTGATAACTATCTATCGTTCGGTCTCATTGCATACTATCAATAATATGTTCATCTTTAACTTGGCCGTCGTCGATTTACTCAGTAGTCTATTAGTAATACCTTTAGGTTTGGCCGTTGTCAGTAGTCAATCACATTTCAGTAATACATTATGCCAAATTCAAGGTTTCTTGCTTATGTTTTTAAATACCACTTCATTAATGACAACAACTCTCATCAGCATTGATCGAGTTATTGCTGTAACGAAGCCATATTATTATGCAGCTATTGCTAAATTTCATGCTATGATCGCTTTAATAGCGGGAATTTGGATATTATCCATCCTCTTTGCAACACTACCTCTACTACATTTACAACATTATGGCTTCGGCTATTATACTCTTGTCTACATCTGCTGGATTAATATCCACGATTATGACGCAAATTTTGTTATTGCTATCGCGGCGATCTGTTTGATCTTGTTTAGTATGATGGCCATCATAATAAGTTATACCATCATATTTTGTATCGCTTGTAATAAAGTTGCCAACCAATTGAGACCAAATGGTTATGGCGATGTGATGAAATCAGTTCGTACCACAGCAATTATTGTCGGTACGAATTTACTCTTTTGGTTGCCATTTGTTACTGCCTGTATTATCGAAATTAAAGATGGACACTACCATCGGAGAGATCCTGACTTTGATGAAACATTATTCAAGGTAGCTTATTTAGCACCCTTTGGCAGTGCTGCTAGTAATCCAATCATTTATATTACCACTAATGGCGTCTTGCGCCTTAAATTTCAAAAATTATTTCAATTTAAAACAAGCTTGCAGCCTCTTCGAAAACGAACCGCAATTTCTCCAAGCCGACGTAACAACTCTACGGCTGCTTGTATAGTAGAGCAAATTAACAGTCTTAACGATTCGTATCGTCCAGATGATGAGAATCAATAG

>Tadh126_evg728 type=CDS; aalen=337,70%,complete; clen=1442; strand=+; offs=350-1363; evgclass=main,okay,match:evg1420869,pct:99/100/.

ccaccATGATAAATTTAACAAATAGTTCATTTCCTTACATGCCAATCTTATCCTACAAAGACATTTTCGTTGTTACAATATTGGGAATAATTGCTTCGGTCGCTTCGTTTGGTAATCTTTTCAATCTCTATCTCATCTATCACTCAAGATCATCACAATTGGTTGGTACTTTACTTCTTGCAAATTTAGGTTTAGCCGATTTATTTTCTGGACTTGTTGTTATGCCGTTAGCTATTTATAGTTTCATCGTACAAGATACTATATCTCCGTTTCTGTGCCAACTGTGGGCAGTATTAGCTAGCCTAAGTATTGGTGTATCCTTTTATACTACTGCAGCCATTAGTATCGACCGCTGTATCGCCGTTGCTAGTCCATTTTATTACTCTCAATCTGTTAATTTACGTGTATTAAAAGTCCAGATTGTAATATTATGGCTAATAACTAGTGTAATCGCCATTATGCCAATGCTAGGCTTGAAAGCCTATGGTATTGGACATTACAGTTATATTACAGGATCGCAACAATGTTGGGTTGATTTTACGGACAGAGATCACAATTATTATATTGGATTGGTTTTGTTGTCGATACTATCGTGCACGATAGTGATCATTGTACTGTGTTATAGTATTATATTTATCATTGCTTGGAGAAAAGGTTTACAAAATCTATCAGGCCATGGTACTATCAAAAAATCGATCCGAACAACTGGTTTAATTGTTGGCACCAGTCTAATGTGTTGGATTCCTCTTATAACTATATCTTACGTCGAATTTATAGACTTCATGTTACTGATATCAGATGACATAGTATTAGCAGCCTATCTTCTTACATTTTGTAGCTCCGCTATTAATCCTGTCGTTTACTCATTGACAAATAGCGAACTTAAGCGTCGTGTCCGACTAACGTTGCGAAGGACAAACGTAATTGCATTTTCACAATCAAAGCGTAAAGGCAAAGTTAACCCGATCGGAATCGATTCTGTTATTGGTAATCTCCATAGCTCTGTAATAGTCGGCTAA

>Tadh127_evg811142 type=CDS; aalen=352,51%,complete-utrpoor; clen=2076; strand=-; offs=1669-611; evgclass=noclass,okay

ccaccATGGTCACAATTAATCAAACGAATTCTACGATAGCAAATCCAGGTATCAATCAAATTTTATCTATTTCTTTCATATCGATTGTTATTCTCATAGGAAGCATAGGGAATTCTATCAACATTTACTTTATTTACAAGATACCATCATTACACAGCTACAACAATATGCTTATAGCCCATCTAGCTGTGGCTGATTTCTTGCAAGCAACTTTAACATTACCCATGGCCATTGCTAATTCTTTATTAAGACATGCCATTCCACCAGCTATGTGTCAAGCGTTCGCTTTTATACTCAATTTTCTTCTCGCCATTGCATTAGCAGCAACGGCAGCCATTAGCATCGATCGCTGCCTTGCTGTTATTTATCCCTATCAATACCAGGCTAAAATGAAAACGAACTACATTATCATGATTGTCATATTCATATGGATACTAGCAGCAGTTTTGGCCGGAACGCCATTACTTGGTTTACAACGATACGGTTTAGGAGAATATACCTTCATCGCTGATGGTCTACAATGTTGGTTTGATTTTCAATATCGTCAAAGAAGTAGTATTATATTTATCATTTTATATATCTACTTGCTATTAACCATCTTAACAACCCTAGTTTCCTATTTGATTATTTTCTGTATTGCCTACAATAAAGGTATCGCTGACATATCTGCAGTCGGTTATGCATCATTACGACGCTCTATCAGGACGACTGCATTAATTGTTGGCAGCAATTTTGCATGTTTTATACCAACATTAATATCAGTATCTATAAGTTATTTCTCTCAACGTGAATTAAATCCAGGTTTAACCGTTGCAGCCTACCTTTTATCATTCTGTAATTCAGCGATTAATCCACTTATTTATGCCTTCACCAATGGAATCCTTCGCCAAAAAATTAAGCAGCATTGTTGTTGCTCAAAGAAGCGATATTGTCTTACAAAATTTGGACTATATAGAACCTCAAAAATTACACCAAGCAATACCGCAGCTGGTCAACCAGCTGCTAAAAGCGACAGCGAACCAAGCTACTGGCAAACTGTTATAAATGATTATGAATCATCGTAA

>Tadh128_evg87029 type=CDS; aalen=353,81%,complete; clen=1305; strand=+; offs=220-1281; evgclass=main,okay,match:evg892328,pct:99/100/.

ccaccATGTCAAGGATGGATCCTATCAACGTGCTAGTTAATGTGAGTAATCACTCGCAAAATCTTGCGAGTAATATTAGTCAGAACCAGTCTTATGCGAATGCTCCTGTCAACTTTATGTTCATGATTCCGGCATTATCCATTATTAGCGTAATTGCAATAATTGCTAACATATTTTGTCTTACTACAATCTACCATACGCCATCCTTACATGACATTAGTAATGTTATGATAGCCAATTTAGCCATAATTGATCTTTTAACGGGGATGACAGTTATACCTACGATTATTACAGTACTTTTATTAGGCAATGAAACTCCATCATGGTTATGCAAACTTCAAGGATTTTTAACGGTTGCCATGTATGGTGCTTCATTATTTGCAATTGTTGGTATAAGTATCGATCGTTTTCGAGCCGTTACCAGGCCGTATCATTATTCCTCTATAATCACCAAGACAATCATTGTAAAATTGCTGATGACAATACCTATCCCCATCATTGTTGCATCCTTGCCATTATTCAATTTGGTCAATCATGGATTTGGCAGCTATATAAAAGAATTTGTTTGCTGGTCCCCCGCCTGTAGGCACAATCAACCAGCTTATATCTCAAGTGCTGCCACCGCTGTATTCATAGTTATATGCATTGGAATTGTTTTATTCTGCTATATCACAATTTTCATCATCGCCTATCATAAAAATAAGAAGAATATCCATCTAAAGCGAAATATCAATATAAAATCTATACGCACAACAGCATTAATAGTCGGCACCAACCTCATCTGCTTTTTACCCCAACTTAGTGCAATTACTTCGGCTCTCATTAATTGTAATAATAATTCTCCGGAATTTTTTCAAGCTGTATTTTACACTATAACACTAAGCAATTCTGCACTTAATCCTATTGTTTATATCTCGACCAACTCAATTTTGCGGAGAAAATTAGTTCAATTAATGCTCGTTAAAGCTTTGAGTAAAAAAGATCCTTACAGCCAACGCGATCATTTACCGCAAAGTAGAAGTCTAGAACTTAGTAACAATACTTCAATTTTAACAACAAGATTTTAA

>Tadh129_evg93544 type=CDS; aalen=344,83%,complete; clen=1244; strand=-; offs=1212-178; evgclass=main,okay,match:evg595,pct:100/90/.

ccaccATGATGGAATCCATACTTTCCGTAGATTCATCTTCAATCTTGTCTGTGCTTGGATCCAACGCAACAAAAGGTACATATGGAATGCAGATGGGTCTATCCAATTCCATTTATTTAAATATCGTTTATGCAGCTATAGTGGTCCTTAGCACTCTAGGCAATGTCACGTATTTATCCGTGATTTATCGATGGCCTGAATTAAGAAGTACCAACAATATTTTCTTAGCGAACTTAGCATCAGCCAATTTATTAATGGCATTGGTTGTAATACCTTTCCACATCATCGTTAATGTAATGCAATACTCGATGACGACTTTGGTCTTATATCAAATTTTAAATTCCTTCTTGCTGTTTATATTCGGGATATCTTTAATCATGATGATGGCCGTCAATATTGATCGATGCTTTGCCATTGCCTCTCCGTATAATTATACTGGTATGGTGACAAGAAAAAAATTGGTCAACCTATTAATTTTAACATGGTGCTCAGCTCTAATTATTGCCGTAGTTCCGTTGATCTTTTTCTTAATTATGCTAGATCGCATTCCACAACAACGATCGCATTCGACAGAACCTATCGAATCAGAAACTGAAAAAATCGAATTATTTCATACAACCTATCGTTTGTATCTTATGATATTAATAGCCTTATTATTCTTAACTATATGTACTGTCTTAATTTGCTATAGCATCATTTTCATAATTGCCCTTATTAAAGGTAAATACAATACTAAAGTAAAAGCATTTCCTAAACATGCTATAAATATCCGTAAATCCATTGCTATAACACTTTTTATTACTATAGCGCATCTTGTGTGTTTAATACCATTAGCTATGGTATTACTGGTACATTGTGAGTATAAATCGAACGTGTTATTCAGAGTATCGATTAATCAAGTTAATTTAATACACTTAATATGTTTAATTACCACTGCACTTAATCCATTGGCTTATATTATTACCAATAGATCACTTCGTAAGAAATTAAAGTATCTACATCAGTATTACTGCAACGCTTCCGATGCCATATTGGCATAA

>Tadh130_evg97324 type=CDS; aalen=328,50%,complete-utrpoor; clen=1972; strand=+; offs=622-1608; evgclass=main,okay,match:evg1733964,pct:98/100/.

ccaccATGGATTTAACGCGTCCCCCCAAAATTTATAATTCTTCCAATCATCTCGACGACTATAATCATAAAATAACAATCCAGATTTCGTACCTCTTTTTGATTACGATTATGGCCGTTTTAGGTAATTTATTCAATTTATTCCACATCTTTCGTTCATCAACTTTACGAGATATGAAAAATACGATGCTAACTGGTCTGGCCTTTGCTGATTTATGTACAGGAAGTATAGTACCACCTTGCATGATTTTGACATTGATTTATCAATTAGAAGATACAACATGTCAAGTTCAAGGTTTCGTTGTTACATATTTTCATGGAGTATCACTGGTTATGTCAGCGGCCATAAGTATTGATCGATGTTCTGCTATTTCTGATCCTTATACTTATTTAGCACATTTACAAGTTTGGCGCTACAGCATCATTGTAACTAGCATTTGGATAATACCTTTACCTTTTGCTATAACACCTCTATTGCCGTTTCAATCATTAGGATTTGGTAGTTATGAACTTACTAGTTTATGCTGGCTAAGATTAAGTACAAACAAATATAACTACGTTGCTCTTGCCGCTTTAGCAGCTGGAATCTGTTCAGTTGTCACTATAATTATTTGTTGTTACGTAATAATTTTCTGTATTGCCTACAGGAAAAAAAATTCCCATATCGATAGTTATGGAAGTATTAGAAAATCGATTCGGACTGCATTTCTTATAGTTGGCTCAAATGTTTTCTGCTGGTTGCCATTAGGAGTTACATGGTCTGTCAGTGTCATAAAATATTTTGTATTTCAAGGAGAAATTAGGATCAACTCACAATTAGAAAGTGCAGTATTATTATTACCATATTCTAATGTTGCCATAAATCCGCTTATTTATGCTGGAACAAATGAAATTTTAAAGGATAGATATGCCAAATTTGGTGTTCGCATTTATAACTACGTATCGGATCGATTACCGTTGCCTGCCAGTAATGCTATAAATGACGCCGCTTAA

>Tadh131_evg101767 type=CDS; aalen=411,55%,complete-utrpoor; clen=2245; strand=-; offs=1464-229; evgclass=main,okay,match:evg1365777,pct:99/100/.

ccaccATGGAAGATAATTTTTCATTAGTTTATAATTATGGTAATCTGACAGACGATCCACTATCTAATACTCAAATTATTATCTGTTCAATTATATTAACACTAGCAACGTTACTATCGTTAATTGGCAACAGCTTGGTGATTACAGTTATTATTCGTAATCGTCCACTTCATACTAATGCTGGTATGCTTATGGCTAATTTAGCAGTGGCCGATTTTTTAGTTGGTTTAGTTATCATGTTTCCTACCCTAATTGCATTTATATGCCAAAAAGATATTTACCCTCGGATTGCCTGTGTTGCTCAAGGATCAGCAATTACAGTTTTAGTTGGTGCTTCGACAGTGACTATATTAGCAATTACTGCTGATCGTTTTATCTCCATTTTTCTATCCTTGCGTTATAATTCAATAGTAACAACAGCTGTCATTATTGGTACTATTGCTTTCTCATGGCTATTTGCTACCGTGGCATCTACTATTCCATTGCTTGGTTTACAGAAATATGGCCTCGGTCGTTATGAATTTCTTCATGGAATGTCCTATTGTTGGTTAGATCCATTGCAAGTTCAACAGAATAACGTTATAATCTCTATAATTGGTTTCTACGTTGTCTTAGAAATATTTATTATTACTACATGTTATTTAGTCATCTTTATTTATGCTCGCCAAAGTGTACGTAAAGTAATTCATTTAAGTGTTGGTGATTTACCTAAAGGGCGGCCTAAAATAACCAAAACAACGAAAACTGTCATGATTGTGGTAGGTGTATTTATGTTTTGTTGGATGCCGTCATTGGTCAATAAGTTCCTAATTATATTCAAAGTTAAAACATTATCCTTATCAGTTCGATTAATCTTCACATGGCTAACATTTTTCAATTCGGCTGCAAATCCAATCATCTATAGTATATTTAATCGTAATTTTATTACTGGTGTACGTAATTTGTTCAAGAAAAGTATGAGTGAAAGATCGATTATAGTGTCTATGTCGGAGCGGAGATCAAGATCATTTCGGCGCCGTTCTCTATCAATCACTAAATTTAGGAACTCGTTACTTCCAATGCATAGAAGTAATACTGTCCAAGATGTTCGCTTTCCATCTTGCAATCAAAGTTTACAAGTTCCTAAGATTGTTATTGCCAAAAGTAAATCTACTGAAGCTCTTTTTAAGCGCCGACGTGATAAAATTGTTATTCATGCTACATTAAATGAAGTTCCCATCGAGATTGAAGATGTTGCTTAG

>Tadh132_evg15166 type=CDS; aalen=365,48%,complete-utrpoor; clen=2262; strand=-; offs=2001-904; evgclass=maina2,okay,match:evg15165,pct:100/99/.

ccaccATGAATAACCTAAGTCGACCAACTATGGATATTAATTCATCGTATGATAAAATTTCCCACGACTACTCCTTCAATAACTCCGATGACACTGACTTAGCTATTTTATTACCTATTTTAGTAACTTTTATCGCTATTATCATGACTACTGCAATGGCTGGCAATCTCTATATCTTTTATGCCATTTATAGATGCCGTAGCTTGCACAAAACATCGTCGTTTTTGATTGCAAATTTGGCCGTTGCAGACATATTAACAGCAATAATTGTTTTACCACCTATGGCCTATTCGGCTATTCAAGGTAAAATGCTATTTAATGATACATTTTGTCAAATATCAGCATACATTAATTATATCATATTTGGAGCTTCATTAAATACCAGCATTACCATCACGATGGATCGATTTGTAGCAGTACTTTGGCCTTTCCGATATGAGCGTATAACAACACCATCTAGGACAGCTTTTATGATTAGTACGTCATGGCTATTTCCATCAATTGTAGCTATGCTACCCTTATTATCATTACAACGTTATGGTTTAGGACGATATAGGTATACAAATAGTACTTATTACTGTGGTATTGACTTACACGATTATCAAGATAATTACCTATTTAATGTAATTTCAGCCGCATCAATTTTAAATAGCGTAGCTGTGGTAGCTATTGCTTATCTCTTTATTTTTCGCGTTGCATATAAGAAATGGGTAAAAAGTCAACAAAAATCGATAGATATTAAAGGAAACAAAGAAGCAAGTTTGCTTAGAACAGTACGTACAACGGCTTTAATTGTTGGTACTAATATCTTATGTTGGTTACCCGCAGTTATACAAGTTCTCATATATACCATTAATCCTCCTAATCAACTTGCTAACAAAGTGGATGTTTTAGGTATCATTTTCGTATGGCTACCATATACAAATTGTGCTGTTAATCCAATTATTTATATCTCTTCTAACTCTCGGTTACGCAACCTAGTGATTAAGCAATGGGAATATTATCGAAGAAATATAATGTGTTTTAAATTCTGCCATCCTCGCAATAAAAATGTAATTGGTCCATGGTTGACATCAGAACCATCGAATTCCTTTCATTTATAG

>Tadh133_evg1659776 type=CDS; aalen=380,58%,complete-utrpoor; clen=1962; strand=+; offs=567-1709; evgclass=maina2,okay,match:evg19761,pct:99/100/.

ccaccATGGATTCTATCAATTCAACGCAATCTTCATTGCCATATTATTCATCAACCAACTATCTGGCTATTGTTATTTTAATTTTGTTAGCCATTGCCTGCATATTAGGCAATTTATTATCTATTATAGTTATCTATCGTAATAAGAATATGCATGATCCGTCTGGCTTACTGATGGCCAACTTAGCCACTGCTGATTTATGCATCGGATTAATATTTATGACTACGACCGTTGTATACACATGGAAACAGTCTTATATGTTTAACGAGTTAAGTTGTGCAATGCGTGCATATCTTATTACTGTTATAGTCACTGTCTCACAATTTACTGCCGCTTATATTGCCATAGATCGCTGTATCTATGTCTGTAAGCCATTGCATTATCCAATGTGGATTACAAAATGGAAAATGGGATCCCTTATTATATTTACATGGATATTAGCAAGTTTATTTGGTTCTATTCCGTTAATGCAGTTGGAACGATATGGCCTCGGTCGTTATACCTATGTCAGTTATATGTATTCATGCTGGCTAGATATAAAAAAATTTCAACAAAATGGCAGCATCTTATTATTGCTATTTAATACTTTTATTGGACTCTTGCTTATCGTTTCAATATGTTACATTATTATCCTGTATAAAGCTAAAAAAGTACATACCACTATTCAGCCATTTAGAAGGCAATTCAATATACGGCAGCAACTACCGACACGAATGGTTAAGCTATCATGGATTAATAAGCCTACTAAAACGTCGGTTATACTCGTTGGTGTTTATTTGATTTGTGCAGCGCCAACAGCTATATTTGGTATAACTATGACATTAAATTTTGAATATTATAATGTTAGTGATGCTTACTATCGCGTTCTACTTTATTGGTGTATGTACGCTAACGCCTTAGCCAATCCATTTATCTTTGGATTTTCACATCGAGATTTCCGTCAGACATATCGATCGTTGTGTTGTAAAAATCCTATGCGATACATAATGCAATTTCTACCGGTTAAGAAGCAGCCACCCAGTGGAAGTTTATCGTTTACGATTGTGCGATCATCCGATAGTAACCATCATAGTATATCGCATTATCCAGCAAGCCGCTCAACGAGAGTGAACGCTGCTGGGGGGAAAAATTTATTTCTTGACTTTTAA

>Tadh134_evg1747753 type=CDS; aalen=358,40%,complete-utrpoor; clen=2656; strand=-; offs=1869-793; evgclass=maina2,okay,match:evg19748,pct:99/100/.

ccaccATGAATGAATTAAAAGTCAAGTATACTTGCGAGAATTATAACAGCACCTTACTCATTAATATAACTTACGAGTATATTCCTACCGAAGAGGCCTTAACATTAACAGTAGTTGTTTTAATATTTTTACTAACATCACTCGTTGGCAATATTGCTGCTGTTGTAGTAGTTTATCGTAATCGCTATCTGCATGATCCCTCCGGTATACTAATAGGTGCTTTAGCAGTTTCTGATATTGGCGTGGGATTTTCTAGTGTTATTTCCACCATAATCTATGATAATCCTTGCTCAAGACTAAATCATGCAGTTTGTGTATTTCTCGCTTTCATGTCAAGTTCATTTATCGGCTCTACATACTCAGCTATAGCAGGTATTACAGTGGATCGTTGCATTGCAATCTGTAAACCTTTACATTATCGACAATATGTTACAAACACCAGAATAATTCAGATTCTTCTTTTCAATTGGTTGTGTATTCTTGCTCTAGCAGCTCTACCGGTTATGGGATTACAACAATATGGTATAGGAAAATATGAATATGTTCCCTATTTAACGTCGTGTTGGATTGATATTACTGCATATAGTAAAAATTACTTGGCTATTATTGTTTTTTACTTGCCTGTCTTCTCAATATTGATACTTGTCATGATATGCTATGTTATAATTCTTCTCAAAGCAAATAGTACTTTTAAACGAATGGCAATTTCAGGTTGGATCTATAATGAAAATAAGCAATTACTGAAGAAATCAACGCGAACGACTATTATTGTTGTGGGTGTCTTTTTTATATGTACAACTCCAGGCCTTGCATTAACATTTGCTACGCTAGTTAATCAAGGATCTGTTGTGTCTGGTAAAGCCTATAAATATTTATTAAATTGGGGAATATACGTCAATATTACGGTTAATCCTTTATTATTTGGTTTCTCTCATAAAGATTTTAGACATGGCTATCATGAAATATGTTCACATTTTTACTCAAAAAAATCAAAAATTTCTGATATCAGTCTACCGATACGACCATCCGTCAATCCTTCAGTATTATCGGGATATAATTTGGTTCAAAATAGTACATTTTAA

>Tadh135_evg19755 type=CDS; aalen=336,52%,complete-utrpoor; clen=1930; strand=+; offs=507-1517; evgclass=main,okay,match:evg19756,pct:100/100/. was not possible to synthesize, complicated gene

ccaccATGGCATTGTATCCTAACTGCTCTCCTCAAGATTATTTCCTCAATACATCCTTCTATCGCGCTGAAGAATCGCCAGTAACAAAAAATATTCTTTTAAGCATACTATCTACACTAGCTGTCTTAGCCATAGGCGGTAATTCGCTAGCAATTCTTGTCGTCTATCGTACTCGATCATTACACGATCCATCAGGTATTCTTCTTGCTGGATTAGCATCGATAGATTTAGCCGTAGGAATAGTTTGTATTTTAACTTCTGTGATATATCACTGCTTGGACGGATATATCATTTCCGAGACTAGTTGCATAATCCGCGGATATCTTATCACAACTCTTATTGGCGCATCTTATAGCACGGTTGCTCTTATTGCTCTTGATCGATGTTTTGCTATACATAAGCCATTACATTATCGACTTTATGTTACTAATAAACGAATACTTCAAGTTCTTATTACGAATATAGTATCAATAGGAATATTATCTGCTATGCCATTCGCGGGATTACAAGATCAAGGTTTAGGCCAATTTGAATACGTCCATTATATATCTGGTTGTTGGTTAGATATTACTAACCATCCTAAAAATTTTCTTTTTTCTCTCTTTATTGCACTAGCAGTCAACTGTATATTAATGATAGTTATTATAGCTTATATCGTGATACTAATTGAAGCTCAACGTATAGATCGAAAGTTAGTTCTCTTAAAATCGTCGTTTGCTGAAAAGAGGCAATGGTTAAGAAAATCGACACGAACAACTTTACTGTTAATCGGAGTGTTTTTAATATGCACTATCCCATCCTCAGCTTTTACACTATTAGCTATTATCCATTGTAAACCTATTGCGTCTCCATTAACGTATAAAATTTTAATTCACTGGGGAATTTATATCAACACGACAATCAATCCATTTCTATTTGGATTCACCCATAAAGATTTTCGGCAAGGTTACCGCCATGTATTTTCACGATTTTCATTATTGGGAACAAGAGTCTCTCATGTTCATTTCAATCAGTAA

>Tadh136_evg19758 type=CDS; aalen=337,61%,complete; clen=1651; strand=+; offs=408-1421; evgclass=main,okay,match:evg1758788,pct:99/100/.

ccaccATGCAATCCGAATGCAACGAATGGAATCAAAGTCATTGCTTCTGTAATCAATCTGAGACATCATTTCATATCACAAAGATTATCTTACGATGCATAACGCCACCGGTATTTATCTTAGCTTTAGGTGGAAATTTCTTATCCGTTCTAGTTGTTTATCGTAAGAGAAGTTTACATAATCCATCAGGTTTCCTTATAGCTACTTTGGCGGCGACAGAGATGGCTACAGGTTTAATCGTAACCTGCTATTTCTTTGCTTTTTCAGCATTACATTGTGGTATGTTGGAAAACTTTTGTATCGTTTTAGCCTATCTATCTACCACTTTCGTCTGTTTCTCACATATAACAGTAGGATTGATCTCAATCGATCGATACATAGCCATCACTATGCCACTACATTATCCAATGTATATTACGAATAAACGACTCAGTAGGATATTAATATTATCCATTACCCATAGCTGCTTCTGCGCCTCATTACCATTAATGAATTTGAACCAAATTCAACTTGGCCGCTACCAATACGTCGAATATTTATCTCTTTGCTGGATAGATATTACAGATTTAAGTGAAAATTTCATTTTCTTAGTAATAATGTATTCTGCTTTACTAATGGTAATATGCATCGTGGTTTCTTGCTATGTTCTCATTTTCGTTAAGGCAAATCAAATCCATCGTAAACGTCGAAATCTTGAATTGACAAAGAGCAGTGTAAGGCATCTGAGCAAGAAGTTAACTCGTACTATCCTAATCGTTATTACCGTATATTTAATTTTTATTATTCCTTCAACAGCTACAATTCTGATTACTCTGAGCCATCGTCGATTGTTCTTATCCCCAACCATGTCTAAAATTATGACTTTTTGGGGCCTACATATCAAAATCATTATTAATCCGCTAATATTTGGATTTTCTCATCGAGACTTTCGACAGAGCTATAATCAATTATTTTATTATTTAGTAAAGCGTAGAAGAACTAGAATTTTCGATCAAGTGTTGCGTCCATCGATACAATAA

>TriAdh137_evg2327 type=CDS; aalen=505,38%,complete-utrbad; clen=3946; strand=+; offs=246-1763; evgclass=maina2,okay,match:evg1101169,pct:99/100/. was not possible to synthesize, complicated gene

ccaccATGGAAAGCGGCAATTATAGCATGGATTATAATACAACTATTCTCCTATCAGATTTAACATCGATTCCATCATTATTATCTAATCAGAAGCAACCAATGCAAAAGCAATCACCATTAGATTGGCGGATTATCTTTGGTGTCACTATCTTTGTAACTGCTCTAGTTGGCAATTTAATATTAATGGCACTGATACTATTTGATCGTCAACTGCGTAAATTAGAGCATGTGCCTATTGCAAGTTTATGTGTTGTCAATATCTTGACAGCTGTTATGATTATCATGCCCTTTTTTATGTTTTTAATCCATCGCAAATGGGTACTTGGCCAAGCATGGTGCTACTTATGGAATCCTATGGGATACTTTCTACCATCGACAGGCATCTTTCATTATATGATTATTTCAATTGATCGTTATCTACGAGTTACTCGTCCATTAGAAGTAACTAAAACAAAAACCAAGCGCCTACTGATATTGCAACTGATATTATCATGGATTATACCAGCGATTGTATCTTATTTGCCTGAGATGATATGGAGGCAATTCGATTTTACTTATTCCTATCGATGTTATTATAAATATCATAGGAACAATATGTTAATTTATAACAGTGTAGTTGTTACAGTTTTATTTGGCCTACCATTATTCGTAACAACGGCTATTTATATTAAACTAACTCGAATAGCATTAGGGCATAGTCGTACAATAAAGCTACAACGTCTATCGATTGCATCGCAACAGTCACAAGAAAAAAGTACCAGAAAGTTACGACATCAAAGTAAAGGAGAAATTATTTTAGCATTTCTTTTATTAGCCTTTGTAATTTGCTGGCTACCAACCTTAATTTATGCTTTAGTCTCTTCCATAACTTATCCAAAAGATCCCAGTAGAAAGATACCATGGCTTTATCCAATGTTGCTTGTTATCAACTTCTGCTATCCAACTGTTAATCCTATTATGTTTGGGTTGTTATATAGACCGGTTAGAAGAGCCCTAGTGAAGCAATATTGTAAAGTTTGCCGATTTAATACATTCATGGACCGTACAACAACTATGATGAGTTCAATATCACGATCCAGTATAGCAATGAAAAATAGTAATAGCTTAAGTAACAGTCAAGATAGACGAGTATTACGTCGACTAAGATCATCTAATAGTAGTTTTAATCCATCTGATACATTAAGTAGCATTAGAGATCATAAAAGACCAAGTTCATCTTGCAAAGATAACATTTCTTACAGCAATAAATCATCACCAGGGCCTTGTAGTCCTACAACTGCACTATATCGACCTTTTCCCAAGACAATTATTGATCAATCTAGCATATATATCCGTGAAAATGTCGGAGAAAATGTGACTAATCGTCTACAGTATCATCAACAGAAAGCTAATCGTCAAACAAATGGGAATCACTCGTATAAAAATTCTCATTCTTCATCGACATCTTCTGGTAGCATGCAATATGTAGATATTAACGATAATATGTTCTATAATGATTATAATCAAGAAACGAATGTATGA

>TriAdh138_evg2834 type=CDS; aalen=323,40%,complete-utrbad; clen=2394; strand=+; offs=1362-2333; evgclass=maina2,okay,match:evg101819,pct:99/100/.

ccaccATGGATAGGCTAGCTGAGAGTAAAGGCCTTTTTTTGAGCAATAATATTACAGCCGTGCACAGTAATTATGCTACTACAACGCTATTTACGGTGATGACGGCAATTATATTTGGAAGTACTGTGTTTGGTAATATTATGTCATTATTTGTTATCTACAAAACATCATCTTTATGGGATAATTCCGGTATTTTATTAGGAAATATGGCAGTTGCTGATTTAATCGTAGGCCTCATTTGTATGCCGACTGCTTTTATTTCAATGTGTATCCAGCGTAGTTACCTGGCTCCAATCTTATGTGTTATAACTGGTTATATTAACACTGTTGTTATCGGAACATCATTACAAACGTTAGCTTTTATTTGTGTTGATCGATGTATTTCAGTTCTCAAGCCCTTACATTATCATATGTACGTTACTAATAATGTATTAAGGCTTATGCTAGTAATATCTTGGGTTTTTCCATCTGTTGTTTCAATAATACCTCTACTAGGACTACAGCAATATGGTTTAGGCAAATACTGCTTTGTCAACTATTTATCGACGTGCTGGTTAGATGTAATTCAAGTCCAACAAAATAGAGTGTTTCTATTGATGATTTATTCATTAGTGGTAATTATTCTTACTTGCATTGTAATCTGTTGCGTACTAATACTTCTAGTTATACATAAATTAAGAAATACGGTCGCCGATATCTGCACATCATCGATTCATGCTCGGGCAGTTTCTAAGTCAACTGTTACTCTTATGTTGATTGTAGGAGTTTTCATCTTATGCTTAATGCCTGCTGTGGTTGTTACTTTTATTACTTGCTTTACTAACGTTCAATTCCATGTAACTATTTACCAAATGGCTATTTGCTTAACTTATTTGAACTCTGCTATCAATCCCATCTTATACGGATTTTCTAGTCAAGATTTTCGCAATAGTTACTATACATTATGGAAGAGACATTTTCGTTTATATCGCTTTTAA

>Tadh139_evg374227 type=CDS; aalen=362,55%,complete-utrpoor; clen=1971; strand=-; offs=1570-482; evgclass=main,okay,match:evg1123651,pct:100/98/.

ccaccATGAGTATCGACGATACGTATCTTCGTGAAGCAACATGGAGTAATATTGTTATTACTGTCATATTTTCATTTCTATTTTTGTTGATTGTTATCGGTAATGTTTTAGCATTATTCGTTATTTATCGACGAAAATCATTACAAGATCCAGCAGGAGTAGCCTTAGCTAACCTTGCTGTTGGCGATCTCTTAACAAGTATCCCATTACTAGGCTCGACCATTATCTTTTGGATCGATGATGCTCATATTAATCCGTTAGCGTGTAAAATATTGGCCTTTATGTCATTTTATCCTTTTACAATTATATTATGGACTACAGCATGGATTACATTTGATCGTTGTTTTGCAGTCTATCATCCATTTAAATATCATCGTATCATTACAGTTAAGACAGTTGTTAGTCTACTCGCCATCGCTTGGATCCTTCTATTAGTATGGTGTATGATTCCGTTATTATCGTTGGGCCATGGTTTAGGGAATTACAACTATGTTAATTATACCCATTCCTGTTGGGTTGATGTGGGCAATTATCGGCAAAATGGAGTCTTTCTATTAATTACTTATTCAGCAGTTTTAGTACTTGTTATTATTGTGGTATTATGTTATAGTATTTTGTTAATTAAAGCGCGACTCGTATTACGACGCATAACTACTATTCCTACTTTTCACTATCCTGCGTCATCATCACTGCCGGCACCAAGTCATAAATCAGTTAAGACAACATTAATTGTTGTTGGTGCATCATTAATGTGCATTTTACCAGCTGTTATCATCATCTATATTACTATCATGTCACATCGTCTTATTTCAGAGCCGTTAATCTATAATTTATTTTGCTTCTGGACATTTTATGTAGATGCTGCACTCAATCCATTCATATTTGGCTTCTCTCATAAAACTTTCCGTCTTGGTTATCGTAAATTGTATCGTCATTTTATGAGTAAATTTATTACACCGCATCATCGAATATATCCTGCAATACCGTCTGCTCCAGCTTCAGTATATAACAGTATCCAAAGTAAAATGAATGATATAGCTAGTAATGGTTACGATTGTTCAACAATCTTTAATTCTAATTTAAATAACGAATAA

>Tadh140_evg370902 type=CDS; aalen=374,83%,complete; clen=1341; strand=-; offs=1304-180; evgclass=main,okay,match:evg1315926,pct:99/100/.

ccaccATGAAGGGTTGGAATTGGCAAAATACGCCTAATGGAAGTGTTGATAATTCTACAACTCAGCATAATTATTCACCAATATATACATTTCAAGCTATAGCGTTGTCTGTCGCAGTTATTGCTGGATTTCCAGCTAATATTATCACCTTAATAGCTTGTTGTTACAGAAATCGTCTTAACGTTGTTTCCAATATATTTATAGCTAACTTGTGTATTTATGAATTATGTATGATATTACTTGATTTTCCTCTATCTCTAGTGAATGCTGTTAATGGCCGACAAATTGTAACCAGCTCACTATGTAAATTTCAATGTATAATTGGTACCATGTGTTCAGCCGGAATTATTTTATCATTAGTTTGTGTTACCATTGATCGCTACTTTACAATTGTAAAGCCCATGAAATATGGTATGATTATGACTATTCGTAAAGCTGTTACCATGATTATATGTGTTTGGGTTTATGTTTTCTTGTTAATTCCAGCTTATTATATACCAGAAGTACAACCTACTTGTTTATATTACTACGAAATGTCCAGTTGCTTTACAAATTGGGCTCAATCGCCTATCGTTGGATATTTTATCCTTGCAACATGTTATGGCACGGCGCTAATCGCCATGATATACACGTACTGGAGGATATTCGTGATTGCGCGCCACCATAATAAAGTTGGAGTTGAACAGTTGAAAGAATTAACAAAGAATGGTTACACTAACCGACCTTCCATTACTCAAATGAGAAGAAAAGGATATAAAACTATCAAAGTTGTTATCGTTATCCTAATTACATATATTTGTTGTTGGACGCCTATAACCATATTAGCAGCTACAAATATTCAGAAATTTTCAGATATGCCACCTTGGGGTTTAGCGATTGCGAATTGCTTGGTTGTTATCAATCCTACAATTAATCCAATTGTATATGGTATTATGAGTCAAGAGTATAGGCATAGCTATCAAAGAATATATCGTATAATTCGAAATAGTCAGACAATACGTCGAAGCTTAATTATTCCATCACAAGCTTCATTTGCAACGAATCAAAAGGATGGAAACGGCAATTTGTCTTTATCACATGATGTCATAACGACAAATCACGAAAGTAGAGAACAAGAAACAGTAACTTGA

>TriAdh141_evg1041245 type=CDS; aalen=449,37%,complete-utrpoor; clen=3638; strand=-; offs=2666-1317; evgclass=maina2,okay,match:evg374057,pct:99/100/.

ccaccATGCAGCAACAAATTTATCTACGATCTCCCAATATCAATATTAGCGATCCAATCCCTACGGGTGGAGAGTATGGTATCAATCAATCTTCACCCTATTGGATTTTACAAACATCAAGTTACATTGGCATTAAATTAACAATCCATGTCTTATTACTTATGATCATACTCATCGGTAATATTACAGCGATCGCAGTGGTTAAATCGACCCAATTATTACACCGTCCAACGGGTTGGTTTATCCTCAATTTAGCTATAACGAACTTTCTCTCTGGTCTATTTCTATCCCCTATCATTATCATTACTACAGCTACAGCGAGTCGCTGGTCGCTAGGTGATGGCATGTGCCAATGGGTTGGCTTCGTTAATAGTACTTTATTTACTGAATCAATTATTACCATTGCCATGATGAGTGTGGATAGTTACCTGGCTATCGTTAAACCATTTCGCTATCATAAATGGATAACACATAATCGAACATTAGCTATGCTAATTTATTCATGGTTACAATCGATTGTTGTCTCCATTTTACCCTTAATAGGTTGGAGTCGCTATACTTATCATCCCTCTGAAGCAATGTGCTTTGTTGATATTTTTGTTGGTACAACATTTACTTTTTTCCTCTTTGGTATTTCATTTATTTGTCCAGTAATTATTATTGTCATACTTTATATTCTTGTCTTACGGGTTGCTGTCAAGAAAGTCGGCGATATGCCATTAATGACTATCAATCAACCACCACCAGAAGAAAAAGGTCGACGATTTAGTCTCATGTTTCGCCAACGACAGCAAATACGACGGAGTAAAAGTAAACTAAAAGCTGTAGCCAATATCTTTATTATATTAGGGCCATTTTTAGCCTGTTGGCTCCCTTATATTATTATTAATATTTATCGTTTAGTAAGTAATGATAAAAAATTTTCACCTATAGCATTATCGATCATATCCACAATGACCATTTTAAATTCAGCAATCAATCCTATTCTCTATCCGCGCTATAGTAAAAATTATCGTATTGGTTTAGGACGATTAATTTCACTTTATTGCCACCCTAAATGTTTACCGTGTTATGTTCGAGAATATGCCTTAGAAAGAAGTCTATCCTTTGTTAGTGAATATCATCATCGGCGATTATCCATGAGGCAATTCGATGAACAAAATCTACTTAAAAATGTGGACAATGTGCGCGCTGCTGCTATACGCCATATCGCTGTTAATCCTTTACTAGGGCGTTCTGGATCAATCCATCCTGTTGCAGATGATGATCCAACCCTGAGCACAACATGTAGTTCAAATACAGTGAACGGAATGTCAAAGAGTCCTACAACTGCCTCGGTTAAATCCCAATCATAG

>TriAdh142_evg1094259 type=CDS; aalen=343,48%,complete-utrpoor; clen=2124; strand=-; offs=1936-905; evgclass=maina2,okay,match:evg58816,pct:99/100/.

ccaccATGAAAATAGAATCAATCGGTCATAATCCGAATATTACTATCTATAACGGCTCCAATCACATACCAGATGAAAATTATGAAGAAATAACACCTCTTGGCATTACAAGTCTGGCAATCCAAATAAGTATTGTTGTTCTTAGTGTCATCTGCAATTCCTTAGTTATCGCTACGATTGTTGGATATCGTCGGCTGCATAATAATACTAACTACTTGGTAGTCAATCAGTGTGTTGTTGATTTCCTCTTTGCAACTCTCATCATTGCAGTTTACCCTATCCAAGCCTTGCTTGGCTATTGGCCGTTTATTAAAGAATGGTGTGCTATCCATTACTGCTTAAGCATTGGCTTAGTGATGGTATCTATTCTTAATTTATGCGCCATTAGTTTTGATCGCTATATAGCTATTTGTCATCCCTATCATCATCCTAGACTCATGACCAAGAAATCGCTGATCATTATTTTACTCTTTGTGTGGATACAGCCATTACTCATCAGTTTACTACCCATTATGTTGTGGCGAACAGATCATGATCTTCCTTTTGGTATATGCTCCTACGATATTAATCGATCATTAATCCAAGAAAGAGTTTATTTAGCATCATTTTTGATGATTAACTATGTGATACCAGCATTTATTATGCTATTATTATATTCATTAATATTTCGCACTGCATGTCGACAAGTCCATCAAATTAATAAGCAAAATATGATTTCTTCGTCTTCACTGTCAGATTCTATTCGTACTCATAAAACCTCCGATAAGGACATTAAGTTAGTCCTTTTATTTCTGGCTATCGTCGGCACCCTATATATTTGTTGGACACCTCTATATACTCTAATGATGATAAGTTACTTTGATGTATTTTATCAGCCTCACATCGGGACTTTTGTGGCAATCATGTTGGCCTTTTCTAATTGTGCTATTAATCCCATAATATACAGCTGCTCCAATGCTCCAATTCGCAAAGGCTTATTCAAATTAGTTGGAATAAAAGTAAATATTCGATCGGAACGATACGTTACCACAGTTTAG

>Tadh143_evg1115093 type=CDS; aalen=362,59%,complete-utrpoor; clen=1836; strand=-; offs=1142-54; evgclass=maina2,okay,match:evg766275,pct:99/100/.

ccaccATGGATGTTAATTCAACTACAGATTATAATATTTCTACGCCAGTAGCTGTTATATTCATAATCTATATGTCAATAGCAACCGTTATAGGGGTTATTGGTAATATCATGGCTTTGATGGTTATCTATCATCAATCGTCATTGCATAATACTGCAGGTATCTTAATGGCCAATCTTGCTGTTGCTGATCTCTTAATTACTATCTTCAACATGCCCGCTACAATGGTCAGCTTGATTAGTAGCCGTTGGCCACTCGGTTCAATTAATTGCCAGTTCAGTGGTTTTATGATGACTTTCGCGACTGGCGTATCTTTATTCACATTAGCTGTTATTAGTATTGATCGTTGTAAATCTATTATCAAGCCATTGCGCTATCAGCAACACTACCCATTTCGACAATATATTGCTTTAATATCTTTGTTATGGTTATTTCCATGCTTGATGGCAACGATGCCCTTGCTTGGTTTACAGTCTATTTCACTAGGAAAATATCAATACCTTGATTCAAAGGCTTATTGCTGGATTGCTGTCGATCAACATCGCCCTAACCTTGGCTATGTAAGTTTAGTCATGTTGAGCATGACAGCAGCATTAGCTATCTTTACCACTTGTTATTTTATGATTTATCGCTCAGCCAAGAAGAGTTATGCTCTAATAGCTTCCCTAACACAACTGTGTGAAATTAATCGTCAAAAATTTAAATCTAATTTAAAAACAGCTTCAACATTATCTGTTATCGTTGGTGTATTTATGTTTTGTTGGTCACCGTTAGTCCTCATTGATATCGTTTCCATTTTTGCTCAAGAGCCATTTTATAATGTTGTATCGATAGCTGCACTTCGCTTGGCTTTAAGCAATTCTGCATACAACCCTATCATTTACGGCATATTTAATCAATCATTTCGTAAAGGTTTTAAGAATATTATGATCCATTGCTGTCTCTTGATGAAAAGAAAGCATAGCAACGAAGTAAAGCAATTTTATGCCGAAGCCCGAGGAGCCGATATCAAGAGTGAAATCCGTCGAAAGACTACAATACAGTATAGTATTCACATTGATAATAACATTTGTAACAACCTAAGTGTAAAATAA

>Tadh144_evg111895 type=CDS; aalen=372,21%,complete-utrbad; clen=5257; strand=-; offs=4525-3407; evgclass=maina2,okay,match:evg590195,pct:99/100/.

ccaccATGGACAATCTAAGCCTAGTAGTCAGCAATGATAGTGTGAATTCCACTTCTATTTTACCTACAACCCCTGGCAATGTCAAAATTTATACCATATATGAGCAATTAATTTTACTGTTGTACTTCTTCACTGTTATGGTAGTTTCCGTTTTCGGTAATTTATTTGTTGTCATCGTCATTATTAAGCATCATCAGTTACGAACTATTACTAATTTTTTCATCATCAACGTGGCCGCAGCCGATATTCTTGTCGGAGCATTAGTAATGCCATTTACCATTTATGCTGTTATTACAAGTGTTCATCGCTTGACGGATTCTCGCTGGTGTATCGTTGAATGGTTTCTATCAGCCTTTCTACCTTCAGCATCCAGTCTATCACTTTGCTGCGTGAGTTTAGATCGATTTTTCGCCATCACAAATCCGTACTTCTATCGCCAAATAGTGACTAAACAACGAGCAGTTATCGGAGAAATTCTTTCATGGACAATTGCCGCTATATGCGCTTCAACACAATTTACATGGACCACAAATCCGATATTATTTTGCGATAAACTCGCTTACGGCACTGGTTTGTTGCATGATAAAATCTACATGACTGTAGCACTATGTATTGTTTGTTTTATCCCCATTCTTTTATTTATTTACATTTACGGCACCATCTTCAGCATTGCAGCTGCTCATGCTAAGAAAATACGACGAGACAGCCGAGCAAGCTCCATCAGTAGTCGGAAATCCATGAAAAAATCTCGTTGTAATAGCGAACAACGAGAATCGAAAGCAGCTAAAGCAACTGCTATCATCGTTGGAGCATTTGTTCTATGCTGGCTGCCTTTCACGATAGTTATGCTAGTGAATCGCTGGTTAGGCAGAACTCCTTTAGTTATAAACGCTTTCACAGCAATGGTCGTAATGGGAAATTCAGCCATCAATCCAATATTATATGTTGTCTTTAAGCGTGACATTAAATGTGTGGTCAAGAAAATTTGCTGCTGCAAACGCGTCCAACAGTACACTAGGCAAAACAACAGCTTTAGCACAACACACATTAAATTAGGACCTAATAATTCAAGTCTTGTCGAAACCAACTCAAATGCTGCTGAAACTGAAAGCTATCGATTCTAA

>Tadh145_evg1182385 type=CDS; aalen=348,50%,complete-utrpoor; clen=2085; strand=+; offs=410-1456; evgclass=maina2,okay,match:evg485056,pct:99/100/.

ccaccATGGATAATACTACATCACCTCTTAGTTCCAATCTCACTGTCACCTCTGATGTCTACATCTTACGTTATGTAACTTTACCTATAATTATGTTTTTTTCCATCTTTGGAAATTTATTAGTTTGCATGGTTACGTATTCCAAACTACAGAATGTCACAAATATCATGATTTTCAATTTAGCCCTTGCTGATCTAATCGTTGGTATAGTTATTATCCCTGCTGTACTTTCAAGTACCATCATAGGCGAATGGATTGGCGATGCTACTTCGTGTAAAGTTATTGGTATCTTAAATGCTTCCATTCTAGTCGTATCGGTTTGGATGTTAGTGGTCATTAGTATTATACGCTATGTCTTAGTTGCTTTTCCACATCGTAGCCATACCTTATTAACCAAACGAAAAGCTTACGCGGTTGTAGTCTTATTGTGGATTTGGTCATTTGTTAGTGCAGCAACAGCTGAATTATGGTCGCCCTATTACTATCATCCACAAATTTTCGATTGCTGGTTTGATTATTCTATACCATTTAAGCACCAATTTTTAATGATCTTAATTAATATTCTCTTACCTCTATCGATTATGATTTTTGCTTATATTAAAATTTACTTAATTCTACGAAAGCAATATCGCAGTTTTCAAAACCATATTCCACGAACCAGAAGCATGGTAATACGTGTTGCTATGAAAGTTGGCTCCAGTGTTGTTGTGCCCATGTTGAATAAAAAATCTCAACGAAGCCCAAGTCAAGATGCACTTGGTAATCGATTGCCTTTAGCCCGCTTGAAAGACGATGCCAAAGTAACCCGTATGATGTTTATTGTCCTGCTGGCATTTTTGTTAGCTTGGTCTCCTGGTGCTTTCATTTTACAATTTTTTTTTTTTATCCAACCGGTATCACCATTTATCCTTAATTTAGTCTTTATAGTCGGCTATAGCAATAGTGCTATTAACTGTTTCATTTATGGTTTTCTAAATAAGACATTTCGGCATAGTTTTGAAAGAATTTTGACGTGTCATAAGTGTCGAAAAAGTACTAATTCGTCGTTTTAG

>Tadh147_evg12615 type=CDS; aalen=389,46%,complete-utrpoor; clen=2496; strand=-; offs=2149-980; evgclass=maina2,okay,match:evg370408,pct:99/100/.

ccaccATGTCATGGAATTATCGCAGTAGTAATTCTACCATTGTAAAAGAAGGTCCTCAATCACGTTCGATAACACCGCAAGCGTTAACTTTCGTTTGTATAACTTTATCCATCTTTGTAGTTAGTTTTGCAGCCAATTTAATTGTTGCTGGTACTATATTGTGTCGGAAGCATCTGCGAAATATCACTCATCTCTTTGTCTTCAATCAATGTATCGCTGATCTCATTCGTGTATCATTGCAAATACCTCCATTAGTGGTTAATATGTTACTAGGCTATTGGCCATTTGGTCCTGTCTACTGTAACTTGAGTTATCCTATTACCATTTTTTTGATGTCTGTATCTATATTAAATGTTTGTAGCATTAGTATTGATCGCTATTTTTCCATCACATCACCATTATTGTATCAGGCCAGACAAATTATGAGTAAGAAAATTGTGCTTGTCCTCTTGCTTTTTGTTTGGCTCGAACCGCTTGTAATTGCTCTATTGCCTATGTTTGTCTGGCGTGATATCGGCTATGATCCTCGACCTGGTTCTTGCTTTGTTCAATGGACAGCCGCGCCTAGCTTCGAACGACCGTACCTAGTTTGCCTTATGGTAGTAAATTTCTTCGTGCCATGCCTTGCTATGAGCATGATTTATGCTCGCATTTTTGTCATCGCTTCTAAACATGCTAAACGTGTTAAATATGAATCCTCAGCGTATAGTATTGAAGCATCAACTAGAGCGCGTATTGATGCTAAAGATATTCGAATAATTGCAGTCTTGGCAACGATCGTTGGATTCTTTTTAATATGCTGGTTGCCATTTTTTATCAATGGGCTTGTATTATTATTTAGCCCTAGACTGATGTCTACTGTATTATATACGGCATCAGTTTACCTCTCTTTCGCCAATGCTGCTATCAATCCATTAATTTATGCAATTTTTACGCGTGATATACGACATCAGATATGTTTTTTATTCTGTCGCCCATTAGCGTCACGTTTTCATTTAATTTATGATAACTATGGTAGGAACCCTACGCGTAGTGGGTTAACATCGCAAACTTATTCATCAACTCCACATTGCAGAAGTATGGTTACAAGCGATTTCCTTATACAAGGATCAAGTTCATTAAACAATGCGCGTGTAATTATGGCCAAAGGCATGCAAGAAGATAGTACAATATAA

>Tadh148_evg1429139 type=CDS; aalen=426,56%,complete-utrpoor; clen=2271; strand=-; offs=1355-75; evgclass=maina2,okay,match:evg1738611,pct:99/100/.

ccaccATGCGGGCCAAGGAAAGCAGTCGATTTTTATTGATTATCTATATTCATATATTATTCTTCTCAATTAAGTCTATGGCGATAGAATTTGGAGATATAAGAGTTGAAGCAAAAGGTTTTCCCCCACTTAGTGGCAACCATGTGGCTGGAAATCAAGTGTCGAATATAAATAGTCGCTATGGTCAAGAAATAGACTTTAAGACGGTTTATATCTTTACAGTATTCATCACCGGAATTATCTGCAATGCCATCGTTATTGCTACTATTGCTTATAATCGCCAACTTCATTCTGTAACTAATTATTTTATTGTCAGTCAATGTTGCGCTGATTTCTTATTTGCGGCTTTAGTTTTACCTGCATACTGTCTGCAAGGTTTATTAGGCTATTGGCCATTTATGCCAGAGTGGTGCGCTATCCATGCTTGCTTATCAATTAGTTTAGTAATGACATCATTACTCAATTTATTTGCAATAAGTCTGGATCGATATTTGGCCATATGTCGGCCTTATCAATATCACACTTGGATGACTACACGCTTCGTGATTGGAATGCTACTATATATTTGGTTTCAGCCGCTTCTACTAGCCTTGATTCCAGTATTTGGTTGGAGAGGTGTTCAATCAACTATTGGCAGTCATTGCGTTGTCGATTTATTTCTATACATAAAACAAGAGAGAATTTATTTGTTAATTATTACATTAATTAACTTTTATATCCCAGCAACAATAATGTTTACTTTATATATCCTGATATTTAAAGTTGCTATCCGCCATATTCATAATATTGAACAACAAAGGAATGTTCAAAGAAAGCAACATTATTTTGGATCTATTCAAGATCAACTTAATCATCGTTATTTGCGCTTAGCTAAAAACGATTTTCGATTAATACTTGTCTTTGTAGTTGTTAATGGCGCATTTTATTTATGTTGGACGCCATTTATGACATTTGGCTTAATAAGCTTATTCCATTTAGGACCTGTAACTGATAACTTAGGAATTGCTTGTAGTCTAATGGCGTTTACAAATTGTACACTCAATCCGATCATACATAACTTTTTTAATCGAGAAATTCGAAGCGGATTTTTCAGGATGGTTTGCTTCCATACCAGACGCAAGCCTATCAATAGGAGCAACACGCATATGTCAGGCACTCAATGTTCGTCTACTCATGGGAGTACATTTGTTATACGTCACGATCTTATTTCTGCTAAGGATAAAGCGATTATCGAGCAACTGGTTAACGTGAGCAGGATGAAAGCAAAGAAAGTGACTTATATTTGA

>Tadh149_evg1609341 type=CDS; aalen=429,53%,complete-utrpoor; clen=2394; strand=-; offs=1729-440; evgclass=maina2,okay,match:evg914988,pct:99/100/.

ccaccATGATTAAGATGGACTTATGGAATTTGACTTATCAAATAAAAAATGTAACATCATCGGGACATGATGGTTATACAGCCAAGCCTAACGACACTGATGAAGCTCTACCTTTAAATCCAAGTGAATGGATAGTTATTGGATTCAGCTTAGCTGTAGTAATTATCAGCGTGGTTTGCAATGTAATAGTGATTGCTACTATATTATGTAACCGTTTTTTACGCACAACAACCAATTATTTCATGATCAGTCAAAGTATTGCCGATCTTTTGTTTGCCTTAATAGTGATTTTAGGCATGGTGATACAAGCATTGCTTGGCTATTGGCCATTTGCGCCATTTTGGTGCGCCATTTATAATGCATGTGGTATTGGTTTAGTCATGGTTTCCATTTTAAACTTCTGTGCAATTAGTATTGATCGCTATATTGCAATTTGTTATCCATTTCTTCAGCACAGGCTTAACTTTACTTTATTATTTATACTTGCCAGTGCTTATATCTGGATGCAACCGATTATCCTATCACTATTACCCATGGTAATATGGCATGATTATACTCAAGTGGATATCAATGTGTGCGGTCATGTTATTACCCATACACGAGAGAAAATCTATTTTCTGGTCTTAGTTATTGTCAATATCTTTCTACCAGCATTGCTGCTATCAGTTCTTTATTCTATAGTATTTAAAACGGCTTGTGGCCATATACAAAAAATTAAACAACAAAATAGCCTTCACCGTCAATTAAGTCGAGCTGCTGATATTAAAAGACCGATTACATCAAAAGATTTTAAATTACTTCTTATTTTCGTATTCATTATTGGTACATTTTATCTTACTTGGACTCCATTTTTCGTTGTCCTCATTATAAACTATTTTGATGACGATCCCTACCATCCTAGTAGGGAAGGTTTCAGATTTTTTAGTTCCATAGCTATTCTAATTAGTTTCACTCATTGCGCTATTAATCCAATCGCTTATAATTTTATGAACTCCGAAATGAGGCGAGGTTTATGGAATTTACTAGGAATTAGTAGGAGACGGCGTCGTGCATCGTTGCGTTTAAGTTTGCAAGCAACTTATACTAAGACATCGAGGGCATCTAGATATTCAAAACGTAGCAACGGTGGTCCGCGTACTAGTAGCAATTGTAATGGCCAGCAATTAGTTGCTAAAAATCAAGATAATAAATTGACTCCAAAGGCTGGTTTAGAAGATAGACTATATCCACTTATATCGCCATTACAATCTTATCCTCCAAATAAAGAACCTGCAATGGCTTTAACTACAGTATGA

>Tadh150_evg19734 type=CDS; aalen=355,45%,complete-utrpoor; clen=2323; strand=+; offs=484-1551; evgclass=main,okay,match:evg900155,pct:99/100/.

ccaccATGGCTTGTACCAATATTACCGCTATCCTAAATAGGACATTAACACCAACTGGCATAAATAAGCTGGCAGAGAATGCAAATATTCCACAAGAGCAAGATTCATCTTTAAATTTGGGTGAAAAATTTAAGTTAGTCACTATTATTTTAATCATGGCTGTAGCAGTCATTGCTAACGGAATTGTTATAACAGCAATTTTACTAAGCAAGAAATTGCGTTCAAAACATAAATTTCTATTGAGCGAATGCTGTGCCAATTTCTTATTTGCACTATTGATCATGCCGCTATCTCCTATTCAAGGCTTGTTAGGCTGGTGGCCTTTTGGCCACATTTATTGCAACATTAACTTTTCTGTATCGATTGCTTTAGTTTTTACCTCTATTCTAAATATTCTAGCTTTAACTATTGAACGTTACATTTCCATTTGTCATCCTTTTAAGCGTCATTTTATATTAAAGTCAAGCCATGTTGTGGTTATGTTGCTTTACGTTTGGATCCAACCTCTAGTTTTATCTTTGATTATATTTATATGGCGTGATATTAGCTATCAATATCGGCCTAAGCAATGTATGCCAGAATCACCTCAAAAGTATCAGACTTATGAGAAACCTTATTTTACTTTTTTAATTACTACTAATTTCTTTATACCGGTGCTGCTTATTGCTGTTATGTATGTTTGCATTATACGTATTGCATGGAGGCAAAATCAACACATTCGATCTCAACATAAACAACTTAATCATAGTCTCAATAAGAAAATGAACATTACAATTAAGACAACTGATTTGCGGGCTACTTTTGTGTCTGTTATCATATTAGGTATATTTTGCGTGTGTTGGATGCCATTTTTTATTATTACTCTAACGGTGGCATATGATATTCGTTACAATCCTGGACAATTTCTATTTATTAGCATTATATTGGCTTATGTTAACTTTGCTATTAATCCAATAATCTATACAATGTTAACTTCGGATATAAAATCAGCGGTGAAAAGTTTGTTCCCATGCCAAGAAAGTTATGCATTCCGTAACAGAAGCCGTAGTTTGAGTAAGACGACTAACGTTTAA

>Tadh152_1606805 854 1606805_m.854 type:complete len:372 (+) 1606805:598-1713(+)

ccaccATGTCTAATCATAGCCTTAGCTCAAATCAGACTAGCGGATTTACACCGCTTGCTAATGCTGAAGCTCAAGTACATTTTACTGCTTATGTGATTGTGATGGTTATAGCTATCCCTGCTAATATAGTCACTATTTTTTCTTTTATCATCGATCGTACATTGTGTACCAATTTTAATCTCTTCTTGCTTAACATGGCTTTCATCGATTTGTTGACCAGTTTATTTCGCATCGGTTTCAACGCTTTCTGTATTGCCACTATTGGAAATTCTACATGGCCATATGGTCAGCCACTATGTGATTTTGACGGATTTATACAATGCGTTGTCTATGCCGCTAACATTTACACTCTCATGGTTATAGCAGTTTGCCGTTATATAGCAATTGTTCATTCCAAACAGCATTTAATTACCAAAAAAGTTGTCTGTATTATTATTGCTATCATTTGGGCCTTTAGTTCCCTGTGTTCGGTCTTTCCTATCATGGGTTGGAATCGATTTCGCTATCAACCTTACGAGCACGCTTGCGTGCCTGATTGGTCTTATGAGAAGAGCTTTCCTTTTTATGTTATGGCTACCGAATTTGTTCTCCCAATGATAATTTTACTTTACTGTTACACCGGAATATTTATCAACCTAAGGAGAAATTCATTTCGAATGCGTACTGTTTCAGAATTGAACAGTGGCAATAATGAGGCCATGAAGAAAATAGCCCAGCGTGAGGGAAAGGTTTCTCGTGGCATGTTTATCATCTTCTTAGCCTTCGTCATTTGTTTTGCACCATATAGCATATGTATGTTTCTATTAAAACCTGTATTCAATATTGAATTAAACAGGGGTTTGGTCTTCTTTTGCGGATTTATGGCCAATGTGAATAGCTTACTGAATCCTATTATTTATGCTATCTTGTATAAACGCTTTCGGCAAGCTTACTATCAAGTGCTGTGCTATTGCTACAGTAAATTAACATTCAGCTCTTTAGTGGATCATGCAGGATGCAATCGCAATCACGGATTGCGATCATCAGAAATGGGTGTCATCGAAAATAATAACGCCAATCATTCTACCAGTCCATCTATGAAAGCTTCTAATGATGATGGTAAAAAATATCGAGTGTTTTAG

>Tadh153_evg1004178 type=CDS; aalen=344,36%,complete-utrbad; clen=2838; strand=-; offs=2259-1225; evgclass=maina2,okay,match:evg1220421,pct:99/100/.

ccaccATGACCGAACTCCTCAATAATGGCACTTGCCCATCCAATTTTACTACGATGAATATGGACACTGTTTCAATTAGGCTAGTTTGCTTATCTATAATTGGATTTCTTGGTATAACTGGCAATATCATGGTAATCTTGGTTACTTATACAAAAATTGGTCGTAAATCACCGACAGATTATTTAATAATTAATAACGCTGTTATTGATGCTATCCAGAGCGGGATTAGATTACCGCTGTTGTTGGTCAATGTTGCAGCCAATTGTAATGCATTAGGAGATTTTCTATGTCATTTTCATGGCTTTCTAAGCGGTGTATGTTTTATTGGTTCCGCTTATTCCTTAGCTGCATTGGCTGTATTTCGTTATTTTAGCGTTTGTAAATCTCCAGCTATTGTACTACATACAAGGCATGCTGTATATGCGATTGGTATAGTTTGGATGATTACGTTGGTTATTTGTCTGTTACCATTTTTGGGCCTAAGCGTTTACGAATATAGTTTAATGGAGAGATCTTGCCTTCCAAGTTGGGGTAGTAGTGTGGCTAATCGTATTAACGCTATTATTGAGTTGATTGTCGATTTTATTGTGCCCTTAGTTATTACATTAACATCATACCTTCAAGTATTTTTAACGTTAAGACGTATCGATCATAACCTGGTACGCCATGTTAGTAACGGAGGATTGCAGCGGCGTAATACAACATCATCAAGGAAACCGATTCATTTAACTATAGCACTATGGCTATTGGTGCTTAATTTCTTGCTGTGTTACATAACGTGGAGTATTCTCACTTTTATTATTATTCCATTTTCAAATTATCAAATCTCTATTTCGGATGATTTATATTTCTTTTCTGTCACCATGTTATACGTAAATAATGCTATCAATCCAGCTATTTATCTTGTCATGTTGAAATCTTTTCGTCGACGCTATATCATATTATTAAAAAAGATTTTCTGCTGGTGGAATAAACGTTCTTCCGTTGTTCGAGTGAAACCGCTTAAATCAGATAATACCAGAGAAGAAAATAGTAAGTGA

>Tadh154_evg1096055 type=CDS; aalen=416,53%,complete-utrpoor; clen=2359; strand=-; offs=2266-1016; evgclass=maina2,okay,match:evg768579,pct:99/100/.

ccaccATGTCAGTATTATCCTGCGGATTCTTATTAAACTCGTTCAAATTAATATTTGAAAGAATGAGTGTTCTAGTGTCATTACGTGGATCTAGCCATGATCATCATGGCAATGCAACCATTCTGAACTTCTCACCTGCAATTTTACTCGCTCGTGGCTTGGCATATATAATATTAGCAATTGCTACGGTTATTAGTAACTGCATTGTCCTCTATGTTTTTCATAAGCATCGTTCCCTTTTACGCATTCCTAGCAATCTTTTTATCCTTAATTTGGCTACTATCGATTTGTTAAATGGATTATTTAGGGATACTTTTAATGGCGTAAGCTGGCTGAATAATGGCTGGACATTTGGTGAGCCGTTATGCCGCTTTAATGGATTTATGCAAACTATTTGCTATGTAGTAACCATATTTACTTTGTTGGCTACTGGAATTTGTCGTTACTTGGTTATCGTCCATTCGTATGGAAAGGTGATTACGAGGAAAGTAGTTTTAATTATTATTGGTTGCATATGGATTTATAGTATTTTGGATTCTCTAATGCCAATTTTTGGTTGGAATCGTTACATTTTTCAGCCACTGGAATTTTCTTGCTTACCTGATTGGTTATCCGTTAGCGGTGCAAGTTATGTCATGTTCGCTTTAATTGCCGACACAATTATTCCGTCGATATTAATTGGATTATGTTACATCGTCATCTATATCGTTATTGCTCGAAATGCTAAAAGAATGGCTAATCGATTCCATATCGAGCATTCCAACAACGTTATTCCTTCAATAGGCAATTCAGTCGATAACTCAGTAGCAAACACCCCTGAAAATGATCGAAATCGACAAGCACGAGCAAGAAGGTCAAATGGGAATACTTTAGGAAGAAAAGAAGTTAAAATTACAAGAGCCATGTTTTCAGTCTATGTTATCTTCCTTGCATGTTATTTGCCTTACGCTCTTACTATATTTATTATCGCACCAGCTGGTGGCAACGTTTCACATAATATTATTTTTATGGTTGGGTACCTTGTTAATGTAAACAGTGCGATCAATCCATTTTTCTATGGAGTCATTTATCGCCAATTTAGGAAAGCGTATATCGAACTGTTCTGTTGTTGTGGCCTTAGTGCAGTGTCTGTAAAGTTCCCCACTAGTCGAAGTCGGCATCGACATCTTCTTAATTCTACGCCAATAATAGATCGAAAAAATCGAGTCGATGTTCTTCGTGATGGAATGACCGATGCTAGCACCAGTAATCTTTAA

>Tadh155_evg1224494 type=CDS; aalen=346,66%,complete; clen=1572; strand=-; offs=1458-418; evgclass=maina2,okay,match:evg369914,pct:99/100/.

ccaccATGAGCGCTGGTGCTATCCATGCTAATCATTGCTTCAACAATTTTACAATCCCTCTATCTCCGTCGTATAGAATATGGCCAAGCATTGCTATCGGCATCATTTTTACTATAGCATTAACTGGCAATTTAGCCATCATCTTTTTTACAATTTCCATTAAAAAATTACATACCTCAACTAATATTTTAATCGCCAATGTTGCTTTAAATGATCTCTTTATTACAATTATTGTCATGCCAATATGTTTCACCATTTCTTTAACTTTGGTATGGCCATTTAGTTGTACATTTTGTATTATAGGTGGATTTTTAGACATGATACTTTACAATAACGCCCTATGCTCTTTAGCAATTTTAAGTATTAATCGTTATTATGCCATTGTCTGTGCCATGGATCCCAATTTTGTTGCCAAGATGTCTATCAAAAGAGCTAAAATCATGGCTGTCGGTGTTTGGATGTATTCAGCAATATGGGCTGCGCCACCATTATTTGGATGGAATCGTTATGTCTTTTACTTTTCCAAGTTAATGTGTAATATGCAGCAACGTGATCCAAGTCTCTATGTGACCGTAGTAACTCTATTTTGTACTGTTCTTCCAACCTTTACTATCTTATATTGCTCAGGTAGAGTAGTTATTACGCTATGGAAGAATGCTAAAAATTCAAGTATAGCAACATCCAATCGACAACGAATTGAACGCCGAGTAACATCCATGATTGGCCTAATTTTATTAGCTTATGTATTATGCATTGCTCCCAATTTTGTAGCTGGTTCACTCTATGCTTTCGATTCAAGTAATAGTGAAGATTTGTCTAAATACAACGATTTTCAAACCATCGCTACTATCGTCTATTTTATGAACAGCGCCGTCAATCCTATTATCAACGGCGCAATGAATCCATATTTACGTCGAGCTGTTAAACAACAATATCACTGGCCACGTTGTCTAGCAGCATGGAGGCATCACTCAGTCAATCCTATTACTGCTACGAAGAAGGTGAGAAAAAGTTTTAGCCAATATGTCTTTAATACAAGCGAATAA

>Tadh156_evg12947 type=CDS; aalen=465,65%,complete; clen=2140; strand=+; offs=321-1718; evgclass=main,okay,match:evg1688447,pct:99/100/.

ccaccATGAGAAATCTTCATGGCCATGATGATGACGATAGCTATGCAGAGGGTGATTATTTTAACTGGACAACCAACCATACTTTACCTGAAACTTACGGACTACAAGAATATTTACAGACCATTAGCATAGGAATACTCTTTTTTATGTCATTGATAGGCAATTCATTGGTTTTACGATACATTGTCAAGAAAAGGAAACTTCATACAGCAACATTTATTCTAATAGCAAACAACTGTGCAGTGGACTTGGCAATAACAATTTTCATTATGCCTATGTGCTTTATGGTTTCCATTTTACGTGATTGGCCTTTCAATTATGCAGCATGCCAATTTTTTGGATTTATAGATATGACACTTTACACCGGTTCCTTAATGGCATTATCTGCAATTAGTGTTATACGCTATTTTGCAATAGTATGTATTCGAGATAAAAATTACCATCTGAAAACCTCTAAGAAAACGGCCATAACTATGATATTTATATGCTGGTTCTATTCACTTGCATGGGGTAGTTTTCCATTAATGGGTTGGAATCATTACATCTTCTTTATTGAGAAATTAATGTGTAATATGAATCATCGAGATCTAAATCATTATACGACCGCTAGTACGATATGCACAACATTAGTACCTTGCTTAGTGGTTATCTATTGCTCATTCAGAATTATTCTAGCCATACGCCAACATAGCCATCATAATCATTTTCATCAAACGCAGCAGCCTCTACAACGTTCTCGTGTTGATCGCAAGGTAACTACGACTTTAATTATTGTCTTTATAGCTTACTGCTCTTGCACGATACCGCAGGTAGTTGTTGCTTCAATTTATAAATGGGATGAAGATGATACCATGAATCAACGTATATCTTATAGAAACATCAAAATCATGGCTACTATTCTATATTTAAGTAATTGCACCGTTAATCCAATCATCTATGGCGTATGCAATAAAAATTTAAGGTCAGCCTTTCATTTTCGTGGTTATTTTGATCGTTTCTTTGACGTTGATGGCGTTTCACAAGGCCCAACAGCCATGGGTCCGACTTCTATTCAAGTCAATGTTAATGAAAGTCCTACTACAAGATCTAGAAAATCACCCATTTCCCTCTCGAAAGTAAAATGCTTTGAAACTAACTCATCTAAGAATTGGCAAGCTAAAACATTGCAAAATCAATCTGGACTTAGACCAATGTCAGCTAAGTTGGTCAATAAAACATCGCCAATGGTGGCAAGAGCAAGATTATCCGATGTAAGTTTAGTTTCTATTGAATCGACCCAAACAGCCATAGAGCCAATATCGCCGTTAAAATCGAATCCCAACGTTAGGAATTGGAAGAATAATACGGGCATTGCAGTAACACATCCAAGGATTAATCATTTAAAAAATCACGATATACACTAA

>Tadh157_evg12949 type=CDS; aalen=340,71%,complete; clen=1428; strand=-; offs=1167-145; evgclass=main,okay,match:evg99149,pct:99/100/.

ccaccATGGATCTTAACGCATCGACCAACTCTACATCCAATGATAAATGGCCACAAGGCACCGAGGCAATTATCATCTCTGTGATATTAATAGTTATCTTTACTATCGCATTAATCGGAAATTTTCCTGTCGTTTTCATGACTTATTGGGAAAAGAAGTTGCATACATCAACTAATATCTTAATAGCAAATCTAGCATTTTGTGATTTACTAATCACAATTTTTATTATACCAATATGTTTTATTGTTGTTCAAGTTGGTATGTGGCCCTTTAGTAGCATCACTTGTCAGATTTTGGGCTTTTTAGAAATGGTATCCTATACAGCATCACTATCATCATTGGCGATGATTAGTATTAACCGGTATTATGCCATCGTTAGAGCCTTGGATCAAGATTTTGCCAAGAAAATGTCTGTCAAAAGAGCTAAATATATGATTGCAATAGTTTGGATTTATTCACTAGTTTGGGCAAGTCCACCGTTATATGGATGGAATCGATATTCATTTGTATTAGGTAAACTTATGTGCAATATGAATCACCGTGAAGGCCAATATCCGCTACTAGCGACAATATTTTGCACTCTTCTACCGTCAACCATTTTGTTTTATTGCACGATAAGAATAATTGTGTATATGTGGCAAAATTTTACCAACCGAAGCATTAATTTAACCCGAAGCAATCGATTAGAGCGACGAGTGACTATTATGATGGGCATAATCGCTATCGTCTATTTCTTATGTATAGGTCCGAACTTATTAATGGCAACTGTAGTCCGATTTAATGAATATTCTGGAAGTTTACATTTATACTACAGAATACAGGCAATCGCAACCATACTCTATTTCAGTAATAGCGCTTGGAATTCCATTATATATGGAGCAATGAATTCCTACTTACGACGTATCGTTCAACAAAAATTATTGACAAATATCTGTTGTAGATGTCGAAGTGACCGTCATTCAAGTACAGTCATTCCTTTGTCGGTTCGATTTCGTAGTAAACGTGCTAATTCACATTTAATGCTATGA

>Tadh158_evg1356946 type=CDS; aalen=359,58%,complete-utrpoor; clen=1855; strand=-; offs=1741-662; evgclass=maina2,okay,match:evg1265808,pct:99/100/.

ccaccATGGTTAGTAACAATTCCAGCTTGTATCCTGAGTATGGGCCGACATCCTTAGCAATTATTCGTGTTGCTGCTTACTCCGTGATAGGTGTAATATCTATTCTTGGCAATTTGCTGGTTTTGATAGCTTTCTATAGAGAAAAATCGTTGAGAATTCCTACAAACTACTTAATCATTAATTTAGCCATTACTGATATTTTAAATGGAATTTTTAAAGATTCTTTTATTATATTGGGCACGCTAACAAATCTTCATATACGTTATAAGGCATTCTGTGACTTTGATGGATTTATGCAAATATTACTTTACTTATTGACAATTTATTCCTTATTAGCTACAGCTATATTCCGTTACTTAGTCATCATTCATTCTTATGGTAAAAAAATTACTGTTCGAGTTGTGAAGATAACTGTTGCTATCATTTGGACTTACAGCCTCGCGATTAGTCTTTCGCCAGTACTCGGATGGAATCGATATGTTTACCAACGAATGGAATCTACATGCTTACCAGATTGGCTGTATCCAGCTGGCAGAAGCTACGTTATGTATGCATTTATCACTGATTCCATTATTCCTTTAAGTCTACTAGGAATCTGTTATTTGTGTATATATATATTTATCAGACGAAATGCTAAGAGATTCATGGTTAATATTAAGAAAATCCAGCCAGATAGCCTGAATAGAAGTTTACTTCGCAAATCCATTAAAAAAGAAATCGAAATTACCAAAAAGATGTTTTTCGTATATTTAGTATTTGTTATTTGCTACTTTCCCTATACCATAGTTGTATTTATCTTAGCCCCAAGTGGTGTAGCAGTCCCTCCAGTTGCCTTTTTCATTGTTGGTTTTTTAGTAAACGCGAATAGTGCAGTTAATCCCTTCGTGTATGCTGTAATCTATAAAAGTATTAGAAAAGCCTACTATGAAATTATATTCAGGTATATCCCTAGTTGCTGCTCTTTACCAGCTTCCACGTTGGATTTTAATCAAACAAAATTATCTACTGCGATCGACAAATGTAGTACTGATATTATATATGACACTGGCGATCAGAGAGAAAATTTAGTGTTAGTAGATATTTAA

>Tadh159_evg1417095 type=CDS; aalen=346,70%,complete; clen=1474; strand=+; offs=368-1408; evgclass=maina2,okay,match:evg1417096,pct:99/100/.

ccaccATGTGGCAAAATTTAAGTACGAATCTTATCGCAGATTATCACCCTCTTCAAAACGTATGGATTGCAAATGTCTCTTTAACAGCTCGAGATCGCTATATTCGCATCGGCCTTTATAGTGTGCTATCGTTAATCACAATTTTGGGAAATATGTTAGTCTTTTTAACTTTCTATAAACATGCATCATTACGTACAACTTCCAATCTATTTATCATCAATCTAGCTATAACTGATTTATTAACTGGAGGAATTAAAGATACATTATTTATTTATGGTCTAACTTCGTATAATTGGCCGAAAAGTGCAATTCTATGCTCATTTGTGGGCTTTATTAATTGTGTTTGTTACGTTGCGACTGTATACACGTTAACAGTCATAGCTGTATTTCGCTATCTTGCAATTGTCTGCAATTTGGGCAGAAAAATTAAGCGTAAGCATTCCATCTTGACTATCGCTTGCGTTTGGATCTACAGCTCTGCATGCTGTTTAATGCCTATTATTGGATGGAGCCGCTATATCTATGAACCTACGGAATGTACTTGTGTACGTAGTTTAGATAAGAAATATTATTCCTATACGATATACATTCTAGTAGCAGATTTTCTATTACCATTATCCGTTGTGTCGTTTTGCTACGGAAATATTTTTGCAAAGATGAAAAGAAATCATCACCACATTAAGCGTGATCTAAGTGATGGACAGAAACTTGACGTTGCGGAAAAGTTATCGCGGAGGGAAGAGATTGTCGCGCGGCGTATGTTTATTATCTTGGCTGAGCATGTCGTCTGTTTTTTACCTTATACAATTATTGTTACGATGTTGGCAGCGAATGGCGTTGATATTGAGCCAGTTTGGTATTTTATTGTTGGTTACTTACTTAATTTAAACAGTGCTCTCAATCCTGTTACGTATATTGTTGTCAATCCTAGGCTTAAGAAGGGCTATATATCAGTCATTTGTTGCAAAGATAACTTTACGCGAGTTAAGCCAACAAAGGCTGGAGGTAATCAGGGTTTTCTAGAACTTCGTAATGTAGCAAAGTAA

>Tadh160_evg1429675 type=CDS; aalen=317,61%,complete; clen=1562; strand=-; offs=1377-424; evgclass=maina2,okay,match:evg1001127,pct:99/100/.

ccaccATGTATCATGGTCAAAGTAATTATAGCGACTATATCAACGGAAATTACAGTGGCGTTCCAATATCAGAAGTAGAGCAGACATTACGTGCTGCAGCATATACTTTTACTGCACTAACAGGCTTAATTGGTAATACAATCGTCTTGATCATATTTTATCAAAGCCCTAGCCTTCGTGTACCAGTCAACTATCACTTAATAAATATTGCCATTGTTGATATTATTACATGCTTATCTCGTCAAATATTTAATATTATCGGTGTAGTTACGAATGGTTGGCCATATGGTAAAGACTGGTGTGATATTAATGGTTTTATGCAAACATTATGTTATTTATTAACGGTTATGTCACTCTTGGCCATTGCAGTTTCTCGTTACATGGTTATTGTGCATAGACGCTCTGTCAGTCATAAGCAAGTTTGTATCACGATAATTATCATCTGGATCTATTCACTTGTTCATTCCTTGTTGCCAATTATAAAGTGGAATCGATATTTCTATCATAAGAAAGAATTTGCTTGCGTTCCAGATTACAAGTATGAACTTACATATCCTTTATTTTGCGTTATAGCTGACTTTTTAATACCGCAAATAATTATTAACTATTGTTATCTAAAAATTTACTTGATTGCAAGAAATAATAGCTTAAAAGTATTTAATAAATCAGCACGACTGGATAAGATTCGCTACAATGTCAAGTTAACTCGAAATATGTTTTACGTCTTTCTTGCTTTTTTAATCTGTTATATACCTTATAGTATATCGATGTTATTTATTTCACCCTTTGGAATTGAATTTCCAGTCTGGTATATTTGGTTTTGCGGTTGGTTAGTTAATTTAAATAGCTCTATCAATCCGATATTATATTACTTTATTTTTCGACGCTTTCGCCAATTATATATTGCAATGTTATGCAAATGTAATTGCAAGATTGCACCCGAAGTTAGCCTGCGATAA

>Tadh161_evg1477936 type=CDS; aalen=392,41%,complete-utrpoor; clen=2828; strand=-; offs=1666-488; evgclass=maina2,okay,match:evg921283,pct:99/100/. was not possible to synthesize, complicated gene

ccaccATGTCTACCAATAGCACACCCTGCATAAATTCGACATGTGGAACACCTAGCGAGTGGATCGCGACTAAGATAGGAATTATAGTATTTATTATTATCACAGCAGTTTTAGGTAATTTGCTTGCTAGTATTGCGATATTAAATGAACGTCGCATGCGAACAGCAACAAATATTTTATTACTCAATTTAGCCTTACTAGATTTCTTAACAGCTTTAGGACGCTTGCCCTTATATTTGATTAATATTGTTGCTGATCGTTATTATTATGGTGTCGTATGGTGTAATATCCATTCTACTATTAGCGGCATCTGCTTCATTGGCTCTATTTATACATTGATGTTTATCAGTTACTTCCGTTATCGCATTATATGCCGTTCAACTACAACTCGGCCTAAACGGCACCATGCAATTTACGTCTCTATTTTTGTTTGGTTATGGAGTACTTTTGTCGCGTTATGGCCTATACTGGGCTGGAGTGCCAATGAATTTAATCCCGTAGAAATGGCATGCATTCCAACATGGAGTGTCGGAATTAGTGGATTAATCAACGGTATCCTGGAAGTTGTTATTGATTTCTTACTACCGTTTAGTATCATCATATTTTGCTACATTCGTATATATATTTATTTAAAAAAGAAACGTCCAGCATTAAAATCATCAAGATCCAAGAGACAAAGTGCGCTAGTCACTAGAACCATGGGAATCGTCATATTAGCCTTCGTATTATGTTATGCACCATGGAGTGTAATTGTATTTATTGTTATTCCATTTGGCCAAGAAAACGTTTATATCCCGACGGGTTTCTACTTTATTACAACTGTCCTTGTATTTATCAACAGCAGTATTAATCCAGTTATTTATGGATTTCAAATACCACAATTTCGGCAACAGTATCGTCGCATTATCTGCCCATGCAAAAAATCCAATATGATAACGCCAAGTGAAGGCGAACCATCATCCATCAAGCCACGTACATGCAGCCATTCTGGTCAAAAAACACAGATTCCTGAAGAATCATTTTCATTCAATCGCCGAACTGCAAGTGAATCAGGACATTACAGGAAATCCAAGAGGTCTTATGTCATGCAAAATCAAATAACAGCTCGCCCGATGGAGTTAAAACGAAGCGATGAAGCTTACTATTCTGATGCTGACAATAGTGCAGCAAAAACTTATGGTTAG

>Tadh162_evg1659740 type=CDS; aalen=365,52%,complete-utrpoor; clen=2076; strand=-; offs=1852-755; evgclass=maina2,okay,match:evg1770560,pct:99/100/. was not possible to synthesize, complicated gene

ccaccATGAACTCATCCAATACTGTCAATAACTCATTTATCCCCTTTAGCGATTTTGAAGCTAATGCCCGTTTTATTGCTTACGCAGTTCTTCTTGCATTTACCGTCATTGGCAATATCATCGTCATAGAGTCATTTATTGTGGTTCGAACTCTTCGTACACCGTTTAACATTTTTCTTCTCAATTTAGCCACTGTTGATTTATTCACCGGTATATTTAGAATGGGTTTTTACCTCTTCAATATCGCTTATGCTAAAGGATTTGAGTGGCCGTTCAGTCCAGCGTTATGTAATTTCAGCGGTTGGGCATTAAGTATTGCTTTCTCAGCTAATGTTCATACCCTAGAATTAATGGCTATTTTTCGCTATATTGCGGTGGTTCATAATAACACCTACAATCTCACGCAAAAGAGAGCTGTTATATCGATCGTTTGCATATGGATTTATTCCAATATTATCGCTCTCCTTCCAATCATCGGTTGGAATCGGTATCGATATCAAGCAAACGAAATTGCTTGTTTACCTGATTGGTATTACGAAAGTAGCTACCCACTTTTTGTTACTATTTGCGATGTGATCATTCCATTAACCATTCTATTCTACTGTTATGTGGCTATTTATATCACGATGAAGAAAACGAGTCGACGTTTCCGTGATGTTTCCAATCAGGGCATGAATATTGCACTCCAGCAAATATCTAAACGAGAAGCTCGTGTTACGCGTGCCATGTTTTTGGTTTTCGTCGCTTTTCTTATTTGTATTGTGCCATATGCTGTCTGTGTATTCATCTTGTTCTCTATATTTCACATTTGGGTTGGTCGAGACATGGCATTTTTAACTGGTTATTTTGTTAATCTAAACAGTGCTATCAATCCTATATTGTATGGTTTCCTTTACTGCCGCTTTAGAAAAGCTTATTACTATACTGCTTGTTTGTGGTGGAGAAAATTATCCAGTATATACACGACTCTATCCAGTCTACACCATCGGCATAATAAAGTACGACTTGTGAAAGACTGTGATGGCGATTTCAATCGATCAAGAGGAACAGAAATTATTAGCATGAATCACCATGATCAAGCGATCATTCAAGAAGTTGACTAA

>Tadh163_evg19497 type=CDS; aalen=319,24%,complete-utrbad; clen=3980; strand=-; offs=2811-1852; evgclass=main,okay,match:evg70947,pct:99/100/.

ccaccATGGCTGATGATCATGCTATAGTCAATAATTCTAACTTACCTCTAGGAAATGCTATTATCAGATCAAGTGTATACTGCCTTTTATTCATTACTGGCTTTTTCGGTAACCTTATCTTTTTAGTAATCTTTGCAAAGTGTCCGTATATGCGCATACCGATTAATTATTTTTTAGTTAATCTGGCCATCGTTGATATTACCGCTACTATCTTTCGTGAACTTGCATTAATTATTGGTTATTTCAACTGTCAGTGGTTATACAATTACCACACTTGTAACTTATTCGGTTTTATGAGAAATATATGTTGTGCTGTTACTATTTTTACACTTTTGGCTATTAGTGTTTGCCGATATTTAGTGATTTGCCAGCATAAAGGGAGAAGAATGACTCGACATGCAGTTTTATGTATTATTGCTTTCGTCTGGTTTTATACCATTTTAGTATGTTTATGTCCGATATTTGGTTGGAATCGATATATTTATCATCAAAATCAGTATGTATGCTCAATAGATTGGAGCTATCAACCTTCTTATATTAGGTATATTATTATTGTCGACGTCTTTTTACCCACGATCATCACCATAATTTGCTATACATTTATTTACATTGTAGTGCATCGACATTCTAGAAGAATACAAGATATTCTACGAATGTCGGGTAGTTTGTCATCCGCCCAATACAATCGAATTGCAAGAGAGAAGAAAGTAACGAAAATTATGTTTGCTGTGTACATAAATTTCTTGATATGCTATCTGCCATTTTCTATAATAACAGCAATACTGATACCGAGTGGAATAAAACCGCCAGCGGAATTATTCTTCGTTGGGACATTTATGATTGATTTAAACAGTTCAATTAATCCTATGCTATATCCCTTCATTTATCGACGCTGTAAAGAAGCTTATTGTGAAGTCTTATGCTGCCATCGATCTCAAACATCGCCAACATCAAATTGTTATTAA

>Tadh164_evg19528 type=CDS; aalen=387,61%,complete; clen=1902; strand=-; offs=1450-287; evgclass=main,okay,match:evg1358279,pct:99/97/.

ccaccATGTCTAATCATAGCCTTAGCTTAAATCAGACTAGGGGATTTACACCGTATACTCTTGCTGAAGCTCAAGTACATTTTACTGCTTATGTGATTGTGATGGTTATAGCTATCCCTGCTAATATAGTCACTATTTTTTCTTTTATCATCGATCGTACATTGTGTACCAATTTTAATCTCTTCTTGCTTAACATGGCTTTCATCGATTTATTGACCAGTTTATTTCGCATCGGTTTCAACGCTTTCTGTATTGCCACTATTAGAAATTCTACATGGCCATATGGTCAGCCACTATGTGATTTTGACGGATTTATACAAGGCGTTGTCTTTGCCGCTAACGTTTACACTCTCATGGTTATAGCAGTTTGCCGTTATATAGCAATTGTTCATTCCAAGCAGCATTTAATTACCAAAAAAATTGTCTATATTATTATTGTTATCATTTGGGTCTTTAGTTCCCTGTGTTCGGTCTTTCCTATCATAGGTTGGAATCGATATCGCTATCAACCTTATGAGCACGCTTGCTTGCCTGATTGGTCTTATGAGAAGAGCTATCCTATTTATATTATGGCTACCGAATTTGTTCTCCCAATGATAATTTTACTTTACTGTTACACCGGAATATTTATCAACCTAAGGAGAAATTCATTCCGAATGCGCACTGTTTCAGAATTGAACAGTGGCAATAATGAGGCCATGAAGAAAATAGCCCAGCGTGAGGGAAAGGTTACTCGTGGCATGTTTATCATCTTCTTAGCCTTCGTCATTTGTTTTGCACCATATAGTGTATGTATGTTTCTATTAGCTTCTGCATTTAATATTCCCATAGACAAGGGTTTGATGTTCTTTTGCGGATTTATGGCCAATGTGAATAGCTTACTGAATCCTATTATTTATGCTATCTTGTATAAACGCTTTCGGCAAGCTTACTATCGGGTGCTGTGTTGTTGCTGCAGTAAATTAACATTCAGCTCTTCATCGGATCATGCAGGATGCAGTAGCAATCACGGATTGCGATCATTAGCAATGGGTGCCATCGAAAATAAGAACGCCAATCATTCTACCAGTCCATCTATGAAAACTTCTTCTATCTTGCCGTTCAGAATTTCTCCTTTTCTGGTTCTTAGTTCAATTTTTCTTGTAATTTTTTTTGGAAAGTTGATTTAA

>Tadh165_evg43354 type=CDS; aalen=309,34%,complete-utrbad; clen=2698; strand=-; offs=969-40; evgclass=main,okay,match:evg936935,pct:99/98/.

ccaccATGGATAATAGAACTAACGAAACTCATCATACACCATTGGAAGTTATTGTTATTAAATCTACTATTTTTATTATTATCGCTATAGCCGCTGTTATTGGCAATAGCTTAGTTATTATTGTTATTTCTCAATATGGACGTAATGCAACCCAGACTGTAACAAACTATCTTATCATGAATTTAGCTATTCTCGACATTATTAATGGATTATTTCAATTACCACTCTATTTAGTCAATATCATAGCTGATAGACAATTTTATGGTGAAATTGGTTGTGCTATTCATGTACAAATTGGAGGTATGGGCTTTATCGGCTCTATTCACAATTTAATGTGGATAGCTATAATACGCTATCTTGTGGTTGTAAAATCTAAAAAATTACAGTCAAGACATGCATACTATTGCATTGGTATTATTTGGGCAACATGTATTTTAGTACCGGCTGTGTCGTTTCTTGGATGGGGTAAAAGTGTTTATGTAGACATTGAGAAAGGCTGCGTTCCCGATTGGAATGGAAATAGCATAAGTGGAATAGCTAGGGGCATCTATGATGCTGTTATCGATTTCGCCATACCTTTAATTGTTATTACTTACTGCTACTTGTTTATTTTTCTCTTTCTTTCAAGGAAACGGAGAGATATGATTTGTGATGGTTTTCAATCACCTACAGCTATGGATAAGGCTAACGTCACTTCAACCATGTTTATCATATTTATTGCATTTCTAATCTGTATTGCGCCTTGGAGTTTATTAGCATTTATTATTATACCAATATCTAAAGTAATAGTACCTGATGGCGTCTACTTTACATCTTCAGCGTTAGTATTCTGTAACTCTGCTATTAATCCAGTCTTATACGCATATCGAAGTAAGGATTTTCGTATAGGATATAAGCAAGCTATTCATAAAATTTGTAGATGCCGTTATAATTGA

>Tadh166_evg70376 type=CDS; aalen=356,79%,complete; clen=1343; strand=-; offs=1186-116; evgclass=maina2,okay,match:evg19501,pct:99/100/.

ccaccATGATCGGAAATCATACATCACATGAAACATCAAATATGGATAGTAGCTATATCTCCAGTGGCCAAATATCCCTATCCGTTAGCATCTATCTTATATTAACAGCTAGCACTGTCGTTGGCAATATCGTCGTCTTATACACCTTTTATCTATATCCCTATCTACGTGTTCCCAGCAATTTATTCATTATTAATTTAGCTATAACAGACATTCTTAACGCTATGGGTCGCAATAATTTCGTATTGATCGGCTTAATAATGGGTCCAGAACCGTATACCCATCTCTTTTGCAATATAAGTGGCTTTATTAACAGTTTAGGCCATGTTGTTATGACAGCAACTCTAGCTGCTACTGCAATATGTCGTTACTTAGTCATTGTTCGCTCATATGCAAGAAAAATTACCACTACAGTTGTCCGTCGTATCATTTTAACAACTTGGTGCTATGCTATCCTCAATTGCTCGATGCCATGGCTTGGCTGGAGCCGTTATACCTATCATCCCTTTGAATTTACTTGTTTACCTGATTGGCACGATGAGAATAATTTTAGTTACATTATATTTGTATTCATCATTGATTCTGTTATTCCATGCATTTTATTAATTGCATGCTACTGGCGTATCTATGCCTTTATATCAGCTAATGCTAATCGAATGCTAATTCATATTAGACAATCATCACTGAAAGCACGTCGTCATCTACCACTGCGTATCATGATAAAGAATGAAATTAAAATCACTAAAGTTTTATTTATTATATGCGTCCTTTTCATGATTTGTATCTTACCTTATAGTATCGTTATTTTCATATTAATACCGTGTAATGTTGATGTTTCCCCCTACGTCGTCTTTTACTGTGGTGTCTTGATCAATTTAAATGGTTCGCTGAATCCTGTTCTTTATGCTGCTATTTATCCTAAGTTGCGTAAAATTTATCGTCGACTAGGTTGTTATTTATGCCACTATGCAACAAAAATACATCAGGAGAGTAGTCGTCGTCGAATTGTCGCCTATCCTCTAAATGACAGTAATCGAATTAAAATTGAAGAGACAAAAGTAACAACCATTGAATAA

>Tadh167_evg70886 type=CDS; aalen=391,33%,complete-utrbad; clen=3482; strand=+; offs=308-1483; evgclass=maina2,okay,match:evg19493,pct:99/100/.

ccaccATGGACAATAAAGCAAAGTATACCTATAATCATAATAACGCTTCGTTCTATGGTTCATCCAACCAAAATGACCATGACGATATAAATCGTGAAGCCTTTGCTTCAGGCATCATATTTTGGAGAGTATTTATTTACTGTGTATTAATAGCAGCGACTGTTATTGGTAACGGTATCGTATTGTTTATATTTTATAAGAGGCCTAGTTTGCGTATTCCTGTTAATTTCTTCATCATTAATTTAGCAATTACCGATATTTTAACAGGCCTAACCAGACAACCAATTATTGTTATCGATTTGTTAAGTAGAGGAGAGTCATTTATTCATACAATCTGCGGCTTAAGTGGGGTAACGCTATGTGTATGTTATGTCGTCACAATCTTTACCTTGCTTGCAACAGCACTATGTCGCTATTTGGTTATTGTACGCTCTTACGGTAGGAAGTTATCGACGAAAGTTGTAGTAATAATTATGTCTTTTATCTGGGTTTATGGAACAATCGATTGCTTATTACCCATCTTTGGTTGGAATCGCTATGTGTACCAACCCGAAGAATACACTTGTTTACCTGATTGGCAAGATTCCAAAGGCGCCAGTTATATCATTTATATTCTTATCATAGATTCTTTAATCCCTTGGACAGTAACTGCAGTATGTTATACCGGTATTTATTTATTTATTAAAAAACATGTTCATCACATGCTTGGACATCTTCAAGCTAACACTCTTACAGTCCATCAACAATTACATTGCCAAGCATTACAAAGGGAAGTCAAAATTACTAAAGTCATGTTTCTTTATTTTCTAACATTTATTCTATGCTATATGCCATACTGCTTGACAATGTTTATTTTAGTTCCAAATAATTTCCACATTTCCCATCATTTGGTTTTTTTTGTTGGTTGCTTAATCAATTTGAATAGTGCAATTAACCCAGTATTATATGCTCTATCGTCAAAGAAATTTAGAAAAGCGTATAACGAGATATTTGGTCGTTTTAGTTTAAGATTATATGTTACTCATGCTATGTCTCGTCGAACAGCCATCATTCAGCCATTCCCTAATCCTTCCGGCCTTGATAAAATAAATGAACTTCAAAATATTGAGCAAAACATTTTTCAAGCTATTTCTGCTACTAGAATATTTCCGCAGCAAACTATATCCGGTACTGAACAATAA

>Tadh168_evg74047 type=CDS; aalen=341,48%,complete-utrpoor; clen=2097; strand=+; offs=236-1261; evgclass=maina2,okay,match:evg910466,pct:99/100/.

ccaccATGACCTGGCCGATAATGAGTAATTCGTGCTATGAAAATATTTCCGATCTATCGCCGACGAAAATCGGATTACGAGCGGGGTATCTAGCGATCATCGGATTCTTTGGAATAGTGGGAAATATATTCTTATTGATTTCATTATTTCTTGATAAGAGAATAAAATCGGCTATTAATTATCTAGTTATTAATCATACCTGCTTAGATCTATTGCAAAGCGGCGTAAGATTACCCTTGTATATAGTCAATATAATCATGAATAGTAATGGCTTAGGTTATGAATTATGTACCTTCCATGGTTTTATTAGCGGTATATATTTTATTGGTACCACTTATACCTTGACGGCGATGGCTATCTTTCGTTATTTGGTTGTGTGCCAATCACCAGCAGTTAATTTAGCTCCTAGGCATGCAATTTATGCCATCGTTCTTGTCTGGATCATTGCTATTATCATGTGCTTACTACCTTTTTGGGGTTTAAGTCATTATGATTATAATATTATTGAAAGATCTTGCTTACCAACCTGGAACGAAGGCTTAGCAAATCAAATTAATGCTGTTTTTGAATCGCTAATTGATTTCGGTTTGCCATTATTGACGACATTTATATGCTATCTGAAGATTTATACCGTCTTGAGAAAAATTAAGATGGATTTAATGCGTTATAGATGTGAATCTATCACAGCAAATACATGCCAGTATTCATCGGATCGAGCTAAAAAACGTTTTACCTTGACATTATCTATCCTATTCCTTAATTTCTTATTGTGCTTTACTGTTTGGAGTATCATTAGTTTTATTGTTTATCCTGTGACTGCTGGCACTGCCTGTGTGCCCGATGATTTGCACTATTTTGCTATATCATTATTATATCTTAATAGTGCATTAAATCCATTGATATTCTTTGCAAGAATGAGATCATTGAGGATGAAGTTTTATTTTATATTGGAAACAATGACTGGACGTAAACAACCTGAAACAACATTGGCAACATCAGGAACAAAACGAAATAATACCATTGAACGATAA

>Tadh169_evg7934 type=CDS; aalen=318,75%,complete; clen=1274; strand=+; offs=254-1210; evgclass=main,okay,match:evg49754,pct:99/100/.

ccaccATGGTTAACGACACTACATCAAATATTACCTCCCAGAGCTATTCTACCGTTACGGTTATATCTGTTCTAAGTTACGTGATTTTAATGATAGTCACCGTTTCAGGAAACGCCTTGGCGCTGAGAGTATTCCATCGATATGTCGAACTTCGAACACCGGTAAACTATTTGTTAATTAATCTAGCCATTGCGGATATATTAAAAGGAGGAATACAAGATGGACTTGTTCTTTGTGGATTATTAACACCTGATTGGGGGTATAATCGGGTTATATGCAATGTTAGTGGATTTATGTTATCAGCTTTCCACGTCGGGACAATTTTCACATTGACTTTGACAGGAATATTCCGTTATCTTATTGTTGTAAACTCCATGAAACATTGGATTACCAGAAAAGTGGTAGTTGCAGCTATTTTAGTCATTTGGTTATATGCTTGTTTAATAGCTTTCTTACCCATTTTGGGATGGAACCACTATACATATGATAAGATGCAACTTTTATGCTTGCCTAACATTACAATGGAAATCTCTTATCCGATATTTGCAATAACGGTGGATATTATGGTACCCATCTTCCTTTTAGGGTATTGTTATTTAAGGGTTTATATGGTCATGAGACAAAACATGCGGCGTACCACAGTCACTTTACATGGAAAAACTCGATCTAGATCGTATTCTCGCTTGCATAAAAGAGAGGTACAGATGACACGAATGATGTTTTCCGTTTACTTAGCATTTGTAATATGTTATATACCTTATGTAATTCAGGCATATGTTCTGATACCACTGCAAATAGATGTACCAAATGTTATAGTCTTTCTTACAGGTTATATTTTAAATACTAATAGTGCTATAAATCCGTTTTTGTATGGATTCACTTCGGAAAAATTTCGTCAAGTTTACAGTGCCTATTTATGTTGTTTATTTTCGTGCCGAAAACGGGTTATGCCTAGTCGCTAA

>Tadh170_evg803040 type=CDS; aalen=353,77%,complete; clen=1364; strand=+; offs=284-1345; evgclass=main,okay,match:evg1753631,pct:100/90/.

ccaccATGGAAAACGAAATTTATGACGGCAGCAACACGTCCAATATTTTATCAGAATCTATCGAATTTACCATTGTTAAAGGAGCAATATTAACGACATTAGGTGTGGTGGCTTTGCTTGGCAATCTGATGGTCGTTTTTTCCATTTTATTACCTAAAAAAATGCGCTCCCCTACTAACTATCTATTATTTAACTTAGCTGTATTAGATATTATCACAGTGACTATACGATTGCCGATCCATTTGATCAATATCATGGAAAATCGTCAAGCAATCGATGATCCTTTATGTCACTTTCATGCATTTCTAACCGGAGTTTGCTTTATTGGCTCAGTTTATTCTATGGTTGGTATTGCTGTCTTTCGTTATATCGTTGTATGTCGATCATTATCTGTCAAACTATCCAATCGACATGCACTCTACGCCATAGGGCTAATATGGTTAATAACAGTATTTATAGCCTTATGGCCATTTTGGGGTTGGAGTAAATTTGTATATGATCCACGTGAAAGAGCTTGTATACCAAGCTATTCAGAAGGCATTAGTGGTCTAATTAATGGAATTTTAGAAATATTTCTTGACTTTGGAATACCTCTAACCACTATATTTGTATGCTACTGGAAGATATACCGCTATATTCGAGCAACTCATCAGCGATTGGCATCCTTCCGTGAATCAGCTAATGTTAAGCATGAATATCGTAAGCGTCGTGTAACTTTAACGATGTGGATTGTATTTGTTGCCTTTTTCATTCTCTATGCCCCTTGGAGTATCATGGCATTTGTTGTTTTTCCAGTCACCAATGGCCAAGCCAATGTACCGGACGGTGTCTACTTTATGGCACAGGCTCTTCTCTATTCTAATTCTGCAGCAAATCCTATTATTTATGGCGTCATGATGGGACAATTTAAAAGGCAATATAAGAAAGTAATCACTTGTGGTTGCTTGACATCCAATGAACAAATGATTCAATCGGATACCGATGCAGAGCGCGTTGTTCCTTATGCATCTCGTGCTATTACTATCGATTGGCTAAAGGCAATGTATATCAATTGTCCAATTTACTAG

>Tadh171_op_evg1145940 type=CDS; aalen=370,57%,complete-utrpoor; clen=1943; strand=+; offs=654-1766; evgclass=maina2,okay,match:evg370964,pct:99/100/.

ccaccATGAGATCCCTAATAACACCACAAAATGAAACTTACAATAAAACAGTACCATCTATTGTAGCTGAAGCTTCGACAATGACATTTTTCGCTATCGCTGCTATTGCAGGAAACATCTTAGTCCTGCTATCAATCAAAAGGAAGAAGAGCTTACAAACTATACGTAATGTGTTTGTTGCTAATCTAGCAGTAGCTGATTTACTCTTTGCCATTATTGGTATGTCATTAGCAGCTGTATCGAGTGTTACTGTTCGGTGGATATTTGGCTACGAAATGTGCCAAATACAAGGATTTGCCAATTCCTTCTTTTGTATTGTATCGCTATTGACATTAAGCGCAGTTAGTATAGATCGTTATTATGTTATACTTCGTCCATTGGATTATCATCGAAAAATGAGCCCTAGAACTGTTACTTGTATGGTTGTGTACATATGGCTTCATGCCTTTGTTTTTGCTGCTTTACCATTTACTGGATGGAGCACTTATCGGTATTACTATGAAGAGTCATTATGTACGGCCGATTGGGGCCATAGCTTAAGCTATACTTTAACAACAATTGGAGCTGCTTTTTTTTTACCTTTAGCTATTATGGCTTATTGTTATTATCATATATTACGAGTTGCTCGGGCACAATCGAAAATAATTGCCGCCGAAATGGCTTCTTTAAATCTTCCGGGAACACCAAGTGAGGACAATAATAATAATAGTAATCAGGAACATAATCAGTCTAATGGCAGTAATTCTTTACCAGTTGATATTAGAGATAAAAGTCGTATAACAAATGCGTTTCGTAAATATCGTCGTGAAGAAAAAGCAACGATTACGTTGATGACTGTCATGGGAACATTTATGATATGTTGGTCTCCCCACTTTTTTGGCATTATTTGCTTAACATTTCCCTACTGTCCATTTCCCAAGTTATTCTTTACAGCTACTACATGGTTAGCACTGATGAATTCAGCCTGCAATCCTGTTATATATGGATTTCTTAATCGAAAATTTAGGCATAGTTTTTATGAGCTATTAGGAGTTAATAAATGCTGCAAAAAGCAAAAAATGATCATTAACGACGATGTAGTTATGGGTGTAACCTACCGCCAAAGTAACGGCGAATAA

>Tadh172_op_evg81053 utrorf type=CDS; aalen=384,18%,complete-utrbad; clen=6213; strand=-; offs=1326-172; evgclass=main,okay,match:evg364583,pct:99/100/.

ccaccATGGCTGATACCTACATTAACAATTTCACGAATAAATCACTAGAGCTATGCAATGGGAGCCTAGTTGTCTCAGATTGTCTATGGCGTATCGTTTTAAGCTGTTTCATGACCCTATTTATGTTGATGAGCTGCATTGGTAATGGCGCCGTCTTACTCGTTCTACGCTACCATCATGATGATATCAAGTCGGCATCTAACTATTTTATCACTAATTTAGCCTTAACTGATTTTTTACTGGGCGTACTATGCATGCCCTGTATTTTGATTTCCTGCTTAAATGGGCAATGGGTTTTTGGTCAGAACTTATGCAGTTTAACAGGGTTTGCTAACTCATTTTTTTGTATTAATTCCATGATTACTTTAGCCGCTGTTAGTGTGGAAAAATACTGTGCTATTGCTTCACCATTGACATATCATCATTATATGAGCAAAAGTAAAGTCACATGTGTAATTTCAATTATATGGATCCATTCAGCTATTAATGCTAGTCTACCCTTTTTGGGCTGGGGAGAATATGTCTACCTTCCTTTCGAAACAATTTGCACAGTTGCTTGGTGGAGCTTTCCAAATTATGTTGGTTTTATAGTTGGTATTAATTTTGGACTACCTACCGTGATCATGAGTTGTACTTATTTCCTCATACTAAAAATTGCTCGTAAACATTCAAGGCGGATAGGTGTATCTACGTCAACTGTAGCAATTTCAACTTATCTAAGCCCAACTGGTACATATAATAACCTTAGTCCAGTTAACGTAACTACAAGTGATCATCAAGAAACTCTGCCATTACCTTGTGATACTGATTTAGGTATTCTAACTAGCAATTCTATCTCGCAACGTATCCGATTACAGTACGTTCGTGAAATGCTAACTAGCCGAAGAATACATTATAAAACACATATTAAAGCAACATTGATGTTATTAATTGTCATCGGTAGTTTTATAGTCTGTTGGCTACCGCATCTTATTAGTATGGTATATTTAACCATTTATGAAATAAGCCCGTTACCCTGTAGTTTTCATCAAATTACAACATGGCTAGCAATGGCTAACTCGGCTTTTAACCCAATCATATATGGAGCTATGGATACATCTATAAGAAAAGGTCTTAAAACCTTACTCGGATCCTGGGTAAAATATTGTAAATTATACTAA

>Tadh173_op_evg17446 type=CDS; aalen=340,22%,complete-utrbad; clen=4518; strand=+; offs=1385-2407; evgclass=maina2,okay,match:evg372020,pct:99/100/.

ccaccATGGATAATAACAATAGCAATACGACTGTACCCAAAATGGATCTGGTTTATCTTATTTTACAAGCCACTATATTATCCATTATTACCCTATTTATGATTGTAGGCAATTTTATCGTTATTGTAGTTGTTAATCGTTCAGAACAGCTACAAAATGCTACCGGTATCTTTATGGCCAACTTAGCCGTTACCGATTTAACTTTAGGTGTTATTTTAATGCCCATTACCATTGCTAGCTCCATCTTAGGCCGTTGGATATTTAGTGATATAATGTGTAAATTTTGCGGTTTTTTAAATGTTATGTTATGCAGCACATCAGCATTAACTGTCATGTTATTGAGTATCGACCGTTTTATTGCTATTTCACGTCCAATGCAATATATTAAAATTATGTCCAAAAAGCGAGCCTTAGTGTTATCCACTTACATGTGGATCCATTCAGCTATCGTCTCGACATTTCCATTACTAGGTTGGGCTAAATATGAATATGTTGAAGCCGAAGCAGTTTGTTTTGCTATCGAAAGTGTATCCTATTTTAATTTTCTATGCGCATCGACAGTATTTTTGGCTATCTTTTTAATTATCATTACCCATATCTACATCTTAAAAGTGGCTATCAAGCAATCACAGCAAATTGTAACCTTAACACCTGGTTTTGAAGATGTCACAAGAGAAACGATGCGCCAAATGAATCGTAAAACAGCCAAGACTGTCTTAATTATTGTTGGTGTCTATTTAGCCTGTTGGATACCTTATATTTCATTAACTTATGTCCAAATTTACGCTAACTATGTTCCACCACCAATTGCTATTACAATCACATCAGGTCTAATTTTTGTCAATTCAGCATCCAATCCAATTATTTATGGTACATTCCATCGACGTTTTCGTAATGCTTTCATTCGGTCATTTATGCCAATTTTGACCAAATGTAGCATAGTTGATGAATACGCTCATAATCGCGAGCAAGCAAGTTATCTTCATACTTCCAATACTGTACGGAAACTTAATAATTCCAATCATTAA
